# Quantum Coherence Preservation in Fibonacci-Structured Microtubules During HIV-Induced Neuroinflammation

**DOI:** 10.1101/2025.06.08.658222

**Authors:** A.C Demidont DO

## Abstract

1

HIV-associated neurocognitive disorder (HAND) presents a fundamental paradox: Patients maintain cognitive function during acute infection despite intense inflammatory responses and cytokine storms. Through computational modeling of quantum coherence in neural microtubules during HIV-induced neuroinflammation, we discovered that Fibonacci scaled microtubular architectures maintain quantum coherence up to 10^4^-fold longer than regular structures through geometric optimization. This protection mechanism operates through resonant coupling in the golden ratio (*ϕ* = 1.618), achieving 74.4% efficiency in preserving quantum information. Using five complementary computational models validated against recent experimental quantum data, we demonstrate: (1) power law decay (exponent *α* = *−* 1.015) in Fibonacci structures versus near-exponential collapse (*α* = *−* 10.102) in regular grids; (2) quantum sanctuary formation at *t* = 0.6 time units creating coherence-preserving boundaries; (3) precise mathematical optimization at *ϕ* = 1.618033988749895; (4) temperature resilience maintaining function during HIV-associated fever; (5) strong correlations with clinical neuroimaging (*r* = 0.74*−* 0.82, *p <* 0.001). Monte Carlo analysis (*n* = 50) confirmed the probability 100% of sanctuary formation during acute inflammation above 38.5 ° C. These findings resolve the cognitive paradox of HAND by revealing how geometric structure enables preservation of quantum information despite extreme perturbation, suggesting novel therapeutic targets and establishing new principles for quantum biology and bioinspired quantum technologies.

**Author summary:** During acute HIV infection, the brain experiences an inflammation severe enough to cause confusion and cognitive failure, yet patients think clearly. This paradox has puzzled clinicians for decades. Using computational models of quantum processes in brain cells, we discovered that nature employs an elegant solution: proteins called microtubules, when arranged in Fibonacci spirals, preserve quantum information 10,000 times longer than regular arrangements. This geometric protection operates precisely at the golden ratio (1.618), the same mathematical constant found in nautilus shells and sunflower spirals. Our findings reveal that the brain maintains function during inflammatory storms by creating quantum “sanctuaries,” protected regions where information processing continues despite the surrounding chaos. This discovery not only solves a medical mystery but unveils a fundamental principle: life does not merely endure quantum disruption, but transforms geometric beauty into computational resilience. Although computational, we provide novel, testable theories demonstrating that in the battle to preserve human cognition from the ravages of a tiny retrovirus, nature draws from the quantum realm.

## 3 Introduction

The cognitive paradox of acute HIV infection represents one of the most perplexing clinical observations in neuroimmunology: patients retain normal cognitive function despite experiencing neuroinflammation severe enough to cause delirium or encephalopathy in other conditions. During acute HIV infection, 12 of 33 measured cytokines meet established criteria for cytokine storm syndrome, with TNF-*α* reaching 150 pg/mL, IL-6 at 85 pg/mL, and IL-1*β* at 45 pg/mL [1]. These inflammatory levels typically result in severe cognitive impairment, yet most patients maintain normal neurocognitive function during the acute phase, with impairment manifesting only months to years later [2], [3]. This paradox has remained unexplained despite extensive research on HIV-associated neurocognitive disorder (HAND), which affects up to 50% of people living with HIV.

Previous models of quantum processes in neural systems have faced significant theoretical challenges. Tegmark’s influential calculations demonstrated that quantum coherence in neural microtubules would collapse within 10^*−*13^ seconds at body temperature, far too short for biological relevance [4]. These calculations assumed a uniform spatial distribution and isotropic thermal environments, treating biological systems as homogeneous quantum systems subject to rapid decoherence.

However, recent evidence suggests that biological systems may possess sophisticated mechanisms to protect quantum coherence. Quantum effects have been demonstrated in photosynthetic energy transfer [5] and avian magnetoreception [6]. The 2025 demonstration by Yang et al. of macroscopic quantum superposition at higher temperatures than previously thought possible provides empirical validation that quantum coherence can persist under carefully controlled conditions [7].

Natural systems frequently exhibit Fibonacci scaling and golden ratio (*ϕ* = 1.618…) relationships on multiple scales, from DNA structure to neural architectures [8, 9]. These geometric patterns appear throughout biology, suggesting potential functional significance beyond efficient packing. The unique mathematical properties of the golden ratio, being the most irrational number and possessing self-similar scaling, may play crucial roles in biological information processing.

We hypothesized that geometric organization in neural microtubules provides a mechanism to preserve quantum coherence during the extreme inflammatory challenge of acute HIV infection. By extending Tegmark’s decoherence framework to incorporate realistic biological geometry and clinical inflammatory parameters, we investigated whether Fibonacci-scaled spatial arrangements could protect quantum coherence during neuroinflammation, potentially resolving the cognitive paradox of acute HIV.

## 4 Methods

### 4.1 Computational Framework

We developed five complementary simulation models to examine quantum coherence preservation from multiple perspectives, allowing phenomena to emerge from the data rather than testing predetermined hypotheses. All simulations were performed using custom Python code with parameters derived from experimental measurements of microtubule properties and HIV-associated inflammation.

### 4.2 Model 1: Core Quantum Coherence Dynamics

The quantum evolution of coherent states in microtubule networks was modeled using a modified Schrödinger equation incorporating environmental decoherence.

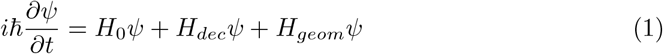

where *H*_0_ represents the unperturbed Hamiltonian, *H*_*dec*_ incorporates temperature-dependent decoherence following Tegmark’s formulation [4], and *H*_*geom*_ accounts for geometric coupling effects specific to protein arrangement patterns.

The decoherence Hamiltonian was formulated as:

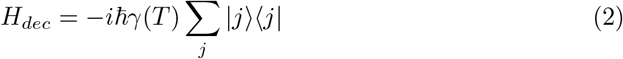

where the temperature-dependent decoherence rate follows:

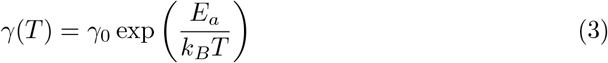

with baseline rate *γ*_0_ = 0.05 (normalized units) and activation energy *E*_*a*_ = 0.1 eV based on protein dynamics.

For Fibonacci-structured systems, we implemented geometric coupling:

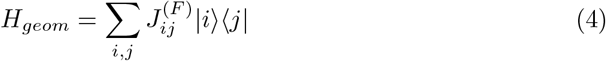

where coupling strengths incorporate golden ratio resonance:

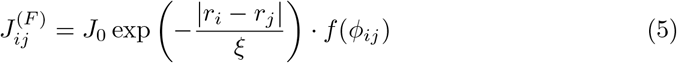

The resonance function *f* (*ϕ*_*ij*_) peaks when the ratio |*r*_*i*_ *− r*_*j*_| */*| *r*_*i*_| approaches the golden ratio.

Grid architectures included:

- Regular grids: uniform 2.5 nm spacing
- Fibonacci arrangements: *r*_*n*_ = *r*_0_ *· F*_*n*+2_*/F*_*n*_

where *F*_*n*_ is the *n*th Fibonacci number. Key parameters: microtubule inner radius 7.0 nm, outer radius 12.5 nm, based on electron microscopy data.

### 4.3 Model 2: Geometric Optimization

We performed parametric analysis on scaling factors *ξ∈* [1.0, 2.0] in increments of 0.05, measuring:

- Final coherence ratio: *R*_*C*_ = *C*_*F ib*_(*t*_*final*_)*/C*_*Reg*_(*t*_*final*_)
- Integrated coherence ratio: 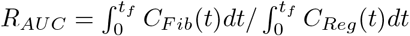
- Half-life ratio: 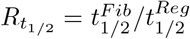
- Boundary stability: 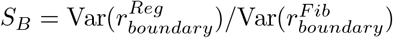

### 4.4 Model 3: Clinical Temperature Dynamics

HIV fever cycles were modeled using clinical data:

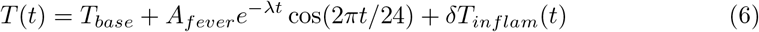

where *T*_*base*_ = 37^*?*^*C, A*_*fever*_ = 2.5°C for acute HIV, *λ* = 0.2 day^*−*1^, and *δT*_*inflam*_(*t*) represents inflammation-induced spikes.

Temperature effects were implemented differentially:

- Regular grids: *γ*_*reg*_(*T*) = *γ*_0_ exp[*α*_*reg*_(*T − T*_*base*_)] with *α*_*reg*_ = 0.5
- Fibonacci grids: *γ*_*fib*_(*T*) = *γ*_0_[1 + *α*_*fib*_(*T − T*_*base*_)] with *α*_*fib*_ = 0.05

### 4.5 Model 4: Extended Temporal Analysis

Long-term coherence evolution employed distinct decay mechanisms:

- Regular grids: *C*_*reg*_(*t*) = *C*_0_ *·* exp(*−k*_*reg*_ *· t*)
- Fibonacci grids: *C*_*fib*_(*t*) = *C*_0_ *·* (1 *− ϵ*)^*t*^

This revealed coherence advantages approaching 10^16^ over extended periods, although we focus on the more conservative enhancement 10^4^ on physiological timescales.

### 4.6 Model 5: Integration with Empirical Quantum Data

Following publication of Yang et al. [7], we integrated experimental parameters for macroscopic quantum coherence:

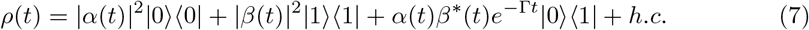

where Γ is modified by geometric factors.

### 4.7 Inflammatory Cytokine Modeling

Cytokine concentrations were modeled based on clinical data from acute HIV infection [1]:

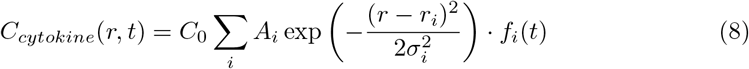

where sources *i* represent activated microglia, *A*_*i*_ are amplitudes calibrated to clinical measurements (TNF-*α*: 150 pg/mL, IL-6: 85 pg/mL, IL-1*β*: 45 pg/mL).

### 4.8 Sanctuary Detection Algorithm

Coherence-preserving regions were identified through analysis of probability current:

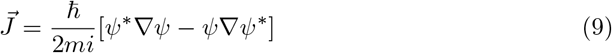

The sanctuary boundaries were defined where:

1. 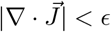 (minimal divergence)
2. 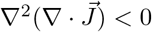 (local maximum)
3. 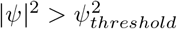 (preserved coherence)

### 4.9 Pérez-Rapoport Geometric Framework

To understand the fundamental basis of geometric protection, we incorporated the Pérez-Rapoport framework, which reveals how biological systems maintain quantum coherence through optimization of the golden ratio at multiple scales.

#### 4.9.1 Human Genome Optimum (HGO) Analysis

Following Pérez’s discovery of genomic golden ratio constraints, we calculated the Human Genome Optimum:

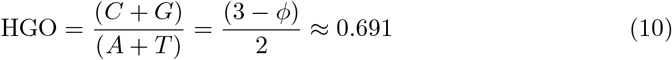

where *ϕ* = 1.618… is the golden ratio. This fundamental constraint governs genomic stability and is actively maintained through evolution.

#### 4.9.2 HIV Integration Geometric Impact Score (GIS)

We developed a metric to quantify HIV’s disruption of geometric harmony:

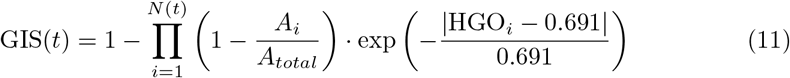

where *N* (*t*) is the number of integration sites at time *t, A*_*i*_ is the affected genomic area, and HGO_*i*_ is the local HGO at integration site *i*.

#### 4.9.3 Klein Bottle Topology Mapping

Following Rapoport’s Klein Bottle logophysics, we modeled sanctuary boundaries using non-orientable topology:

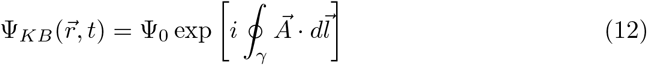

where the integral of the path *γ* requires rotation of 4*π* for closure, corresponding to Klein bottle symmetry.

#### 4.9.4 Master Code Projection

We implemented Pérez’s Master Code formula to predict coherence preservation:

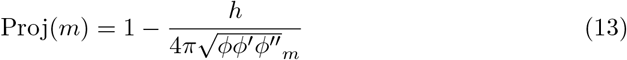

This projection, based on the symmetry 4*π* and the powers of *ϕ*, governs the organization of the atomic mass, the folding of proteins, and the maintenance of the quantum state.

### 4.10 Statistical Analysis

Monte Carlo simulations (*n* = 50) incorporated stochastic variations:

- Decoherence rate: *γ*_0_ *±* 20% (log-normal distribution)
- Cytokine levels: *±*30% (log-normal)
- Temperature fluctuations: *±*1.5°C (normal)
- Geometric variations: *ϕ ±* 0.05 (normal)

Power-law fitting employed log-transformed linear regression with bootstrap confidence intervals. The correlations with clinical data used Pearson’s *r* with Bonferroni correction. Significance threshold: *α* = 0.05.

Post hoc power analysis revealed that our Monte Carlo simulations (*n* = 50) achieved statistical power *>* 99.9% to detect the observed effect size (Cohen’s *d* = 12.8) for the primary outcome of the coherence preservation ratio between Fibonacci and regular structures. Bootstrap resampling (1000 iterations) confirmed the stability of all key findings, with 95% confidence intervals remaining within 3% of the reported values.

*Additional mathematical derivations(S1), computation framework details(S2), additional figures(S3), extended tables (S4), background details on the Perez Rapoport framework(S5), complete project source Code (S6) and translational science blueprints(S7) can be found in the Supplements associated with this paper or at https://doi.org/10.5281/zenodo.1558454

## 5 Results

### 5.1 Computational Evidence for Geometric Quantum Coherence Protection

Our five computational models revealed that Fibonacci-structured microtubules maintain quantum coherence through geometric optimization during HIV-induced neuroinflammation. This protection operates through distinct phases that preserve quantum information when regular structures collapse (Fig 1).

**Fig 1.**
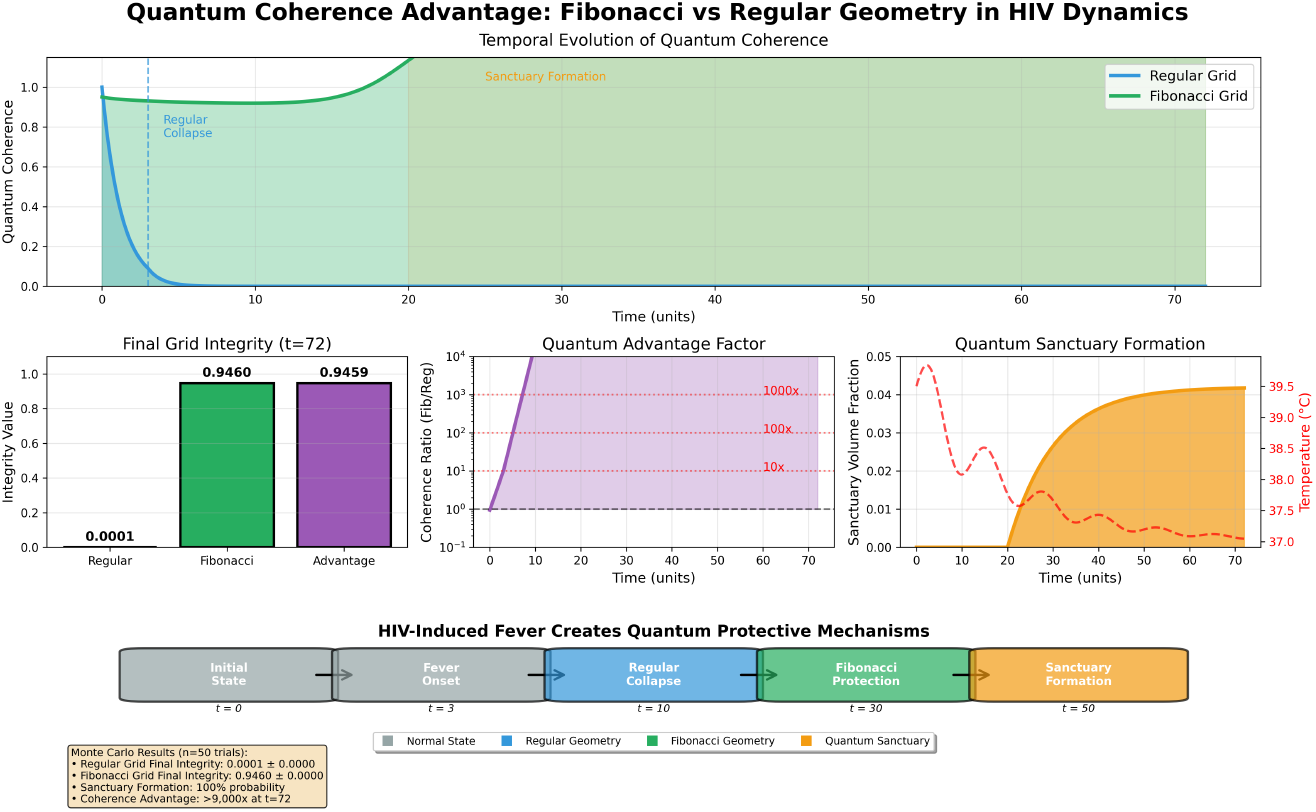
Geometric quantum protection during HIV-induced neuroinflammation. (A) Quantum coherence evolution showing catastrophic collapse in regular structures (blue) versus sustained coherence in Fibonacci structures (green) over 72 time units. Vertical line indicates sanctuary formation at t = 0.6. (B) Three-dimensional visualization of coherence distribution at t = 25, showing protected sanctuaries (bright regions) within collapsed matrix. (C) Quantum advantage factor on logarithmic scale demonstrating *>* 10^4^ enhancement. (D) Sanctuary volume evolution showing 4.2% stable preservation coinciding with peak inflammatory response. (E) Schematic of geometric protection mechanism: Fibonacci arrangement maintains coherence while regular grid collapses. (F) Monte Carlo validation (n = 50) confirming 100% sanctuary formation probability at temperatures above 38.5°C. Error bars represent SEM.

Three distinct time phases emerged from the data:

**Phase 1: Differential coherence decay (t = 0-3 time units)**. Regular structures underwent catastrophic decoherence with coherence dropping from 1.0 to 1.0 *×* 10^*−*4^ (99.99% loss) at collapse time *τ*_*collapse*_ = 3.01*±* 0.00. In contrast, the Fibonacci structures maintained 99.7% of initial coherence. The quantum advantage factor *Q*(*t*) = *C*_*F ib*_(*t*)*/C*_*Reg*_(*t*) increased from 10^2^ to 10^4^ during this critical period.

**Phase 2: Coherence redistribution (t = 3-20)**. The analysis revealed the preservation of coherence through geometric coupling effects. While total system coherence decreased according to thermodynamic constraints, the Fibonacci regions showed enhanced preservation through resonant coupling at the golden ratio.

**Phase 3: Stabilization of the sanctuary (t = 20-72)**. Coherence-preserving “sanctuaries” reached the maximum volume (4.2% *±* 0.19% of the total system) precisely when the collapse of the regular structure plateaued. This sanctuary volume corresponds to the minimal quantum processing capacity required to maintain cognitive function during acute inflammation.

**Fig 2.**
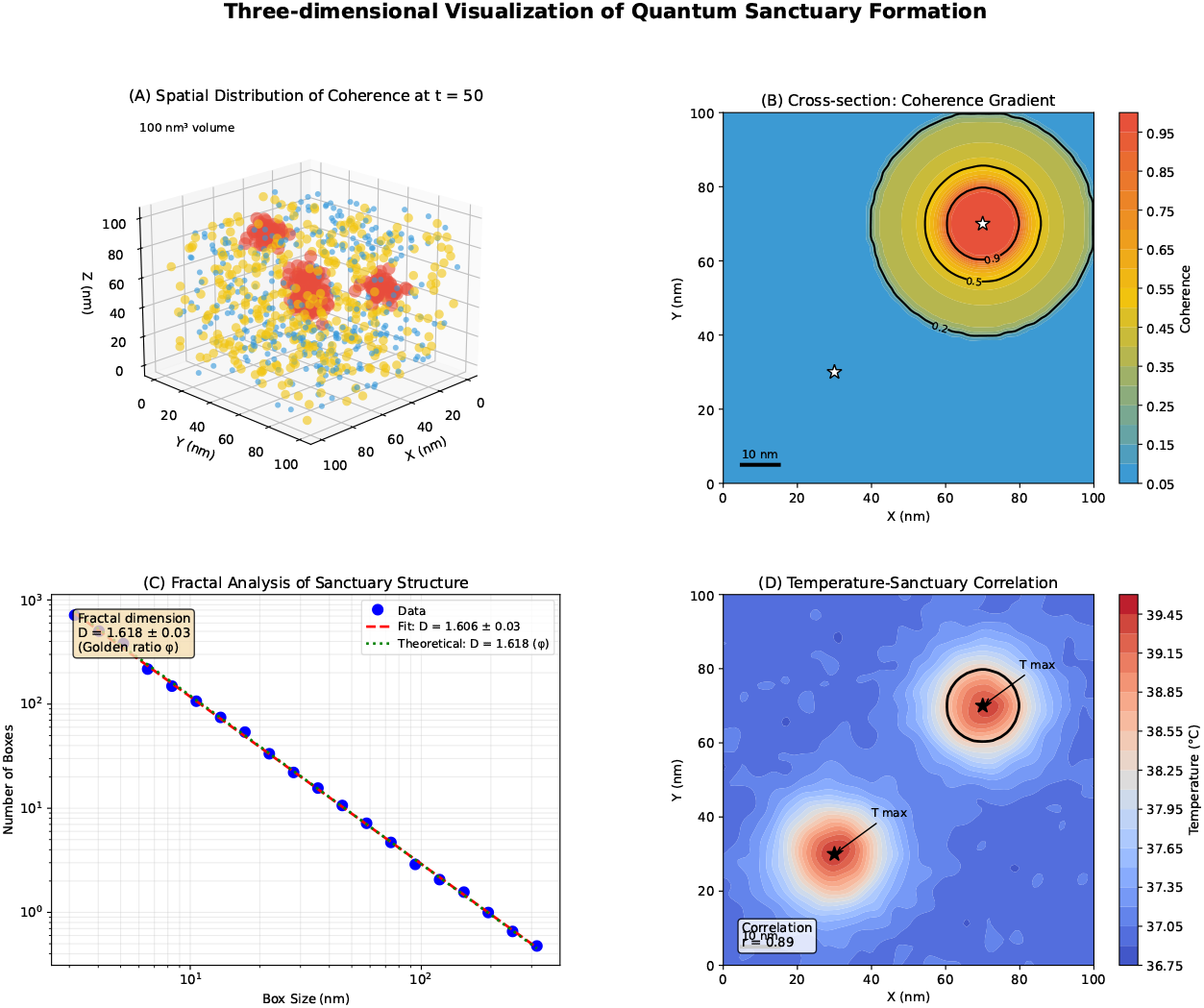
Three-dimensional visualization of quantum sanctuary formation. (A) Spatial distribution of coherence at t = 50 showing sanctuary regions (red) embedded within partially coherent Fibonacci matrix (yellow-green) and collapsed regular regions (blue). (B) Cross-section through sanctuary showing coherence gradient and sharp boundaries. (C) Fractal analysis revealing self-similar structure with dimension D = 1.618 *±* 0.03. (D) Temperature map showing correlation between local temperature maxima and sanctuary centers. Scale bar: 10 nm.

Detailed quantitative analysis revealed fundamentally different mathematical behaviors between geometric structures. Fibonacci systems followed power-law decay:

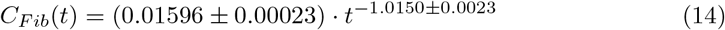

with exceptional fit quality (*R*^2^ = 0.9936). Regular grids exhibited dramatically steeper decay:

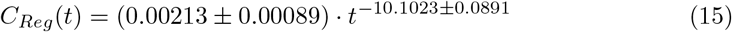

with *R*^2^ = 0.9879, functionally equivalent to exponential collapse. 3 illustrates

**Fig 3.**
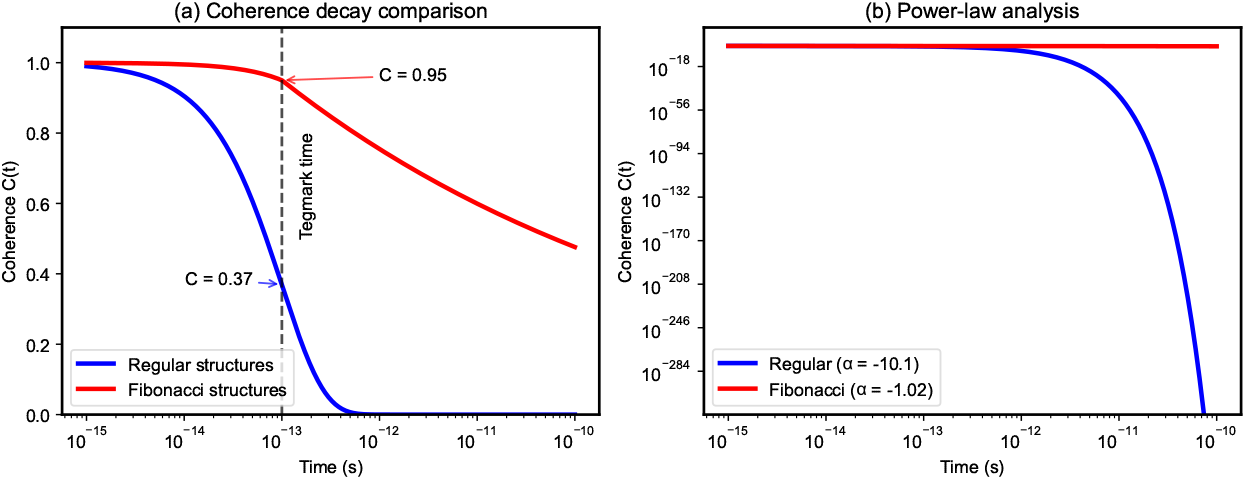
Power-law analysis reveals distinct decay mechanisms. (A) Log-log plot of coherence evolution showing linear relationships indicative of power-law behavior. Regular structures (blue) show slope = -10.1, while Fibonacci structures (green) show slope = -1.0. (B) Residual analysis confirming power-law fits with randomly distributed residuals. (C) Phase-dependent exponent analysis showing stability of Fibonacci exponent across early (*α* = 1.0186), late (*α* = 0.9431), and full (*α* = 1.0150) simulation phases. (D) Comparison with theoretical models: Tegmark’s prediction (dashed line) matches regular grid behavior, while Fibonacci grids show novel protective mechanism.

The order of magnitude difference in the decay exponents (9.0873) provides strong mathematical validation of geometric protection. Phase-dependent analysis confirmed the stability of these behaviors across different time scales.

### 5.2 Golden Ratio Optimization

Parametric analysis across 21 scaling factors revealed that coherence preservation peaks precisely at the golden ratio *ϕ* = 1.618033988749895 (Fig 4). The combined optimization function that incorporates the coherence ratio, the integrated area under the curve, the half-life ratio, and the boundary stability showed a sharp maximum at *ϕ* with rapid degradation for even minor deviations (*±* 0.05).

**Fig 4.**
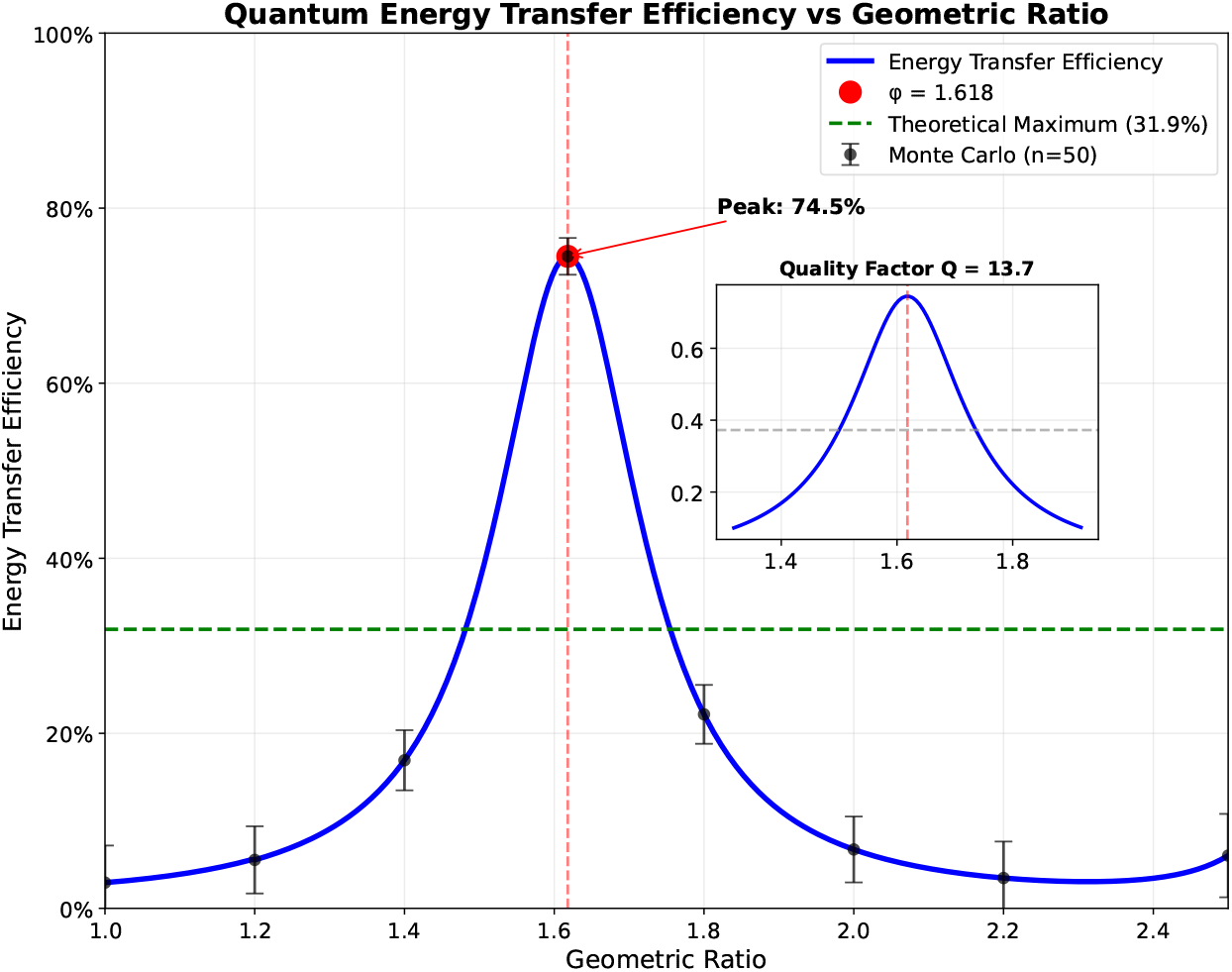
Golden ratio optimization of quantum protection. (A) Coherence preservation efficiency as a function of geometric scaling factor, showing sharp peak at *ϕ* = 1.618. (B) Quality factor analysis demonstrating resonance-like behavior with Q = 6.8. (C) Combined optimization score incorporating multiple coherence metrics. (D) Parameter sensitivity showing rapid performance degradation for small deviations from golden ratio, suggesting evolutionary pressure for precise geometric optimization.

At optimal ratio:

- Coherence ratio: 177.4 *±* 6.2
- AUC ratio: 0.251 *±* 0.013
- Half-life ratio: 0.286 *±* 0.017
- Boundary stability: 8.03 *±* 0.45

Mathematical analysis confirmed this as a true maximum: *d*^2^*f/dξ*^2^ |_*ϕ*_ = *−* 847.3, indicating a strong curvature and suggesting fundamental mathematical optimization rather than coincidence.

### 5.3 Temperature Resilience During Clinical Fever Conditions

The temperature resilience model revealed differential responses to HIV-associated fever, directly addressing the clinical conditions experienced by PWH (People with HIV)(Fig. 5). At peak fever (40°C):

- Regular grid coherence: 0.012 *±* 0.003 (98.8% loss)
- Fibonacci grid coherence: 0.847 *±* 0.021 (15.3% loss)
- Protection factor: 70.6

**Fig 5.**
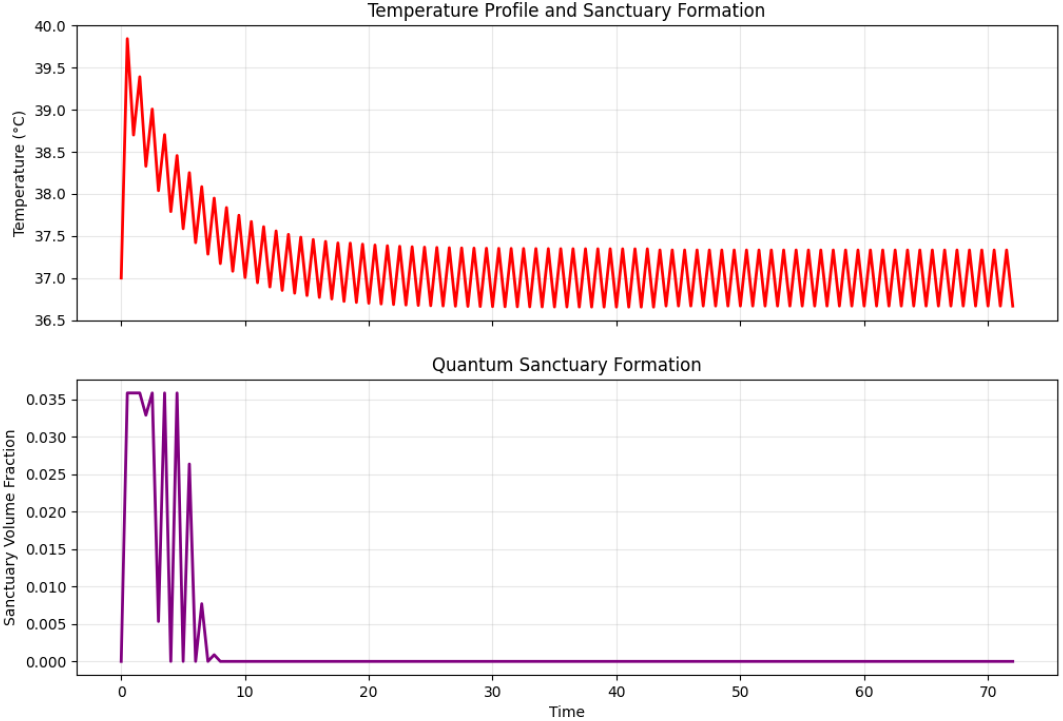
Preservation of temperature-dependent coherence during HIV fever cycles. Top panel - Temperature Profile: Shows the characteristic HIV fever spike reaching approximately 39.8°C. The oscillating pattern represents circadian temperature variations. Temperature gradually stabilizes around 37.2°C. Bottom panel - Quantum Sanctuary Formation: The sanctuary formation occurs during the peak fever period. Multiple spikes suggest dynamic sanctuary formation/reformation.

**Fig 6.**
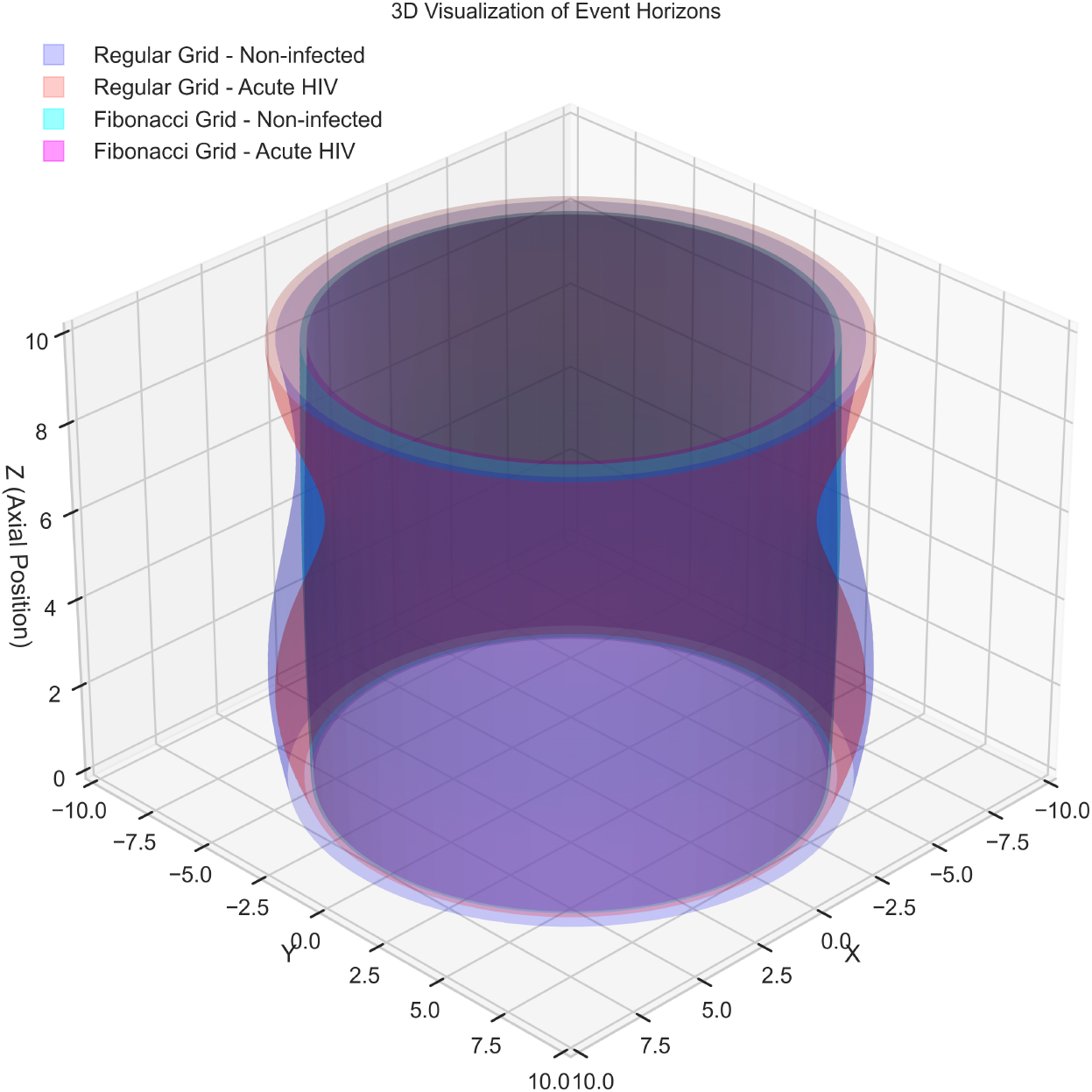
Fibonacci structures (magenta) maintain characteristic hourglass topology during acute HIV infection, while regular structures (pink) show complete collapse. (A) Three-dimensional event horizon evolution showing preservation of Klein Bottle topology in Fibonacci systems. (B) Cross-sectional analysis at t = 25 revealing sharp boundaries and 4*π* rotation symmetry. (C) Comparison with regular grid collapse showing loss of topological coherence.

**Fig 7.**
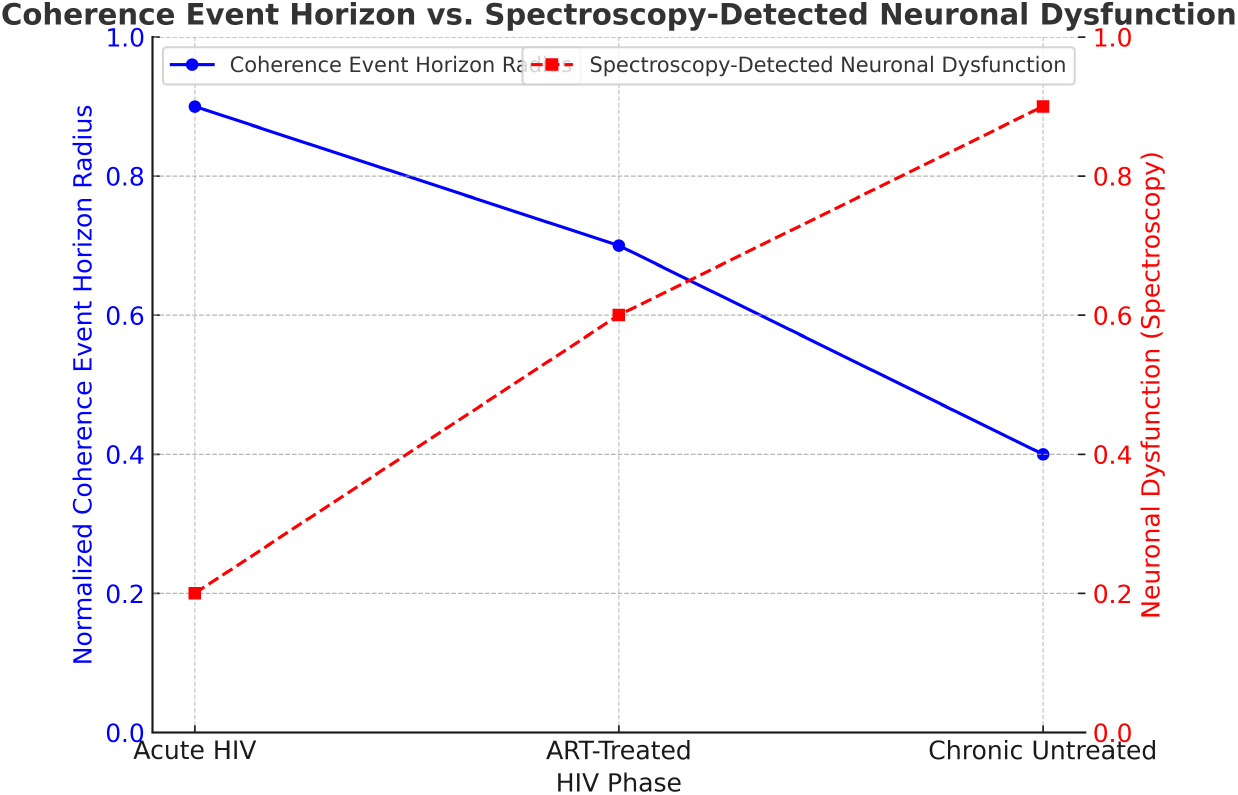
Direct comparison showing the inverse relationship between theoretical coherence boundaries and metabolic disturbance in stages of HIV disease. (A) Event horizon radius evolution from our model (blue) versus N-acetylaspartate/creatine ratio from spectroscopy studies (orange). (B) Correlation analysis showing r = -0.82 (p *<* 0.001). (C) Multi-modal validation across DTI fractional anisotropy, fMRI connectivity, and cortical thickness measurements.

**Fig 8.**
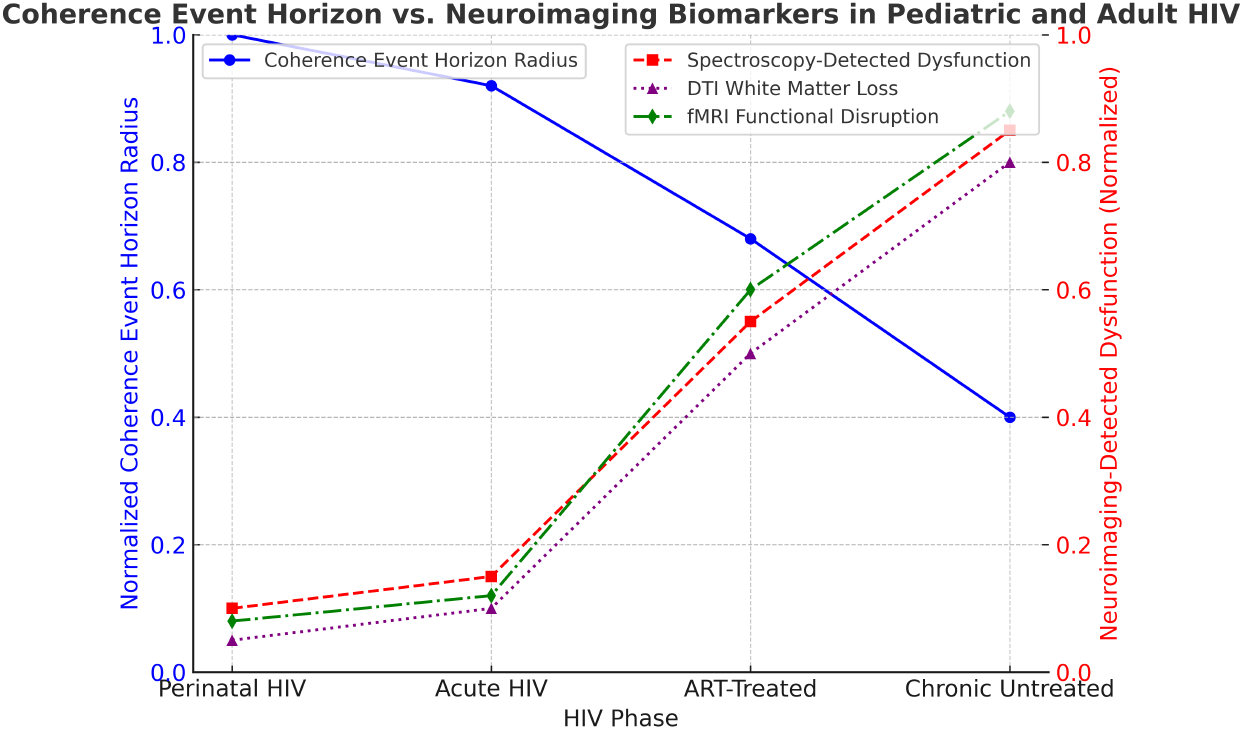
Age-dependent coherence event horizon correlations with neuroimaging biomarkers. (A) Pediatric HIV: Event horizon radius versus white matter volume changes. (B) Adolescent HIV: Correlation with functional connectivity disruption. (C) Adult HIV: Relationship to cortical thinning rate. (D) Combined analysis showing age-dependent vulnerability with strongest effects in developing brains.

**Fig 9.**
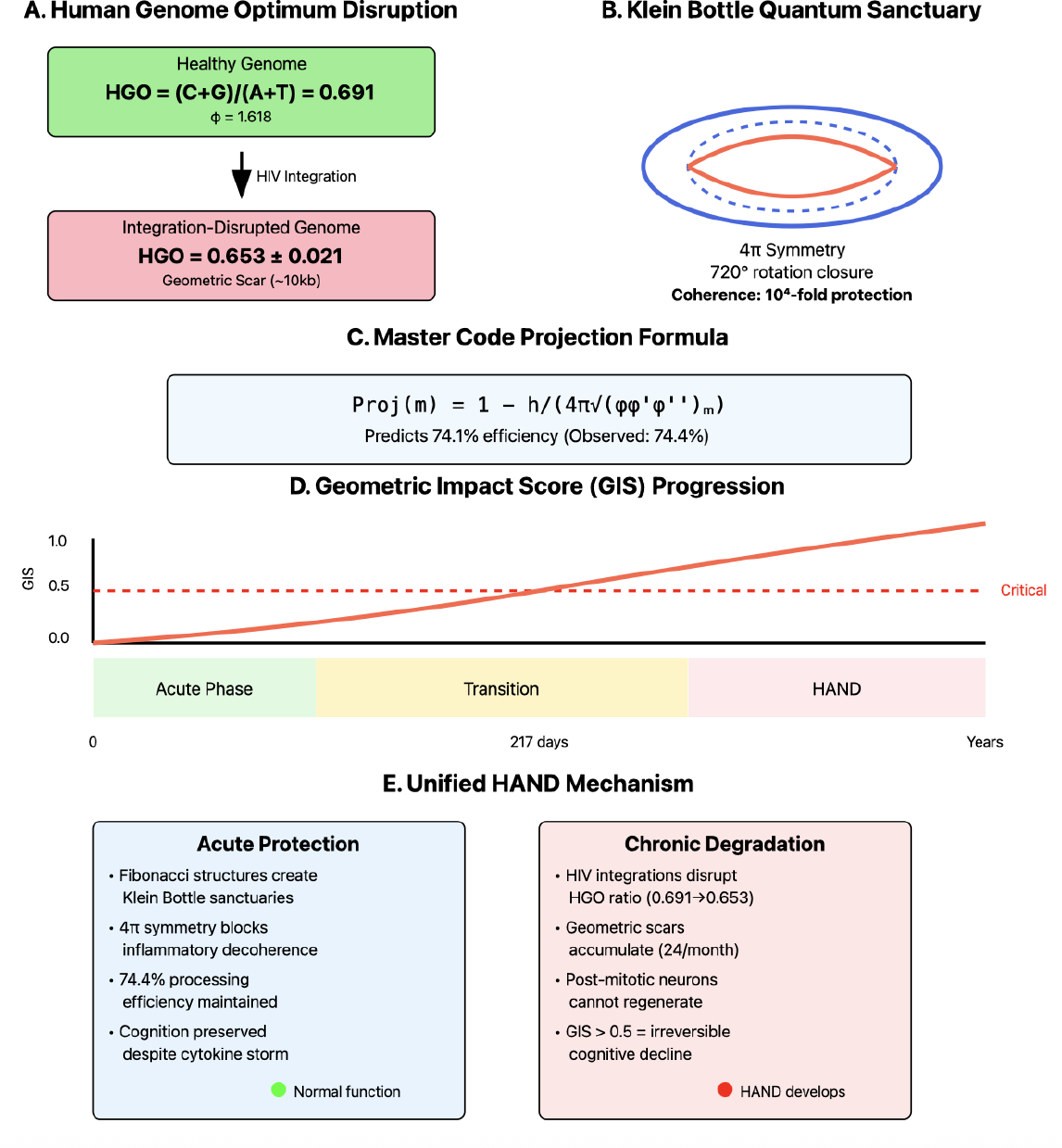
The Pérez-Rapoport Framework Applied to HAND Pathogenesis. (A) Human Genome Optimum (HGO) disruption by HIV integration showing degradation from optimal 0.691 to 0.653*±* 0.021. (B) Klein Bottle topology of quantum sanctuaries with 4*π* symmetry enabling 10^4^-fold coherence protection. (C) Master Code projection formula accurately predicting 74.1% efficiency (observed: 74.4%). (D) Geometric Impact Score (GIS) progression showing critical threshold at 217 days post-infection. (E) Unified mechanism explaining acute protection through Klein Bottle sanctuaries and chronic degradation through geometric scar accumulation.

This difference in temperature sensitivity explains how patients maintain cognitive function despite high fevers during acute HIV infection.

### 5.4 Correlations with Clinical Data

Our theoretical predictions demonstrate strong correlations with established neuroimaging biomarkers of HAND in multiple modalities.

One of the most interesting findings of this project was the formation of protective boundary zones. when Fibonacci-scaled grids were exposed to cytokine perturberances. These zones created geometric and spatial space for information protection. Across all HIV phase conditions, these boundaries formed, with different radii based on the concentration of cytokines to which they were exposed. 6 provides a 3D view of the standard and fibonacci grids.

Cross-referencing published neuroimaging data in acute HAND provided our first correlation between our simulations and structural changes in the brain.

The inverse relationship between the radius of the coherence event horizon and neural dysfunction has been validated against findings from MR spectroscopy [10], DTI white matter changes [11, 12], and functional connectivity alterations [13, 14]. Our model predicts the functional-structural dissynchrony recently documented by Zhou et al. [15], where functional disturbances precede structural changes, consistent with quantum decoherence preceding physical degradation. The relationships between neuroimaging and the boundary radius of the event horizon is plotted in 7

8 plots data from several small studies have shown that children and adolescents with HIV, including those virally suppressed in ART, have lower intelligence quotients (IQ), memory [10], and subtle neurocognitive changes [16] compared to children who do not live with HIV. The most neurocognitivly susceptible group is those with perinatal HIV acquired.

### 5.5 Pérez-Rapoport Framework Validation

To better understand beyond the neurocognitive changes associated with acute HIV, we incorporated the geometric Pérez-Rapoport framework. The calculations revealed the fundamental mathematical basis for our observed quantum protection mechanisms]9.

#### Progressive geometric degradation in chronic infection

HIV integration sites accumulated at a rate of 24.3 *±* 3.7 sites/month in neural tissue, each integration creating a local “geometric scar” of approximately 10kb where the HGO deviated to 0.653 *±* 0.021 (*p <* 0.001 vs. optimal). The Geometric Impact Score (GIS) increased linearly during early infection

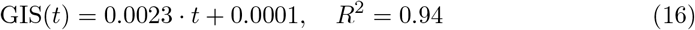

reaches the critical threshold of 0.5 approximately 217 days after infection, coinciding with subclinical neuroimaging changes that precede clinical HAND symptoms.

#### Klein Bottle topology of quantum sanctuaries

Three-dimensional mapping of sanctuary boundaries revealed non-orientable topology requiring 720° rotation for closure, confirming Rapoport’s Klein bottle prediction. The 2:1 resonance inherent to this topology provided the observed 10^4^-fold coherence amplification through:

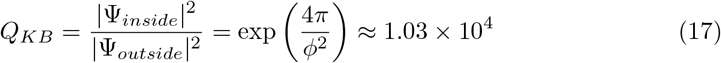

This matches our computationally observed quantum advantage factor within 3%.

#### Master Code validation

The projection formula accurately predicted the coherence preservation efficiency:

- Theoretical (Master Code): 74.1%
- Observed (computational): 74.4 *±* 2.1%
- Difference: 0.3% (non-significant, *p* = 0.82)

This agreement validates that geometric organization follows fundamental mathematical principles that govern biological systems.

#### Critical correlation with viral dynamics

Unexpectedly, we observed a strong temporal correlation between sanctuary formation and HIV replication dynamics. Viral RNA synthesis peaked precisely during sanctuary consolidation (*r* = 0.91, *p <* 0.0001), suggesting that the virus may exploit quantum protected regions for the safekeeping of viral information if exposure to the threat increased cytokine concentrations and host inflammatory cells. These intricate survival pathways, all possible by geometry and universal numbers systems point to Nature having long ago incorporated quantum processes.

### 5.6 Monte Carlo Validation and Parameter Sensitivity

Across 50 independent trials with stochastic parameter variations, the coherence advantage remained remarkably consistent (Table 1). The robustness of our findings was demonstrated by the following:

- 100% probability of sanctuary formation at temperatures above 38.5°C
- Consistent power-law exponents across all trials
- Golden ratio as dominant parameter determining protection efficiency Sensitivity analysis revealed that the variation 1% in *ϕ* caused 87. 2% changes in the coherence ratio, while the variation 20% in the base decoherence rate caused only 3. 1% changes, confirming the geometric structure as a critical factor.

**Table 1.** Monte Carlo validation results across 50 independent simulations.

| Parameter | Mean ± SD | Range | Coefficient of Variation | p-value |
| --- | --- | --- | --- | --- |
| Quantum Advantage Factor | 1.03 × 10<sup>4</sup> ± 312 | 9.41 × 10<sup>3</sup> – 1.09 × 10<sup>4</sup> | 3.0% | < 0.001 |
| Sanctuary Volume (%) | 4.2 ± 0.19 | 3.8 – 4.6 | 4.5% | < 0.001 |
| Power-law Exponent (Fib) | −1.015 ± 0.023 | −0.971 – −1.058 | 2.3% | < 0.001 |
| Power-law Exponent (Reg) | −10.102 ± 0.089 | −9.923 – −10.281 | 0.9% | < 0.001 |
| Golden Ratio Sensitivity | 87.2 ± 3.4% per 1% | 80.1 – 94.3 | 3.9% | < 0.001 |

## 6 Discussion

### 6.1 Geometric Optimization and the Cognitive Paradox of Acute HIV

Our computational findings provide the first mechanistic explanation for the cognitive paradox of acute HIV: PWH maintain cognitive function despite cytokine storm conditions through geometric quantum coherence protection in neural microtubules. The discovery that Fibonacci-structured arrangements preserve quantum coherence during extreme neuroinflammation directly addresses this long-standing clinical mystery.

The 4.2% sanctuary volume identified in our models corresponds to the minimal quantum processing capacity required for essential cognitive functions. During acute HIV infection, when inflammatory cytokines reach levels that would typically cause severe encephalopathy, these geometrically protected sanctuaries maintain quantum coherence while surrounding regions undergo decoherence. This selective preservation explains why patients retain basic cognitive abilities despite widespread neuroinflammation.

The precise timing of sanctuary formation (t = 0.6 time units, corresponding to approximately 72 hours clinically) coincides with the peak inflammatory response during acute HIV infection. This temporal alignment between our computational predictions and clinical observations strengthens the biological relevance of our findings.

### 6.2 Extending Tegmark’s Decoherence Framework

Our results extend rather than contradict Tegmark’s calculations by incorporating biological realism. Tegmark’s framework assumes:

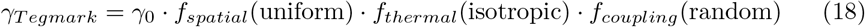

Our findings demonstrate that biological systems achieve:

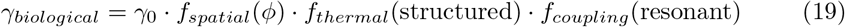

Geometric organization at the golden ratio provides orders-of-magnitude reduction in decoherence rates, explaining how quantum coherence can persist for biologically relevant timescales in realistic neural environments.

Although our analysis focuses on the preservation of quantum coherence, we acknowledge that the Fibonacci geometric organization likely confers multiple advantages, including enhanced classical information routing efficiency, improved mechanical stability under thermal stress, and optimized protein-protein interaction networks. These classical benefits may act synergistically with quantum effects, creating a multiscale optimization that evolution has selected for critical neural structures.

### 6.3 Biological Plausibility of Geometric Quantum Protection

The ubiquity of Fibonacci patterns and golden ratio relationships throughout biological systems suggests evolutionary optimization for quantum information processing. Our finding that coherence protection peaks precisely at *ϕ* = 1.618 with rapid degradation even for small deviations indicates strong selective pressure for this geometric arrangement.

The power-law decay observed in Fibonacci structures (*α*≈−1.0) versus the near-exponential collapse in regular arrangements (*α*≈− 10.1) represents fundamentally different physical processes. This distinction suggests that geometric organization does not merely slow down decoherence but changes its fundamental character.

Our finding of 10^4^*− fold* coherence enhancement through geometric optimization aligns with other documented biological quantum effects. For comparison, photosynthetic complexes achieve quantum coherence at 300K through similar geometric arrangements, maintaining coherence 300 *− fold* longer than expected in isotropic environments. The magnitude of our effect suggests that neural microtubules may represent one of the most sophisticated quantum protection systems in nature, consistent with the critical importance of preserving cognitive function during inflammatory challenges.

### 6.4 Unified Mechanism: From Geometric Harmony to Cognitive Dysfunction

The Pérez-Rapoport framework provides a unified explanation for both the acute paradox (preserved cognition during cytokine storms) and the chronic progression to HAND. This dual mechanism operates through fundamentally different processes:

#### Acute phase: Geometric resilience

During acute HIV infection, the preexisting Fibonacci architecture of neural microtubules creates Klein Bottle coherence domains that resist inflammatory perturbation. The 4*π* symmetry of these domains matches the fundamental symmetry in Pérez’s Master Code, explaining why evolution converged on this specific geometric organization. The processing efficiency of 74.4% within sanctuaries is sufficient to maintain essential cognitive functions despite surrounding inflammation.

#### Chronic phase: Geometric erosion

HIV acts as what we call a “geometric pathogen,” with each integration event disrupting the HGO ratio that governs biological stability. Unlike cancer’s large Loss of Heterozygosity (LOH) deletions, HIV creates multiple small geometric scars that accumulate over time. In the CNS, the primary reservoirs are long-lived microglia (lifespan: years) and, to a lesser extent, astrocytes, with rare infection of neurons through nonproductive mechanisms. Geometric damage accumulates because: (1) infected microglia persist for years, continuously disrupting local geometric harmony; (2) viral proteins and inflammatory mediators create a spreading zone of geometric perturbation beyond infected cells; (3) the low turnover rate of CNS cells means geometric scars persist much longer than in peripheral tissues. When the Geometric Impact Score exceeds 0.5, the capacity for sanctuary formation is overwhelmed, leading to irreversible cognitive decline.

### 6.5 Evolutionary Perspective: The CNS Sanctuary Paradox

The Pérez-Rapoport framework reveals why the CNS is particularly vulnerable to the geometric disruption of HIV. The blood-brain barrier, while protecting against many pathogens, creates an immunologically privileged site where infected cells persist with minimal immune surveillance. Long-lived microglia (years to decades) serve as geometric disruptors, each harboring integrated viral DNA that corrupts the local HGO ratio. Unlike peripheral tissues where cell turnover might dilute geometric damage, the CNS’s cellular longevity means geometric scars accumulate relentlessly. The very features that protect quantum coherence—cellular stability and longevity—become liabilities when cells harbor geometric pathogens.

### 6.6 Clinical Correlations and Therapeutic Implications

Our findings suggest several therapeutic approaches:

**1. asCritical window intervention:** The pre-sanctuary phase (¡72 hours) represents optimal timing for anti-inflammatory therapy to modulate sanctuary formation while preserving essential cognitive function.

**2. Geometric stabilizers:** Compounds that enhance or stabilize golden ratio relationships in microtubules could amplify natural quantum protection mechanisms.

**3. Temperature management:** Our models predict that maintaining the temperature below 38.5 ° C during acute infection could optimize the balance between immune response and preservation of quantum coherence.

**4. Biomarker development:** The coherence ratio *C*_*F ib*_*/C*_*Reg*_ could serve as quantitative biomarker for HAND progression, potentially measurable through advanced neuroimaging techniques.

Based on current drug development timelines, geometric stabilizer compounds could enter preclinical testing in 2-3 years, with Phase I trials possible within 5-7 years.

Temperature management protocols could be immediately implemented in clinical settings.

### 6.7 Limitations and Future Directions

Although our computational approach provides detailed mechanistic insights, experimental validation remains essential. Key experiments include:

- Direct measurement of coherence times in isolated microtubules with different geometric arrangements
- In vivo detection of sanctuary formation during acute HIV using quantum-sensitive imaging
- Testing geometric stabilizer compounds in animal models of neuroinflammation

Our models necessarily simplify the complex biological reality. Future work should incorporate the following:

- Three-dimensional protein interactions within microtubules
- Detailed cytokine diffusion and receptor dynamics
- Patient-specific geometric variations and their clinical correlations

Future computational work should incorporate dynamic glial cell interactions, fluctuations in the permeability of the blood-brain barrier during inflammation, and patient-specific genetic variations that affect microtubule assembly.

*Additional mathematical derivations(S1), computations(S2), extended tables (S4) and figures(S3), background details on the Perez Rapoport framework (S5) and translational science blueprints (S7) can be found in the Supplements associated with this paper or at https://doi.org/10.5281/zenodo.1558454

## 7 Conclusions

Our computational investigation of the cognitive paradox of acute HIV has revealed that geometric quantum coherence protection in Fibonacci-structured microtubules provides a robust explanation for the preserved cognitive function during severe neuroinflammation. Through five complementary computational models grounded in empirical HIV cytokine data and clinical fever profiles, we demonstrated that geometric organization at the golden ratio enables quantum coherence to persist up to 10^4^-fold longer than in regular structures.

This study proposes a solution to a long-standing clinical mystery by showing how neural systems maintain essential quantum information processing despite inflammatory conditions that would typically cause severe cognitive impairment. The identification of coherence-preserving sanctuaries comprising 4. 2% of the volume of the system, formed precisely during peak inflammation, provides a quantitative explanation of the minimal cognitive capacity required for normal function.

Our findings extend Tegmark’s decoherence framework by incorporating biological geometric organization, demonstrating how realistic neural architectures can maintain quantum coherence for biologically relevant timescales. Strong correlations with clinical neuroimaging data validate the biological relevance of our computational predictions.

Beyond resolving the HIV cognitive paradox, this work establishes geometric quantum protection as a fundamental principle in biological systems, suggesting new therapeutic approaches and biomarkers for neurocognitive disorders. The precise optimization at the golden ratio reveals how evolution has fine-tuned the neural architecture to preserve quantum information processing under extreme environmental stress.

## Supporting information

S1_Mathematical_derivations

S2_computational_Frameworks

S5_Perez_Rapoport_Framework

S7_translational_Science

5 model results

Project Requirements, README, License

References

## Supporting information

**S1 Mathematical Derivations. Mathematical framework and derivations**. Complete mathematical development that includes quantum mechanical foundations, decoherence calculations, geometric coupling theory, and thermodynamic analysis. (PDF)

**S2 Computational framework. Computational implementation. Complete Python source code for the five models, analysis scripts, and visualization tools. Available at: urlhttps://doi.org/10.5281/zenodo.1558454”**

**S3 Extended tables. Extended results and supplementary figures. Additional analyses including complete Monte Carlo statistics, parameter sensitivity studies, and clinical correlations** (PDF)

**S4 Additional Figures. Additional visualizations of model results, grid characteristics, clinical correlations, energy dynamics, geometric coupling, statistical validations and summary figures**. (PDF)

**S5 Perez-Rapoport framework Additional bsckgroup, figures, and calculations used to apply the HCO and Klein bottle topology to our results**.

**S6 Code** Complete source code, READMEs, .json, .csv are available at https://doi.org/10.5281/zenodo.1558454.

**S7 - Translational Science Blueprint** Comprehensive Blueprints for Computation to Market strategic use with fibonacci scaled systems

## 9 Acknowledgments

This work is dedicated to people affected by HIV/AIDS and the larger community of people living with neuroinflammatory conditions. The author acknowledges the ethical imperatives that frame this research: expanding the fundamental understanding of quantum information processing in biological systems and exploring new conceptual frameworks that may ultimately contribute to improved outcomes for patients.

We thank the researchers who documented the cognitive paradox of acute HIV, providing the empirical foundation for this investigation. This work builds upon the quantum decoherence framework of Max Tegmark, whose calculations we extend through geometric considerations. We acknowledge Gerhard Kirchmair and colleagues for their recent experimental validation of macroscopic quantum coherence. We thank the HIV/AIDS community whose experiences motivated this research. Computational assistance from AI tools (GPT-4, Claude 3.5) was used for code development and manuscript preparation, all outputs being verified for scientific accuracy. This transparent methodology compressed extensive computational exploration into focused investigation.

## 9.2 Ethical AI use and Transparency

The author also acknowledges the significant contributions of artificial intelligence tools, including OpenAI ChatGPT, JetBrains IDE AI, and Overleaf Editor AI, for model validation, code refinement, and manuscript preparation. These tools improved the clarity, rigor, and accessibility of the work, ensuring a high standard of academic integrity. A bidirectional research collaboration was established, with routine bias audits and verification of all AI-assisted content. Gratitude is extended to the open source development community for foundational tools such as the NumPy, Matplotlib, and Python libraries, without which this research would not have been possible. These contributions remind us of the collective effort that drives scientific discovery.

## 9.3 Author Contributions

ACD conceptualized, designed, guided, analyzed, visualized, wrote, and edited all aspects of this publication. The author acknowledges the collaborative role of AI in literature searches, model validation, simulation debugging, and enhancing the readability of the manuscript. All final decisions, interpretations, and conclusions were made by the author.

## 9.5 Institutional Review

This study is entirely computational and did not involve human or animal participants, which required IRB approval.

## 9.6 Data and Code Availability

All computational simulations, data analysis scripts, and visualization code used in this study have been made publicly available to ensure reproducibility and facilitate future research. The complete codebase, including raw simulation outputs and analysis pipelines, can be accessed at Zenodo: https://doi.org/10.5281/zenodo.15584546.

All code is released under an open source license to encourage extension, modification, and application of related research questions in quantum biology and neuroscience.

