## Supplementary material for "Quantum Coherence Preservation in Fibonacci-Structured Microtubules During HIV-Induced Neuroinflammation": S2_computational_Frameworks

### Geometric Quantum Coherence Protection in Fibonacci-Structured Microtubules: A Quantum Solution to the Cognitive Paradox of Acute HIV-associated Neuroinflammation

Adrian Charles (AC) Demidont,DO[1*]

**1** Nyx Dynamics, Independent Researcher, Fairfield, Connecticut, United States of America

*

### Supplementary Information 2: Computational Frameworks

#### 1 Table of Contents

1. Quantum Mechanical Foundation 2. Decoherence Framework 3. 3-coherence-energy-transfer 4. 4-energy-conservation-proofs 5. 5-golden-ratio-optimization 6. 6-power-law-analysis 7. 7-thermodynamic-consistency 8. 8-statistical-framework

—

### Quantum Mechanical Foundation

### Modified Schroedinger Equation

The fundamental equation governing quantum evolution in microtubules incorporates both unitary evolution and environmental decoherence:

$$i\hbar\frac{\partial\psi}{\partial t} = H_0\psi + H_{dec}\psi + H_{geom}\psi + H_{int}\psi$$

Where: - $H_0$ = Unperturbed Hamiltonian - $H_{dec}$ = Decoherence Hamiltonian - $H_{geom}$ = Geometric coupling Hamiltonian - $H_{int}$ = Interaction Hamiltonian

##### Unperturbed Hamiltonian

The baseline Hamiltonian for microtubule quantum states:

$$H_0 = -\frac{\hbar^2}{2m_{eff}}\nabla^2 + V_0(r)$$

Where the effective mass $m_{eff}$ accounts for collective excitations:

$$m_{eff} = \frac{M_{tubulin}}{N_{coherent}}$$

with $M_{tubulin} = 55$ kDa and $N_{coherent}$ the number of coherently coupled tubulin dimers. The confining potential:

$$V_0(r) = \begin{cases} 0 & r < R_{inner} \\ V_{depth}\left[1 - \exp\left(-\frac{r-R_{inner}}{\lambda}\right)\right] & R_{inner} \le r \le R_{outer} \\ V_{depth} & r > R_{outer} \end{cases}$$

##### Decoherence Hamiltonian

Following Tegmark's framework with modifications:

$$H_{dec} = -i\hbar \sum_j \gamma_j(T, r)|j\rangle\langle j|$$

The position- and temperature-dependent decoherence rate:

$$\gamma_j(T, r) = \gamma_0 \exp\left(\frac{E_a}{k_B T}\right) \cdot f_{spatial}(r) \cdot g_{cytokine}(r, t)$$

Where: - $\gamma_0 = 10^{13}$ s$^{-1}$ (baseline rate) - $E_a = 0.1$ eV (activation energy) - $f_{spatial}(r)$ = spatial modulation factor - $g_{cytokine}(r, t)$ = inflammatory enhancement

##### Geometric Coupling Hamiltonian

The key innovation - geometric structure modifies quantum coupling:

$$H_{geom} = \sum_{i,j} J_{ij}^{(F)}|i\rangle\langle j| + \text{h.c.}$$

Coupling strengths for Fibonacci arrangements:

$$J_{ij}^{(F)} = J_0 \exp\left(-\frac{|r_i - r_j|}{\xi}\right) \cdot \mathcal{R}(\phi_{ij})$$

The golden ratio resonance function:

$$\mathcal{R}(\phi_{ij}) = \exp\left[-\alpha\left(\frac{|r_i - r_j|/|r_i|}{\phi} - 1\right)^2\right]$$

The peak resonance occurs when the spatial ratios match $\phi = 1.618$.

##### Interaction Hamiltonian

Enables energy transfer between collapsing and protected regions:

$$H_{int} = \sum_{k \in \text{reg}} \sum_{l \in \text{Fib}} V_{kl}|k\rangle\langle l| + \text{h.c.}$$

Transfer matrix elements:

$$V_{kl} = V_0 \frac{\langle k|\nabla^2|k\rangle}{\langle l|\nabla^2|l\rangle} \cdot \exp\left(-\frac{|r_k - r_l|}{R_c}\right)$$

—

#### Decoherence Framework

##### Master Equation Approach

Evolution of the density matrix:

$$\frac{d\rho}{dt} = -\frac{i}{\hbar}[H, \rho] + \mathcal{L}[\rho]$$

The Lindblad superoperator:

$$\mathcal{L}[\rho] = \sum_k \gamma_k \left(L_k \rho L_k^\dagger - \frac{1}{2}\{L_k^\dagger L_k, \rho\}\right)$$

#### Coherence Measures 44

Von Neumann entropy: 45

$$S = -\text{Tr}(\rho \ln \rho) = -\sum_i \lambda_i \ln \lambda_i$$

Quantum coherence (l1-norm): 46

$$C_{l_1}(\rho) = \sum_{i \neq j} |\rho_{ij}|$$

Relative entropy of coherence: 47

$$C_r(\rho) = S(\rho_{diag}) - S(\rho)$$

#### Modified Tegmark Timescale 48

Original Tegmark decoherence time: 49

$$\tau_{Tegmark} = \frac{\hbar}{k_B T}\sqrt{\frac{m}{M}} \approx 10^{-13}\text{ s}$$

Modified for geometric structure: 50

$$\tau_{modified} = \tau_{Tegmark} \cdot F_{geom}(\phi) \cdot F_{thermal}(T) \cdot F_{coupling}$$

Where:

$$F_{geom}(\phi) = \exp\left[\beta(\phi - 1)^2\right]$$

$$F_{thermal}(T) = \frac{1}{1 + (T/T_c)^n}$$

$$F_{coupling} = \prod_i \left(1 + \frac{J_i}{k_B T}\right)$$

— 51

### Coherence Energy Transfer 52

#### Analogy Development 53

Classical Hawking radiation temperature: 54

$$T_H = \frac{\hbar c^3}{8\pi k_B G M}$$

Biological analog - "coherence temperature": 55

$$T_C = \frac{\hbar \gamma_{collapse}}{2\pi k_B N_{states}}$$

#### Energy Flux Calculation

Energy emitted from collapsing region:

$$\frac{dE_{emit}}{dt} = \sigma A T_C^4$$

Where $\sigma$ is an effective Stefan-Boltzmann constant:

$$\sigma_{bio} = \frac{2\pi^5 k_B^4}{15h^3c^2} \cdot \frac{v_{phonon}^2}{c^2}$$

#### Absorption in Fibonacci Regions

Absorption cross-section enhanced by golden ratio:

$$\sigma_{abs}(\omega) = \sigma_0 \frac{(\omega/\omega_\phi)^2}{1 + (\omega/\omega_\phi - 1)^2}$$

Peak absorption at $\omega_\phi = 2\pi c/\lambda_\phi$ where $\lambda_\phi = a_0 \cdot \phi$.

#### Net Energy Transfer

Energy balance equation:

$$\frac{dE_{Fib}}{dt} = \eta \int_\Omega \frac{dE_{emit}}{dtd\Omega} d\Omega - \Gamma_{loss} E_{Fib}$$

Harvesting efficiency:

$$\eta = \frac{\int \sigma_{abs}(\omega) n(\omega) d\omega}{\int n(\omega) d\omega}$$

where $n(\omega)$ is the emission spectrum.
—

### Energy Conservation Proofs

#### Total Energy Conservation

Define total system energy:

$$E_{total} = \langle H_0 \rangle + E_{coherence} + E_{thermal}$$

Coherence energy:

$$E_{coherence} = \sum_{i \neq j} |J_{ij}||\rho_{ij}|^2$$

Time derivative:

$$\frac{dE_{total}}{dt} = \frac{d}{dt}\text{Tr}(\rho H) = \text{Tr}\left(\frac{d\rho}{dt} H\right) + \text{Tr}\left(\rho \frac{dH}{dt}\right)$$

Using the master equation:

$$\frac{dE_{total}}{dt} = -\sum_k \gamma_k \langle L_k^\dagger L_k \rangle + \frac{\partial V_{cytokine}}{\partial t}$$

#### Local Energy Flow

Energy current density:

$$\vec{J}_E = \frac{\hbar}{2mi}\left[\psi^*(\nabla H)\psi - \psi(H\nabla)\psi^*\right]$$

Continuity equation:

$$\frac{\partial u_E}{\partial t} + \nabla \cdot \vec{J}_E = S_E$$

where $S_E$ represents sources/sinks.

#### Sanctuary Energy Balance

Within sanctuary volume $V_s$:

$$\frac{d}{dt}\int_{V_s} u_E dV = -\oint_{\partial V_s} \vec{J}_E \cdot d\vec{A} + \int_{V_s} S_E dV$$

Boundary conditions ensure:

$$\oint_{\partial V_s} \vec{J}_E \cdot d\vec{A} > 0$$

(net energy influx during collapse phase).

—

### Golden Ratio Optimization

#### Variational Principle

Define coherence functional:

$$\mathcal{F}[\xi] = \int_0^{t_f} C(\xi, t) dt$$

Euler-Lagrange equation:

$$\frac{\delta \mathcal{F}}{\delta \xi} = 0$$

Yields optimal scaling:

$$\xi_{opt} = \arg\max_{\xi} \mathcal{F}[\xi]$$

#### Analytical Solution

For power-law coherence decay:

$$C(\xi, t) = C_0 t^{-\alpha(\xi)}$$

Exponent dependence:

$$\alpha(\xi) = \alpha_0 + \beta(\xi - \phi)^2$$

Optimization condition:

$$\frac{d}{d\xi}\int_0^{t_f} t^{-\alpha(\xi)}dt = 0$$

Solution:

$$\xi_{opt} = \phi + \frac{\alpha_0}{2\beta t_f}\ln\left(\frac{t_f}{t_0}\right)$$

For large $t_f$: $\xi_{opt} \to \phi$.

#### Resonance Theory

Fibonacci scaling creates resonance condition:

$$\omega_n = \omega_0\frac{F_{n+1}}{F_n}$$

In the limit:

$$\lim_{n\to\infty}\frac{F_{n+1}}{F_n} = \phi$$

Quality factor:

$$Q = \frac{\omega_\phi}{\Delta\omega} = \frac{\phi}{\xi - \phi}$$

Diverges as $\xi \to \phi$.

—

### Power Law Analysis

#### Coherence Decay Forms

General form:

$$C(t) = C_0 f(t/\tau)$$

Power law:

$$f(x) = x^{-\alpha}$$

Exponential:

$$f(x) = e^{-x}$$

Stretched exponential:

$$f(x) = e^{-x^\beta}$$

#### Fitting Procedure

Log-transform for power law:

$$\ln C = \ln C_0 - \alpha \ln t$$

Linear regression:

$$\alpha = -\frac{\sum_i (\ln t_i - \overline{\ln t})(\ln C_i - \overline{\ln C})}{\sum_i (\ln t_i - \overline{\ln t})^2}$$

Goodness of fit:

$$R^2 = 1 - \frac{\sum_i (\ln C_i - \ln \hat{C}_i)^2}{\sum_i (\ln C_i - \overline{\ln C})^2}$$

#### Phase-Dependent Exponents

Early phase (sanctuary formation):

$$\alpha_{early} = \alpha_0 + \delta\alpha \cdot H(t_{sanctuary} - t)$$

Late phase (established sanctuary):

$$\alpha_{late} = \alpha_0 - \epsilon$$

Where $H$ is the Heaviside function.

—

### Thermodynamic Consistency

#### Entropy Production

Total entropy:

$$S_{total} = S_{system} + S_{environment}$$

Rate of change:

$$\frac{dS_{total}}{dt} = \frac{dS_{system}}{dt} + \frac{dS_{environment}}{dt}$$

System entropy from density matrix:

$$S_{system} = -k_B \text{Tr}(\rho \ln \rho)$$

#### Second Law Compliance

For isolated system:

$$\frac{dS_{total}}{dt} \geq 0$$

Decomposition:

$$\frac{dS_{total}}{dt} = \sigma_{reg} + \sigma_{Fib} + \sigma_{boundary}$$

Where: - $\sigma_{reg} > 0$ (collapse increases entropy) - $\sigma_{Fib} < 0$ (coherence decreases entropy) - $\sigma_{boundary} > |\sigma_{Fib}|$ (boundary formation compensates)

#### Free Energy Landscape

Helmholtz free energy:

$$F = U - TS$$

For Fibonacci structures:

$$F_{Fib} = U_0 - T(S_0 - \Delta S_{coherence})$$

Lower free energy drives sanctuary formation:

$$\Delta F = F_{Fib} - F_{reg} < 0$$

#### Maximum Work Principle

Maximum work extractable:

$$W_{max} = -\Delta F = k_B T \ln\left(\frac{Z_{Fib}}{Z_{reg}}\right)$$

Partition functions:

$$Z = \sum_n e^{-E_n/k_B T}$$

Golden ratio maximizes $Z_{Fib}/Z_{reg}$.
—

### Statistical Framework

#### Monte Carlo Methodology

Parameter distributions:

$$p(\theta) = \prod_i p_i(\theta_i)$$

Individual distributions: - $D_0 \sim \mathcal{N}(\mu_{D_0}, \sigma_{D_0}^2)$ - $\Gamma_0 \sim$ LogNormal$(\mu_\Gamma, \sigma_\Gamma^2)$ - $T \sim$ Beta$(a_T, b_T)$ (scaled) - $\phi \sim \mathcal{N}(1.618, \sigma_\phi^2)$

#### Sensitivity Analysis

Logarithmic derivatives:

#### Volume Calculation

Sanctuary volume:

$V_s(t) = \int_{|\psi|^2 > \psi^2_{crit}} dV$

Critical threshold:

$\psi^2_{crit} = \psi_0^2 \exp(-\gamma_{eff} t)$

#### Stability Analysis

Lyapunov exponent for boundary:

$\lambda = \lim_{t \to \infty} \frac{1}{t} \ln \left( \frac{|\delta r(t)|}{|\delta r(0)|} \right)$

Stable sanctuary: $\lambda < 0$

#### Formation Dynamics

Rate equation:

$\frac{dV_s}{dt} = v_{in} A_{in} - v_{out} A_{out}$

Where: - $v_{in}$ = inward velocity at inner boundary - $v_{out}$ = outward velocity at outer boundary - $A$ = surface areas

Equilibrium: $v_{in} A_{in} = v_{out} A_{out}$

—

### Wave Function Analysis

#### Spatial Modes

Expansion in eigenmodes:

$\psi(r,t) = \sum_n c_n(t) \phi_n(r)$

Mode equations:

$H_0 \phi_n = E_n \phi_n$

#### Mode Coupling

Evolution equations:

$i\hbar\frac{dc_n}{dt} = E_n c_n + \sum_m V_{nm} c_m - i\hbar \sum_m \Gamma_{nm} c_m$

Coupling matrix:

$V_{nm} = \langle \phi_n | H_{geom} | \phi_m \rangle$

Decoherence matrix:

$\Gamma_{nm} = \gamma_n \delta_{nm} + \gamma_{nm}^{(off)}$

#### Coherent Superposition

Two-mode approximation:

$|\psi\rangle = \alpha|0\rangle + \beta|1\rangle$

Coherence measure:

$C = 2|\alpha\beta| = 2\sqrt{p(1-p)}$

where $p = |\alpha|^2$.

#### Decoherence Dynamics

Density matrix evolution:

$\rho(t) = \begin{pmatrix} |\alpha|^2 & \alpha\beta^* e^{-\Gamma t} \\ \alpha^*\beta e^{-\Gamma t} & |\beta|^2 \end{pmatrix}$

Purity:

$\mathcal{P}(t) = \text{Tr}(\rho^2) = |\alpha|^4 + |\beta|^4 + 2|\alpha\beta|^2 e^{-2\Gamma t}$

—

### Quantum Information Measures

#### Entanglement Entropy

For bipartite system:

$S_E = -\text{Tr}(\rho_A \ln \rho_A)$

where $\rho_A = \text{Tr}_B(\rho_{AB})$.

#### Quantum Fisher Information

For parameter estimation:

$F_Q(\rho, \theta) = \text{Tr}(\rho L_\theta^2)$

where $L_\theta$ is symmetric logarithmic derivative.

#### Quantum Mutual Information

Between regions A and B:

$I(A:B) = S(\rho_A) + S(\rho_B) - S(\rho_{AB})$

#### Channel Capacity

Maximum information transmission:

$C = \max_{p(x)} I(X:Y)$

For Fibonacci channels:

$C_{Fib} = \log_2(1 + \text{SNR} \cdot F(\phi))$

where $F(\phi)$ is the enhancement factor.

—

### Numerical Methods

#### Time Evolution

Split-operator method:
$\psi(t+\Delta t) = e^{-iH_0\Delta t/2\hbar} e^{-iV\Delta t/\hbar} e^{-iH_0\Delta t/2\hbar}\psi(t) + O(\Delta t^3)$

#### Spatial Discretization

Finite difference for Laplacian:
$\nabla^2\psi \approx \frac{\psi_{i+1}-2\psi_i+\psi_{i-1}}{\Delta x^2}$

#### Stability Criteria

CFL condition:
$\Delta t < \frac{\Delta x^2}{2D_{max}}$
Quantum CFL:
$\Delta t < \frac{m\Delta x^2}{\hbar}$

#### Convergence Tests

Richardson extrapolation:
$\psi_{exact} = \psi_{\Delta x} + C\Delta x^p + O(\Delta x^{p+1})$
Order verification:
$p = \frac{\ln(|\psi_{2\Delta x}-\psi_{\Delta x}|/|\psi_{\Delta x}-\psi_{\Delta x/2}|)}{\ln 2}$
—

### Asymptotic Analysis

#### Long-Time Behavior

For $t \to \infty$:
Regular grids: $C_{reg}(t) \sim A_0 t^{-\alpha_{reg}} \sim A_0 t^{-10.1}$
Fibonacci grids: $C_{Fib}(t) \sim B_0 t^{-\alpha_{Fib}} \sim B_0 t^{-1.0}$
Ratio: $\frac{C_{Fib}(t)}{C_{reg}(t)} \sim \frac{B_0}{A_0} t^{9.1} \to \infty$

#### Short-Time Expansion

For $t \ll \tau_c$:
$C(t) = 1 - \gamma t + \frac{\gamma^2 t^2}{2} - \frac{\gamma^3 t^3}{6} + O(t^4)$
The difference emerges at second order:
$\Delta C(t) = C_{Fib}(t) - C_{reg}(t) = (\gamma_{reg} - \gamma_{Fib})t + O(t^2)$

#### Critical Time

Sanctuary formation time from:
$\left.\frac{d^2V_s}{dt^2}\right|_{t=t_c} = 0$
Solution:
$t_c = \frac{1}{\gamma_{reg}} \ln\left(\frac{\gamma_{reg}}{\gamma_{Fib}}\right)$
For our parameters: $t_c = 0.6$ time units.
—

### Summary of Key Mathematical Results

#### Core Findings

1. **Power law exponents**: - Fibonacci: $\alpha = 1.0150 \pm 0.0023$ - Regular: $\alpha = 10.1023 \pm 0.0891$

2. **Golden ratio optimization**: - Peak at $\phi = 1.6180 \pm 0.0001$ - Curvature: $\partial^2 C/\partial\xi^2|_\phi = -847.3$

3. **Energy harvesting efficiency**: - $\eta = 0.744 \pm 0.023$ - Theoretical maximum: $\eta_{max} = 0.764$

4. **Coherence advantage**: - Short term (t=3): $4.5\times$ - Long term (t=120): $1.79 \times 10^{16}$

5. **Sanctuary volume**: - Mean: $4.2\% \pm 0.19\%$ - Formation probability: 100

#### Thermodynamic Compliance

Total entropy production rate:

$\frac{dS_{total}}{dt} = 3.7 \times 10^{-21}$ J/K/s $> 0$

Confirms Second Law compliance despite local coherence enhancement.

#### Clinical Correlations

Boundary radius vs. neuroimaging: - DTI FA: $r = 0.74$ ($p < 0.001$) - Cortical thickness: $r = -0.82$ ($p < 0.001$) - fMRI connectivity: $r = -0.76$ ($p < 0.001$)

#### Fundamental Constants

Derived biological constants: - $\gamma_{bio} = 1.23 \times 10^{13}$ s$^{-1}$ - $J_\phi = 2.7$ meV - $\xi_{correlation} = 8.3$ nm - $v_{phonon} = 1570$ m/s

These complete the mathematical framework for Coherence Energy Transfer in HIV-associated quantum decoherence.

### 2 Data Availability

Data are available at `https://doi.org/10.5281/zenodo.1558454` Code is available at `https://doi.org/10.5281/zenodo.1558454`
