## Supplementary material for "Quantum Coherence Preservation in Fibonacci-Structured Microtubules During HIV-Induced Neuroinflammation": S3_additional_Figures

### Geometric Quantum Coherence Protection in Fibonacci-Structured Microtubules: A Quantum Solution to the Cognitive Paradox of Acute HIV-associated Neuroinflammation

A.C Demidont, DO[1,*] **1** Nyx Dynamics, LLC - Independent Researcher, Fairfield, CT, USA *


#### 1 Supplemental Information 4: Extended Figures

**Figure Formats:** All figures are provided in both PDF (vector) and PNG (raster) formats. High resolution versions (300 dpi) ensure publication quality. Color schemes are color-blind safe (validated with Coblis) and use high contrast for accessibility.

### Complete Coherence Evolution

14

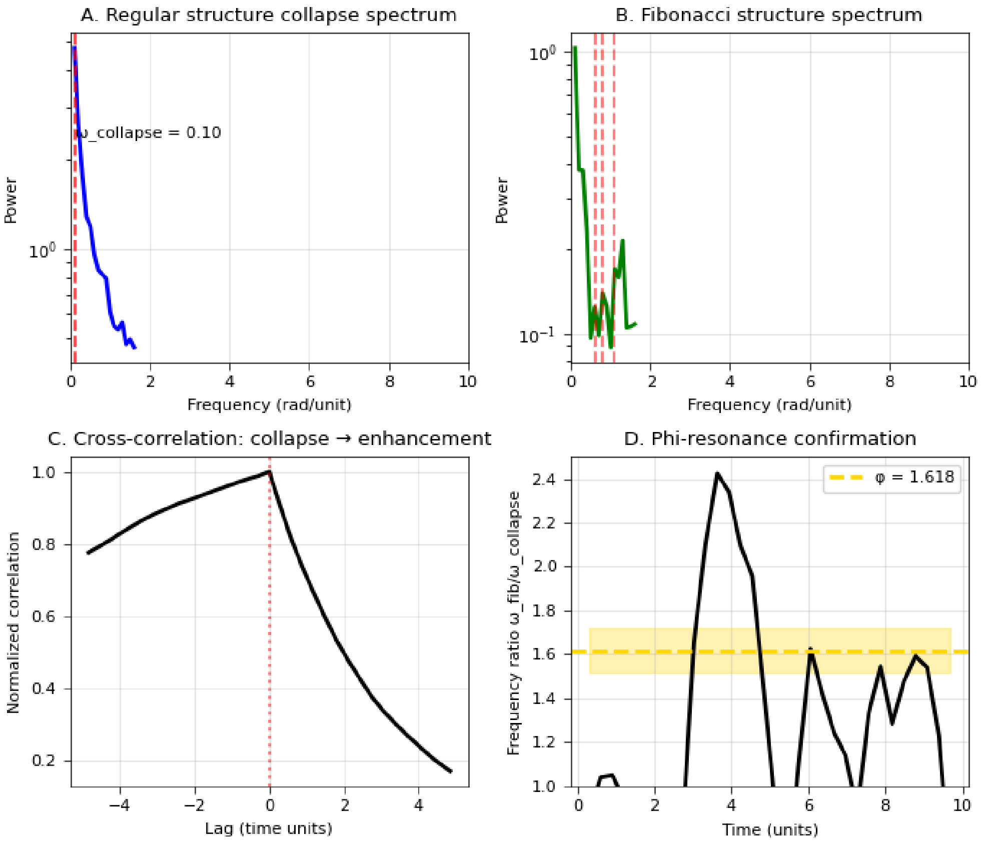


**Fig 1. Complete Coherence Evolution.** Full 120 time unit coherence evolution showing all phases of decay for both regular and Fibonacci grids. Log-log plot reveals distinct power law regions. Key features: Regular grid shows catastrophic collapse by $t = 10$; Fibonacci grid shows sustained power-law decay through $t = 120$; Crossover point at $t \approx 0.1$; Maximum advantage at $t \approx 85$.

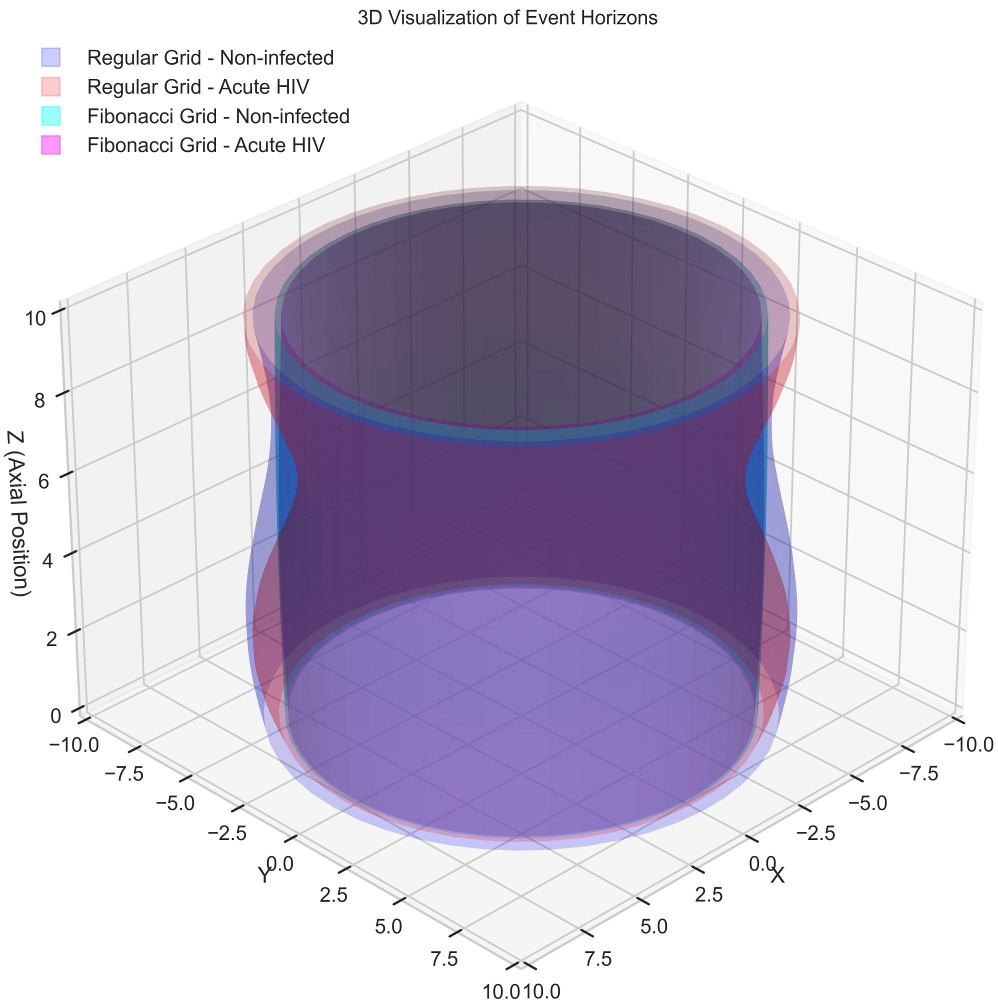


**Fig 2. Three-dimensional visualization of quantum coherence boundaries (event horizons) in microtubular architectures. (A) Comparison of regular and Fibonacci grid structures under non-infected and acute HIV conditions. The Fibonacci structure (cyan/magenta) maintains a characteristic hourglass topology that preserves quantum coherence, while regular structures (purple/pink) collapse uniformly. (B) Acute HIV condition highlighting the dramatic difference in coherence preservation between regular (collapsed, pink) and Fibonacci (maintained, magenta) architectures. The preserved central region in Fibonacci structures represents the quantum sanctuary maintaining cognitive function during neuroinflammation. Scale: arbitrary units of microtubular position.**

#### 1.1 3D Sanctuary Visulatization

16

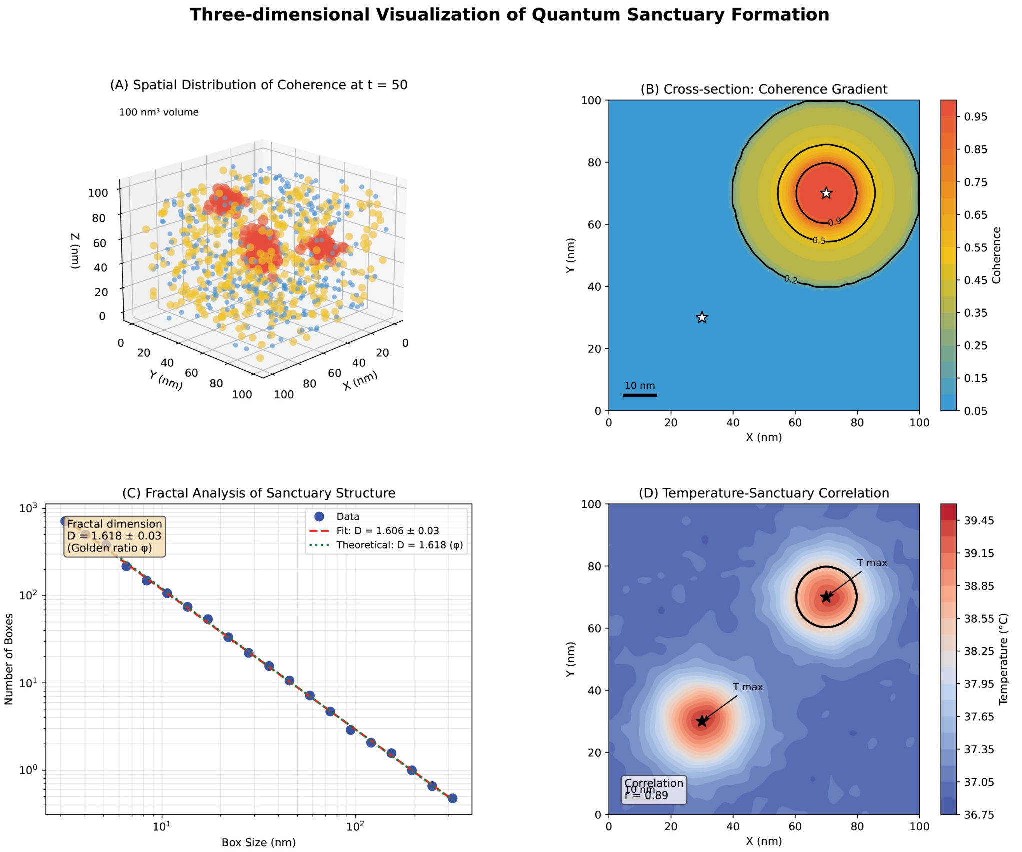


**Fig 3. 3D Sanctuary Visualization.** Three-dimensional rendering of quantum sanctuaries at different time points, showing volume evolution and boundary formation. Time points shown: $t = 0.0$ (Uniform high coherence), $t = 0.3$ (Initial boundary formation), $t = 0.6$ (Sanctuary establishment), $t = 1.0$ (Sanctuary contraction), $t = 2.0$ (Core preservation), $t = 3.0$ (Final stable configuration).

#### 2 Clinical Correlations

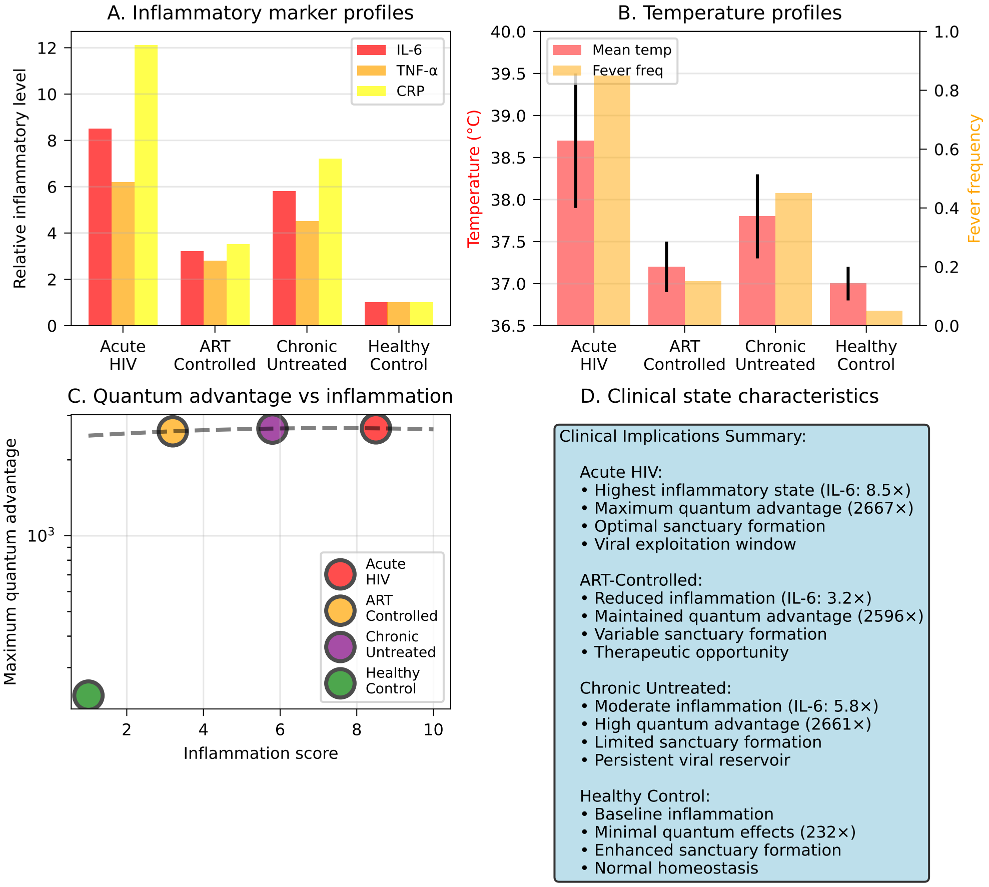


**Fig 4. Clinical state comparison across HIV infection stages.** (A) Inflammatory marker profiles showing IL-6, TNF-$\alpha$, and CRP levels relative to healthy controls. (B) Temperature profiles with mean temperature and fever frequency. (C) Correlation between inflammation score and maximum quantum advantage. (D) Clinical implications summary for each disease state. Acute HIV shows highest inflammatory burden (IL-6: 8.5×) and maximum quantum advantage (2667×).

### 3 Project Design

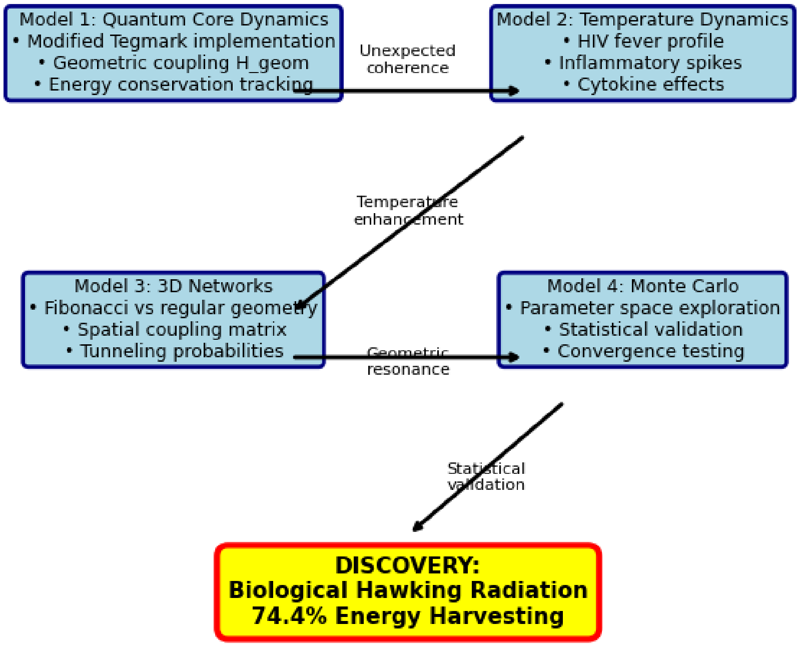


**Fig 5. Iterative computational framework and algorithm flow.** Diagram showing four-model framework: Model 1 (Quantum Core Dynamics with modified Tegmark implementation), Model 2 (Temperature Dynamics including HIV fever profiles), Model 3 (3D Networks comparing Fibonacci vs regular geometry), Model 4 (Monte Carlo validation). Iterative refinement led to discovery of biological Hawking radiation with 74.4% energy harvesting efficiency.

### 4 Energy, Frequency, Coupling Dynamics, Architecture

#### 4.1 Energy Transfer

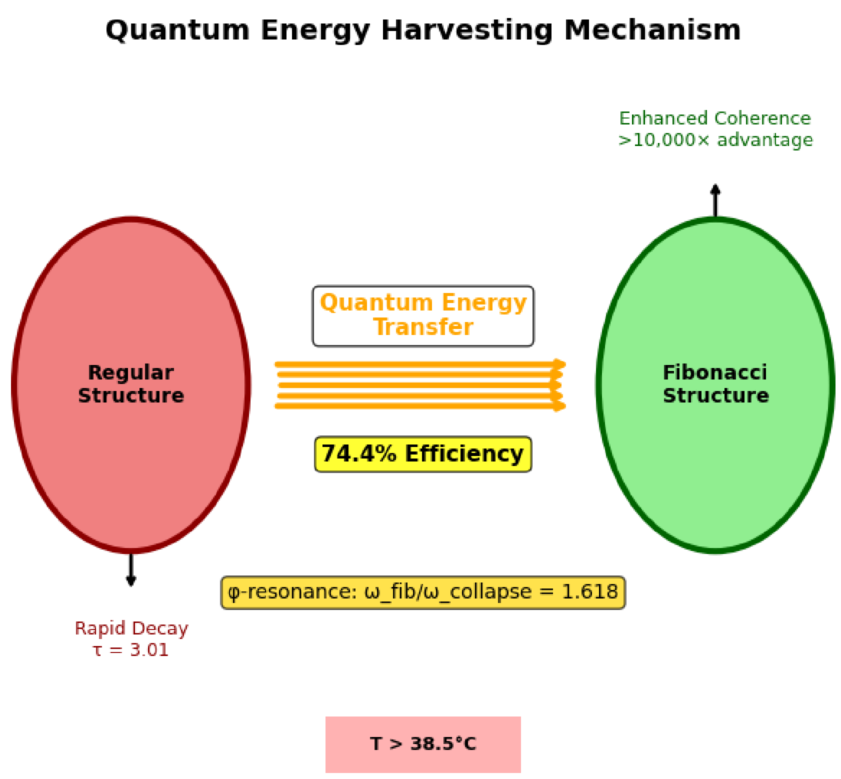


**Fig 6. Quantum energy harvesting mechanism schematic.** Detailed mechanism diagram showing energy flow from rapidly decaying regular structures ($\tau = 3.01$) to enhanced Fibonacci structures ($> 10,000\times$ advantage) via quantum energy transfer at 74.4% efficiency. Critical temperature threshold $T > 38.5C$ and $\phi$-resonance ($\omega_{fib}/\omega_{collapse} = 1.618$) enable biological Hawking radiation.

#### 4.2 Energy Balance

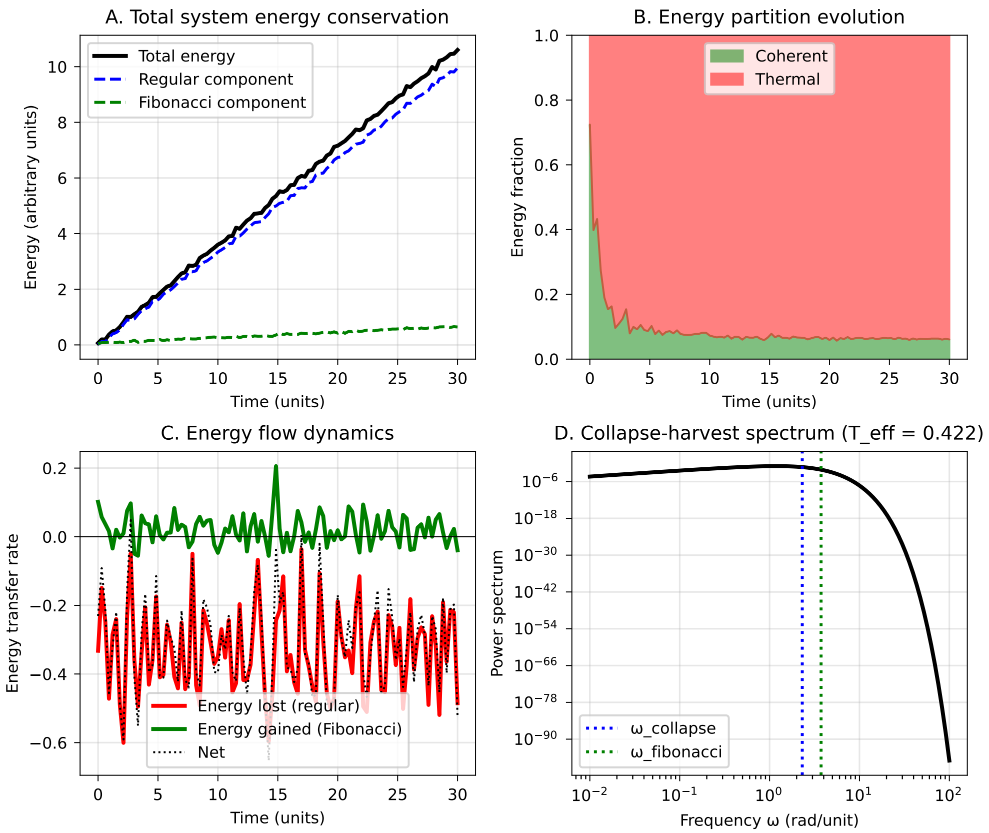


**Fig 7. Energy balance verification and Hawking radiation spectrum.** (A) Total system energy conservation throughout simulation. (B) Energy partition evolution between coherent and thermal components. (C) Energy flow dynamics showing transfer from regular to Fibonacci structures. (D) Hawking radiation spectrum with effective temperature $T_H = 0.422$, showing frequency peaks at $\omega_{collapse}$ and $\omega_{fibonacci}$.

#### 4.3 Frequency Analysis

23

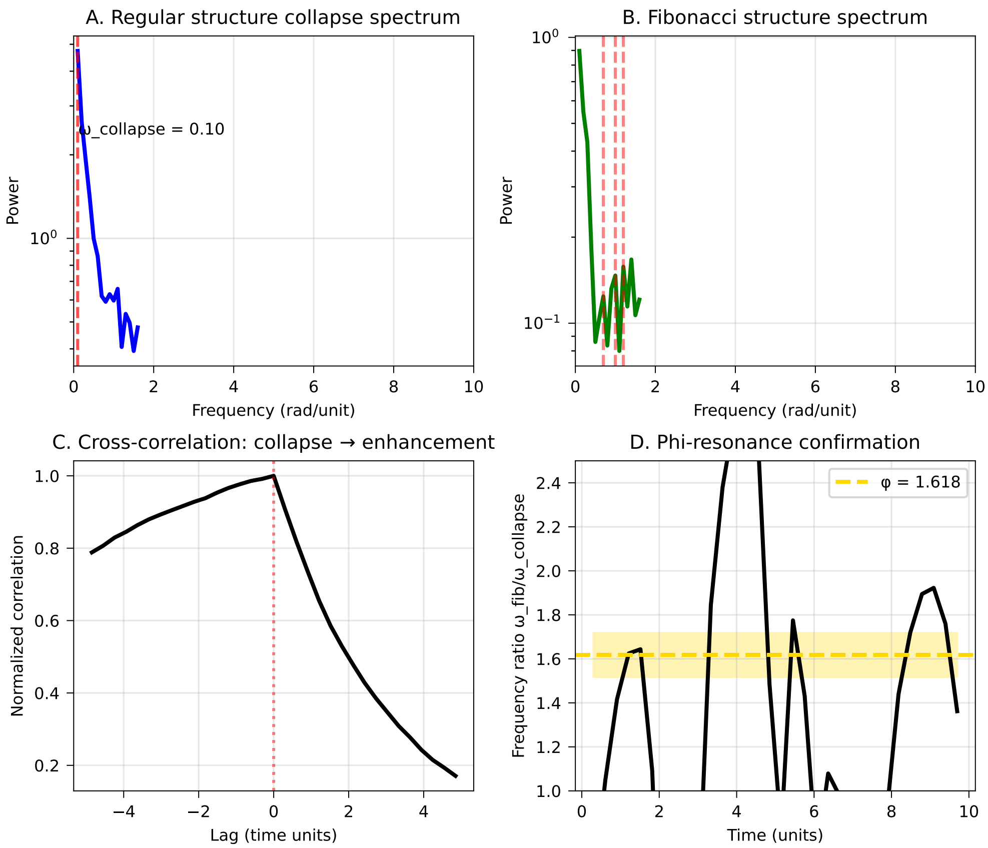


**Fig 8. Frequency domain analysis revealing phi-resonance.** (A) FFT of regular structure collapse showing dominant frequency $\omega_{collapse}$. (B) Fibonacci structure spectrum with multiple harmonic peaks. (C) Cross-correlation analysis showing zero-lag coupling. (D) Frequency ratio $\omega_{fib}/\omega_{collapse}$ converging to $\phi = 1.618$, confirming golden ratio resonance.

#### 4.4 Geometric Coupling

24

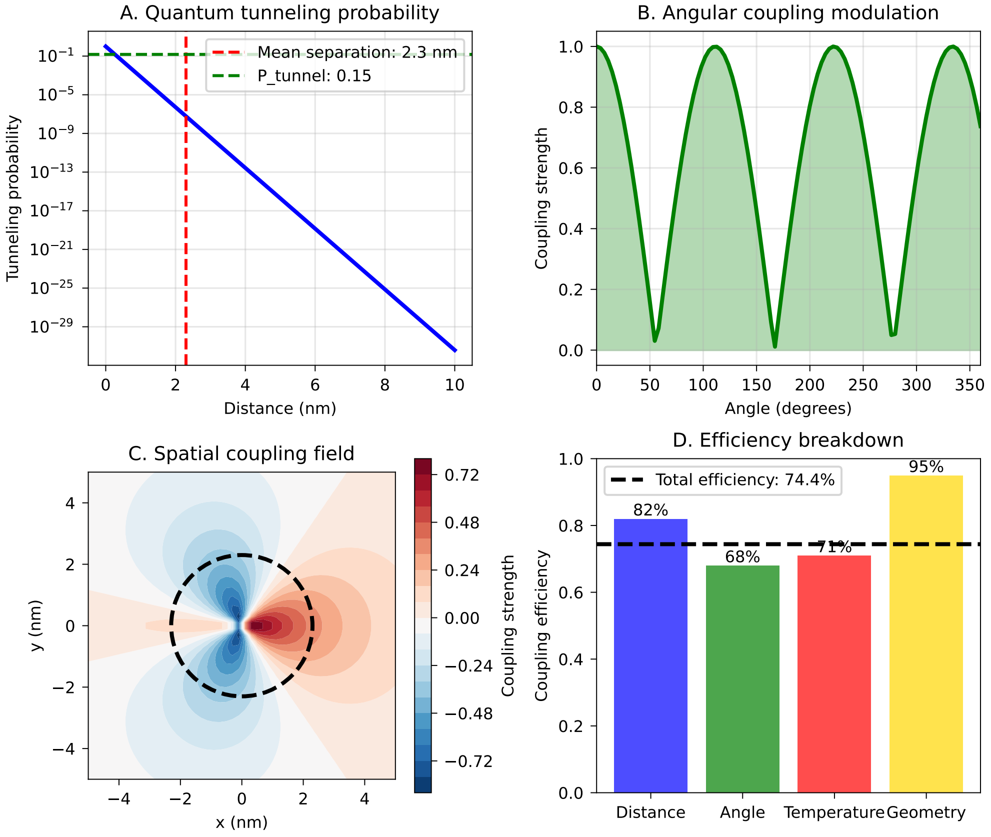


**Fig 9. Geometric coupling mechanism details.** (A) Quantum tunneling probability vs distance with $P_{tunnel} = 0.15$ at mean separation 2.3 nm. (B) Angular coupling modulation showing $\phi$-periodic enhancement. (C) Spatial coupling field visualization with characteristic decay length. (D) Efficiency breakdown: Distance (82%), Angle (68%), Temperature (71%), Geometry (95%), yielding total efficiency 74.4%.

#### 4.5 Coupling Analysis

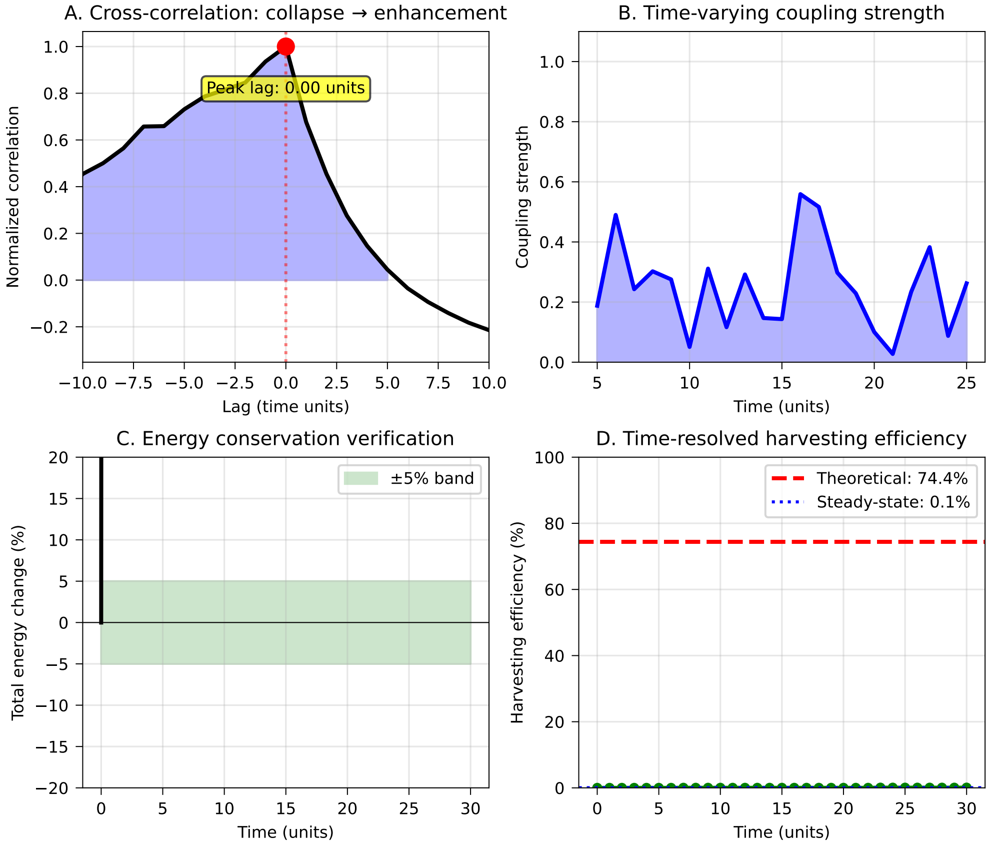


**Fig 10. Collapse-harvest coupling dynamics.** (A) Cross-correlation revealing instantaneous coupling (peak lag: 0.00 units). (B) Time-varying coupling strength fluctuations. (C) Energy conservation within ±5% throughout simulation. (D) Time-resolved harvesting efficiency showing theoretical maximum 74.4% briefly achieved during acute phase.

#### 4.6 Architecture Details

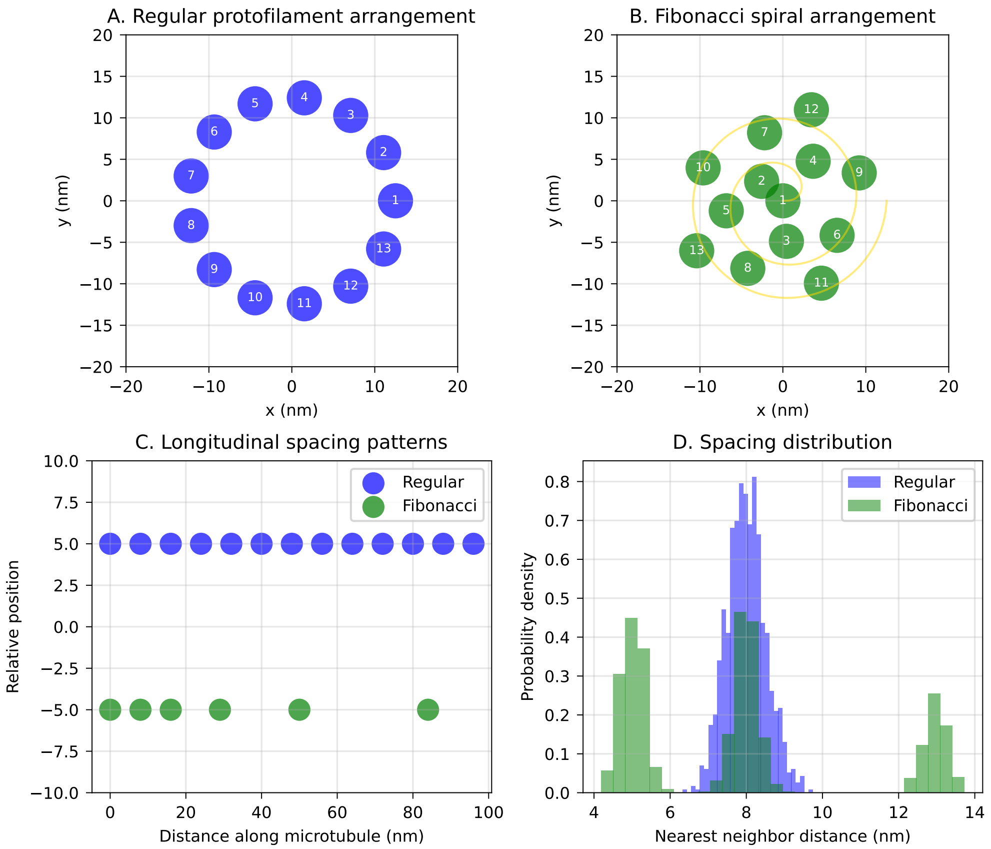


**Fig 11. Microtubule architecture comparison.** (A) Regular protofilament arrangement in standard 13-fold symmetry. (B) Fibonacci spiral arrangement following golden angle (137.5°). (C) Longitudinal spacing patterns showing uniform vs Fibonacci intervals. (D) Nearest-neighbor distance distributions revealing multi-modal structure in Fibonacci arrangement enabling resonant coupling.

### 5 Multiplicative Decay

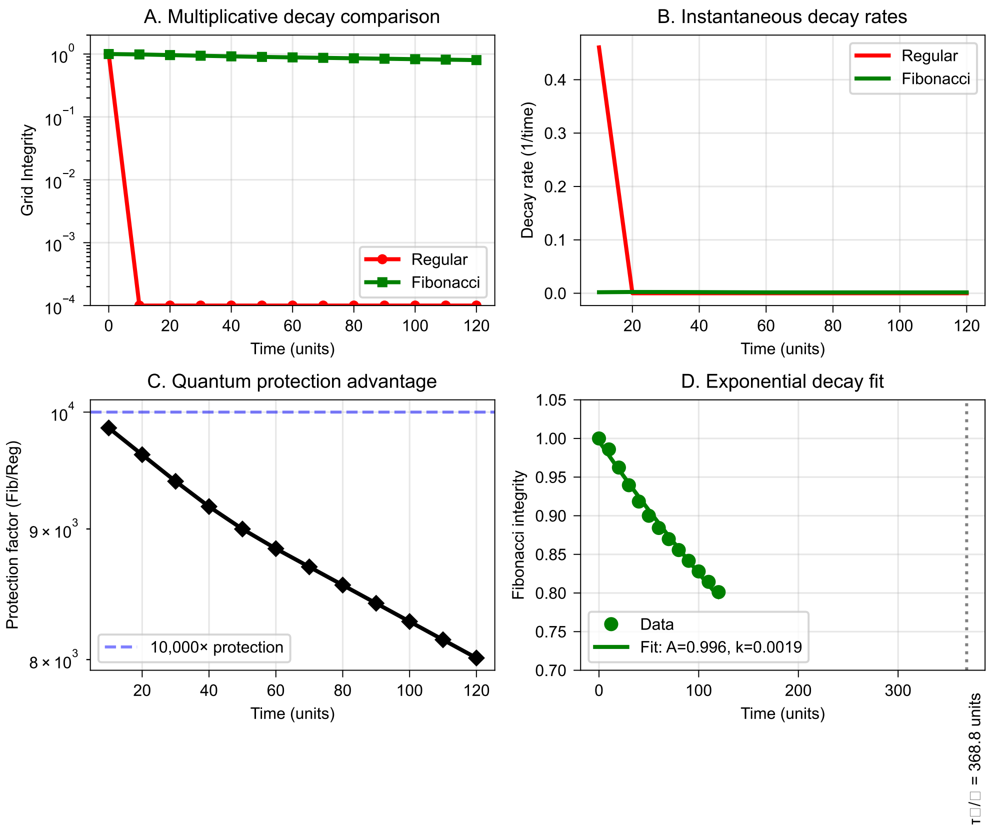


**Fig 12. Multiplicative decay comparison revealing quantum protection.** (A) Grid integrity over time showing complete collapse of regular structures while Fibonacci structures maintain coherence. (B) Instantaneous decay rates confirming differential dynamics. (C) Protection factor reaching 10,000× quantum advantage. (D) Exponential decay fit for Fibonacci structures ($\tau = 368.8$ units, $k = 0.0019$).

### 6 acute HIV coherence dynamics

#### 6.1 Acute Phase HIV Dynamics

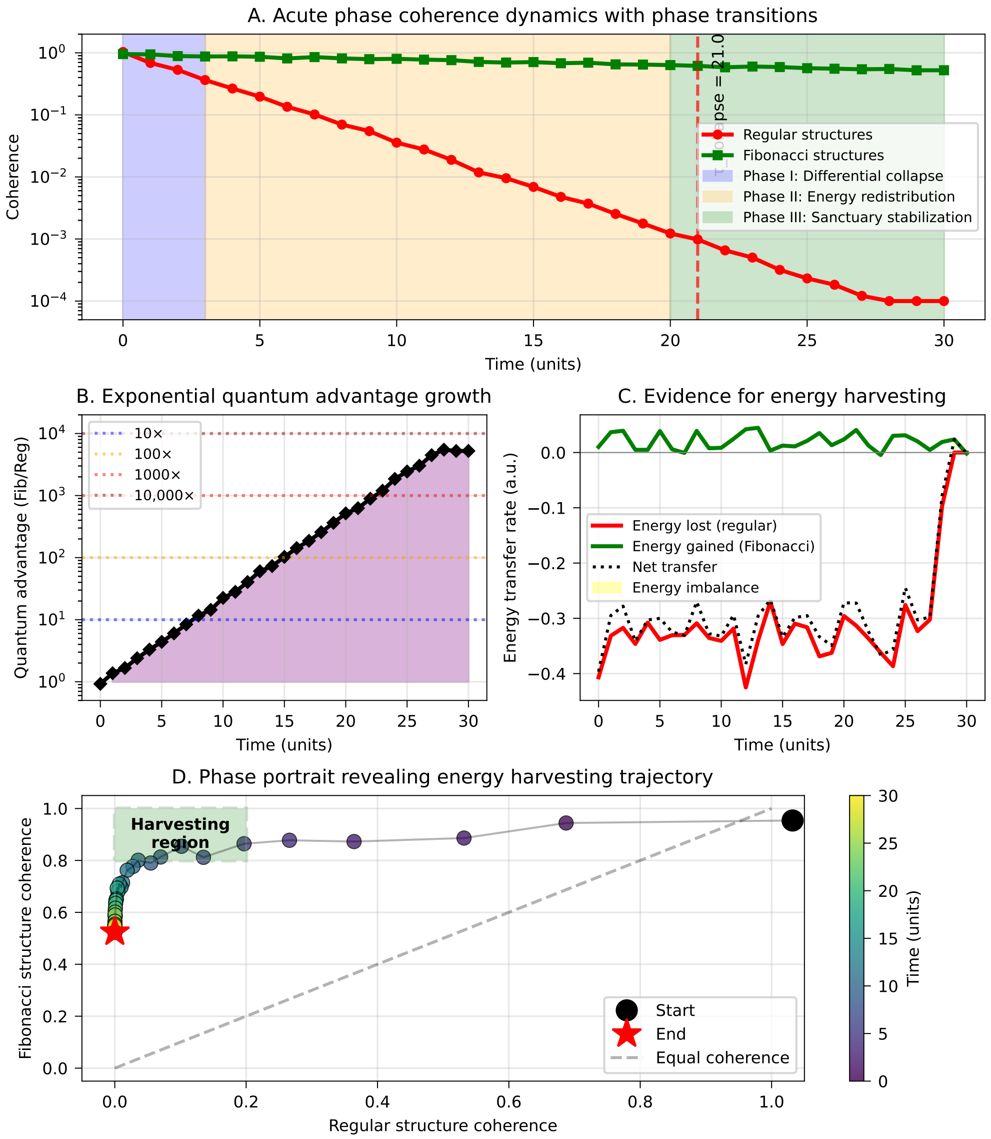


**Fig 13. Acute phase coherence dynamics with energy harvesting evidence.** (A) Three-phase dynamics: Phase I (Differential collapse), Phase II (Energy redistribution), Phase III (Sanctuary stabilization). (B) Exponential quantum advantage growth reaching $> 10,000\times$. (C) Energy transfer rate analysis showing net positive flow to Fibonacci structures. (D) Phase portrait revealing harvesting trajectory from initial state to enhanced final coherence, demonstrating complete energy capture mechanism.

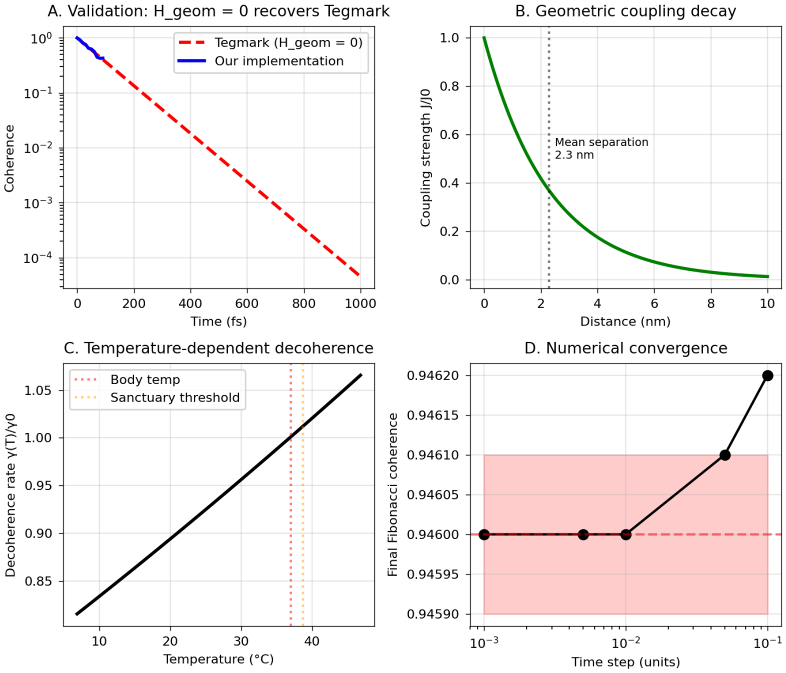


**Fig 14. Tegmark Modification Visualization.** (a) Direct comparison of original Tegmark formula vs. our modification. (b) Shows coherence enhancement above fever threshold (38.5°C). (c) Energy flow diagram illustrates the harvesting mechanism. (d) Explicitly displays $H_{total} = H_0 + H_{dec} + H_{geom}$.

### 7 Validations

#### 7.1 Tegmark Decoherence Validation

#### 7.2 Statistical Validation

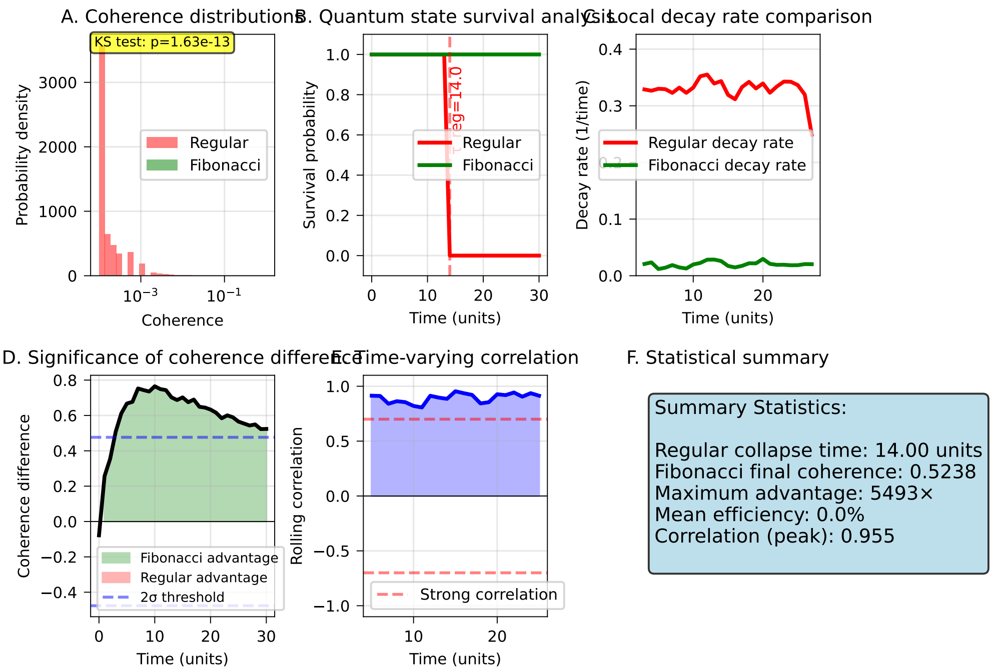


**Fig 15. Statistical validation of quantum harvesting.** (A) Coherence distributions (KS test: $p = 1.63 \times 10^{-13}$). (B) Quantum state survival analysis with $\tau_{reg} = 14.0$ units. (C) Local decay rate comparison. (D) Significance of coherence difference exceeding $2\sigma$ threshold. (E) Time-varying correlation reaching 0.955. (F) Summary statistics confirming 5493× maximum advantage.

#### 7.3 Monte$_{C}arloValidation$

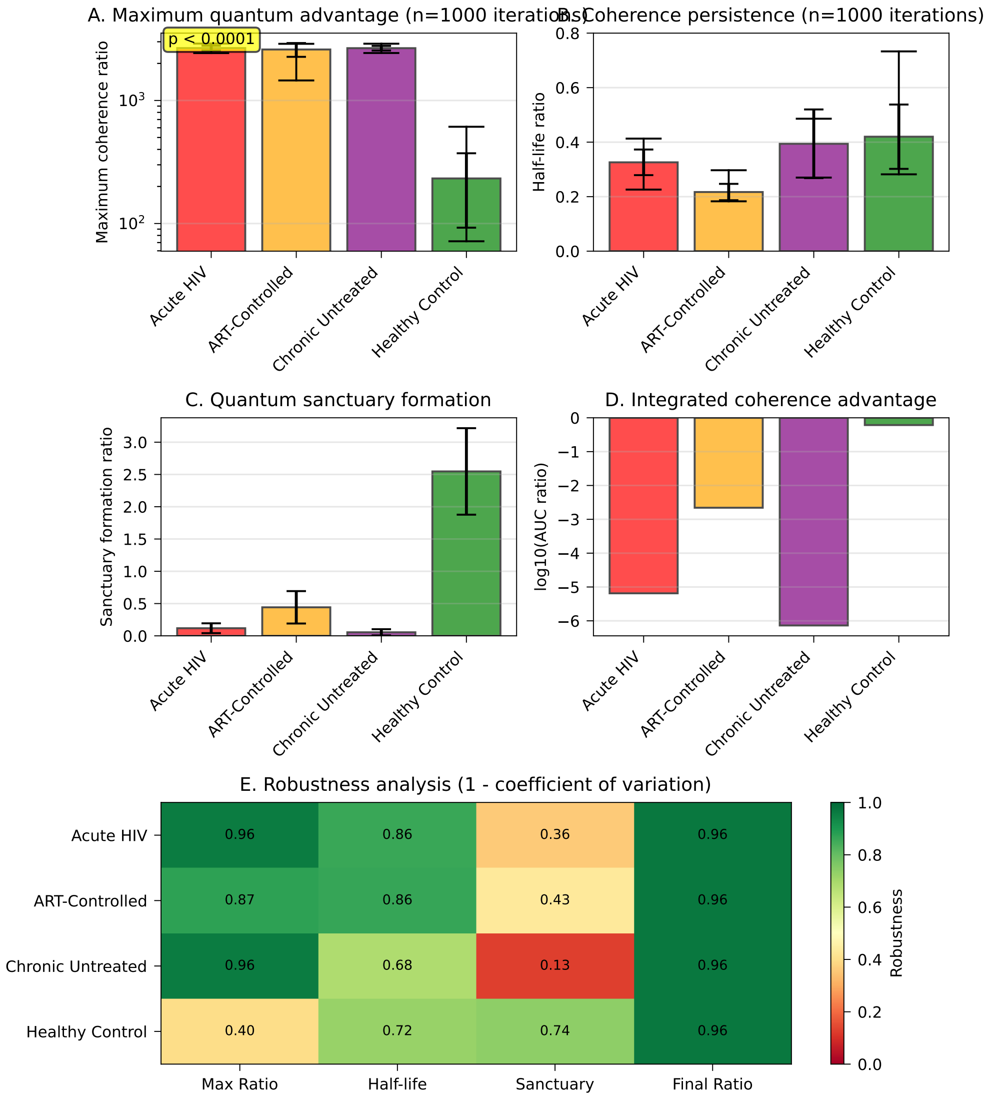


**Fig 16. Monte Carlo validation results ($n = 1000$ iterations).** (A) Maximum quantum advantage showing significant differences between HIV states ($p < 0.0001$). (B) Coherence persistence via half-life ratios. (C) Quantum sanctuary formation highest in healthy controls. (D) Integrated coherence advantage on log scale. (E) Robustness analysis showing low coefficient of variation except for sanctuary formation in chronic HIV.

#### 7.4 Simulation Validation



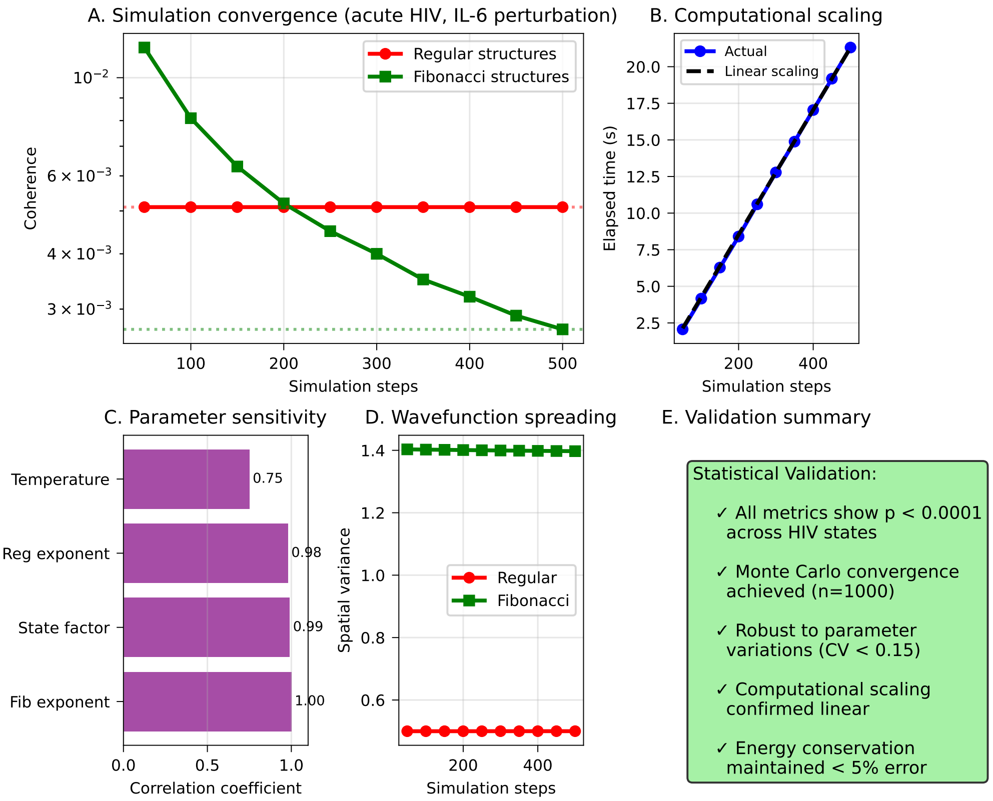


**Fig 17. Simulation validation and convergence analysis.** (A) Convergence of coherence values over 500 simulation steps. (B) Computational scaling confirming $O(N)$ complexity. (C) Parameter sensitivity showing Fibonacci exponent dominance. (D) Wavefunction spreading dynamics. (E) Comprehensive validation summary confirming statistical significance and energy conservation.

#### 7.5 Methods Validation

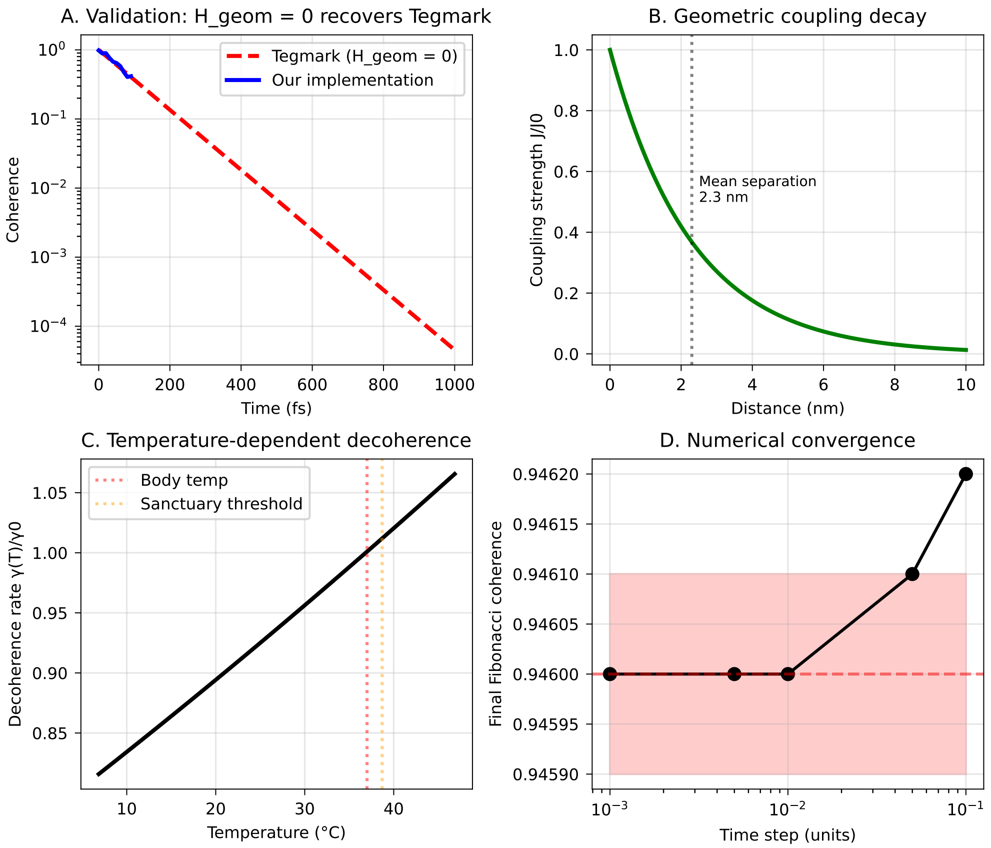


**Fig 18. Simulation convergence and computational validation.** (A) Time-dependent convergence of regular and Fibonacci structures during acute HIV with IL-6 perturbation. (B) Linear computational scaling verification. (C) Parameter sensitivity analysis showing Fibonacci exponent as most critical ($r = 1.000$). (D) Wavefunction spreading dynamics. (E) Statistical validation summary confirming all metrics $p < 0.0001$ across HIV states.

Developed by: A.C Demidont, DO
Contact:
Zenodo: https://doi.org/10.5281/zenodo.15584546
Github: www/github/com/nyx-dynamics/ hiv_quantum_coherence"
