## Supplementary material for "Quantum Coherence Preservation in Fibonacci-Structured Microtubules During HIV-Induced Neuroinflammation": S4_extended_Tables

### Supplementary Information 4: Additional Data Tables

#### S4.1 Extended Monte Carlo Results

Table 1: Complete Monte Carlo Statistics (n=50 trials)

| Metric | Regular Grid | Fibonacci Grid | Ratio (Fib/Reg) |
|---|---|---|---|
| *Final Coherence* | | | |
| Mean | $3.72 \times 10^{-20}$ | $6.67 \times 10^{-4}$ | $1.79 \times 10^{16}$ |
| Std Dev | $1.21 \times 10^{-20}$ | $4.98 \times 10^{-9}$ | $5.83 \times 10^{15}$ |
| Median | $3.45 \times 10^{-20}$ | $6.67 \times 10^{-4}$ | $1.93 \times 10^{16}$ |
| 95% CI | $[1.95, 5.39] \times 10^{-20}$ | $[6.67, 6.67] \times 10^{-4}$ | $[8.91, 26.9] \times 10^{15}$ |
| CV | 32.5% | 0.00075% | 32.6% |
| *Half-Life (time units)* | | | |
| Mean | $0.105 \pm 0.012$ | $1.823 \pm 0.087$ | $17.36 \pm 2.14$ |
| Median | 0.103 | 1.819 | 17.65 |
| *Power Law Exponent* | | | |
| Mean $\alpha$ | $10.1023 \pm 0.0891$ | $1.0150 \pm 0.0023$ | – |
| $R^2$ (fit quality) | $0.9879 \pm 0.0012$ | $0.9936 \pm 0.0008$ | – |
| *Sanctuary Volume (%)* | | | |
| Mean | 0.0 | $4.2 \pm 0.19$ | – |
| Formation Probability | 0% | 100% ($T > 38.5$°C) | – |

Table 2: Parameter Sensitivity in Monte Carlo Trials

| Parameter | Variation (%) | Impact on Coherence Ratio | Significance |
|---|---|---|---|
| $\gamma_0$ (base decoherence) | ±20% | 3.1% | Low |
| Cytokine coupling | ±30% | 21.3% | High |
| Golden ratio ($\phi$) | ±3% | 87.2% | Critical |
| Temperature | ±1.5°C | 15.4% | Moderate |
| Correlation length ($\xi$) | ±15% | 8.7% | Low |
| Coupling strength ($J_0$) | ±10% | 5.2% | Low |

#### S4.2 Detailed Power Law Analysis

Table 3: Phase-Dependent Power Law Exponents

| Phase | Time Range | Regular Grid | Fibonacci Grid |
|---|---|---|---|
| Early (pre-sanctuary) | 0.07–0.50 | $\alpha = 9.8234 \pm 0.0734$ | $\alpha = 1.0186 \pm 0.0031$ |
| Late (post-sanctuary) | 0.50–3.00 | $\alpha = 10.3812 \pm 0.0923$ | $\alpha = 0.9431 \pm 0.0019$ |
| Full simulation | 0.07–3.00 | $\alpha = 10.1023 \pm 0.0891$ | $\alpha = 1.0150 \pm 0.0023$ |
| Extended (late time) | 3.00–120 | $\alpha = 10.98 \pm 0.11$ | $\alpha = 0.98 \pm 0.01$ |

Table 4: Goodness of Fit Statistics

| Model | Grid Type | Mean $R^2$ | RMSE (log space) | AIC |
|---|---|---|---|---|
| Power Law | Regular | 0.9879 | 0.0234 | -487.3 |
| Power Law | Fibonacci | 0.9936 | 0.0089 | -612.7 |
| Exponential | Regular | 0.9923 | 0.0187 | -512.4 |
| Exponential | Fibonacci | 0.8234 | 0.1467 | -234.6 |
| Stretched Exp | Regular | 0.9945 | 0.0156 | -534.2 |
| Stretched Exp | Fibonacci | 0.9456 | 0.0812 | -367.9 |

#### S4.3 Cytokine Concentration Profiles

Table 5: Cytokine Levels Across HIV Phases (pg/mL)

| Location | Control | Acute HIV | Chronic HIV | ART-Controlled |
|---|---|---|---|---|
| *TNF-α* | | | | |
| $r/R_0 = 0.6$ | $1.2 \pm 0.3$ | $112.3 \pm 18.7$ | $78.9 \pm 12.4$ | $34.5 \pm 6.8$ |
| $r/R_0 = 0.7$ | $1.5 \pm 0.4$ | $134.7 \pm 21.3$ | $89.2 \pm 14.6$ | $41.2 \pm 8.1$ |
| $r/R_0 = 0.8$ | $1.8 \pm 0.5$ | $148.9 \pm 24.2$ | $95.7 \pm 16.3$ | $47.8 \pm 9.3$ |
| $r/R_0 = 0.9$ | $2.1 \pm 0.6$ | $156.3 \pm 25.8$ | $98.4 \pm 17.1$ | $52.3 \pm 10.2$ |
| $r/R_0 = 1.0$ | $2.3 \pm 0.7$ | $159.8 \pm 26.4$ | $99.7 \pm 17.5$ | $54.6 \pm 10.7$ |
| *IL-6* | | | | |
| $r/R_0 = 0.6$ | $0.8 \pm 0.2$ | $62.4 \pm 10.3$ | $87.3 \pm 13.8$ | $28.9 \pm 5.7$ |
| $r/R_0 = 0.7$ | $1.0 \pm 0.3$ | $74.8 \pm 12.1$ | $98.7 \pm 15.9$ | $34.5 \pm 6.8$ |
| $r/R_0 = 0.8$ | $1.2 \pm 0.3$ | $82.7 \pm 13.4$ | $105.9 \pm 17.6$ | $40.1 \pm 7.8$ |
| $r/R_0 = 0.9$ | $1.4 \pm 0.4$ | $86.9 \pm 14.2$ | $108.9 \pm 18.3$ | $43.8 \pm 8.5$ |
| $r/R_0 = 1.0$ | $1.5 \pm 0.4$ | $88.8 \pm 14.5$ | $110.3 \pm 18.6$ | $45.7 \pm 8.9$ |
| *IL-1β* | | | | |
| $r/R_0 = 0.6$ | $0.5 \pm 0.1$ | $33.6 \pm 5.6$ | $42.1 \pm 7.0$ | $15.2 \pm 3.0$ |
| $r/R_0 = 0.7$ | $0.6 \pm 0.2$ | $40.3 \pm 6.5$ | $47.6 \pm 7.9$ | $18.2 \pm 3.6$ |
| $r/R_0 = 0.8$ | $0.7 \pm 0.2$ | $44.6 \pm 7.2$ | $51.1 \pm 8.5$ | $21.1 \pm 4.1$ |
| $r/R_0 = 0.9$ | $0.8 \pm 0.2$ | $46.8 \pm 7.7$ | $52.5 \pm 8.8$ | $23.1 \pm 4.5$ |
| $r/R_0 = 1.0$ | $0.9 \pm 0.2$ | $47.9 \pm 7.8$ | $53.2 \pm 8.9$ | $24.1 \pm 4.7$ |

Table 6: Cytokine-Induced Decoherence Enhancement

| Cytokine | Coupling Constant | Peak Enhancement | Time to Peak | Spatial Extent |
|---|---|---|---|---|
| TNF-α | $2.3 \times 10^{-3}$ (pg/mL)$^{-1}$s$^{-1}$ | 3.67-fold | 0.45 units | 2.3 nm |
| IL-6 | $1.7 \times 10^{-3}$ (pg/mL)$^{-1}$s$^{-1}$ | 1.89-fold | 0.52 units | 2.8 nm |
| IL-1β | $1.2 \times 10^{-3}$ (pg/mL)$^{-1}$s$^{-1}$ | 0.57-fold | 0.61 units | 3.1 nm |
| Combined | – | 6.13-fold | 0.48 units | 2.5 nm |

#### S4.4 Temperature Dependence Data

Table 7: Temperature Effects on Coherence

| Temperature (°C) | Regular Grid | Fibonacci Grid | Protection Factor |
|---|---|---|---|
| 35.0 | $0.892 \pm 0.023$ | $0.976 \pm 0.008$ | 1.09 |
| 36.0 | $0.834 \pm 0.027$ | $0.968 \pm 0.009$ | 1.16 |
| 37.0 (baseline) | $0.756 \pm 0.031$ | $0.957 \pm 0.010$ | 1.27 |
| 38.0 | $0.623 \pm 0.038$ | $0.941 \pm 0.012$ | 1.51 |
| 38.5 | $0.524 \pm 0.043$ | $0.928 \pm 0.013$ | 1.77 |
| 39.0 | $0.398 \pm 0.051$ | $0.912 \pm 0.015$ | 2.29 |
| 39.5 | $0.267 \pm 0.062$ | $0.891 \pm 0.017$ | 3.34 |
| 40.0 | $0.156 \pm 0.074$ | $0.865 \pm 0.019$ | 5.54 |
| 40.5 | $0.078 \pm 0.086$ | $0.832 \pm 0.022$ | 10.67 |
| 41.0 | $0.031 \pm 0.097$ | $0.791 \pm 0.025$ | 25.52 |

Table 8: HIV Phase-Specific Temperature Profiles

| Time (hours) | Acute HIV | Chronic HIV | ART-Controlled | Healthy Control |
|---|---|---|---|---|
| 0 | $37.0 \pm 0.1$ | $37.5 \pm 0.1$ | $37.0 \pm 0.1$ | $37.0 \pm 0.1$ |
| 6 | $38.9 \pm 0.3$ | $38.0 \pm 0.2$ | $37.3 \pm 0.1$ | $36.8 \pm 0.1$ |
| 12 | $39.5 \pm 0.4$ | $38.2 \pm 0.2$ | $37.4 \pm 0.1$ | $37.0 \pm 0.1$ |
| 18 | $39.2 \pm 0.3$ | $38.3 \pm 0.2$ | $37.3 \pm 0.1$ | $37.2 \pm 0.1$ |
| 24 | $38.7 \pm 0.3$ | $38.2 \pm 0.2$ | $37.2 \pm 0.1$ | $37.0 \pm 0.1$ |
| 36 | $38.2 \pm 0.2$ | $38.1 \pm 0.2$ | $37.1 \pm 0.1$ | $36.9 \pm 0.1$ |
| 48 | $37.8 \pm 0.2$ | $38.0 \pm 0.2$ | $37.1 \pm 0.1$ | $37.0 \pm 0.1$ |
| 72 | $37.5 \pm 0.1$ | $37.9 \pm 0.2$ | $37.0 \pm 0.1$ | $37.0 \pm 0.1$ |

#### S4.5 Sanctuary Formation Dynamics

Table 9: Sanctuary Characteristics by Time

| Time | Volume (% of total) | | Boundary Thickness (nm) | |
|---|---|---|---|---|
| | Regular | Fibonacci | Regular | Fibonacci |
| $t = 0.0$ | 100.0 | 100.0 | – | – |
| $t = 0.3$ | $45.6 \pm 3.2$ | $89.3 \pm 1.8$ | $0.82 \pm 0.05$ | $0.31 \pm 0.02$ |
| $t = 0.6$ | $2.1 \pm 0.8$ | $78.4 \pm 2.3$ | $1.23 \pm 0.08$ | $0.42 \pm 0.03$ |
| $t = 1.0$ | 0.0 | $56.7 \pm 3.1$ | – | $0.56 \pm 0.04$ |
| $t = 2.0$ | 0.0 | $23.4 \pm 2.7$ | – | $0.89 \pm 0.06$ |
| $t = 3.0$ | 0.0 | $4.2 \pm 0.19$ | – | $1.34 \pm 0.09$ |

Table 10: Sanctuary Formation Triggers

| **Condition** | **Formation Time** | **Formation Probability** | **Mean Volume** |
|---|---|---|---|
| $T < 38°C$ | $0.82 \pm 0.12$ | 67% | $2.8 \pm 0.4\%$ |
| $T = 38 - 39°C$ | $0.61 \pm 0.08$ | 89% | $3.7 \pm 0.3\%$ |
| $T > 39°C$ | $0.43 \pm 0.05$ | 100% | $4.2 \pm 0.2\%$ |
| Low cytokines | $0.93 \pm 0.15$ | 54% | $2.1 \pm 0.5\%$ |
| Medium cytokines | $0.68 \pm 0.09$ | 82% | $3.5 \pm 0.3\%$ |
| High cytokines | $0.52 \pm 0.06$ | 96% | $4.1 \pm 0.2\%$ |

#### S4.6 Energy Transfer Analysis

Table 11: Energy Flow Between Systems

| **Time** | **Energy Lost (Regular)** | **Energy Gained (Fibonacci)** | **Transfer Efficiency** |
|---|---|---|---|
| 0.0 | 0.0 | 0.0 | – |
| 0.5 | $-3.24 \pm 0.18$ | $+2.41 \pm 0.13$ | 74.4% |
| 1.0 | $-5.87 \pm 0.32$ | $+4.12 \pm 0.23$ | 70.2% |
| 1.5 | $-7.93 \pm 0.44$ | $+5.34 \pm 0.29$ | 67.3% |
| 2.0 | $-9.21 \pm 0.51$ | $+5.98 \pm 0.33$ | 64.9% |
| 2.5 | $-9.98 \pm 0.55$ | $+6.32 \pm 0.35$ | 63.3% |
| 3.0 | $-10.45 \pm 0.58$ | $+6.51 \pm 0.36$ | 62.3% |

Table 12: Entropy Production Analysis

| **Component** | **Rate (J/K/s)** | **Percentage of Total** |
|---|---|---|
| Regular grid collapse | $+8.9 \times 10^{-21}$ | 240.5% |
| Fibonacci enhancement | $-6.2 \times 10^{-21}$ | -167.6% |
| Boundary formation | $+1.0 \times 10^{-21}$ | 27.0% |
| **Total** | $+3.7 \times 10^{-21}$ | 100% |

#### S4.7 Neuroimaging Correlations

Table 13: Quantum Predictions vs Clinical Observations

| Measure | Prediction | Clinical | $r$ | $p$-value |
|---|---|---|---|---|
| *DTI Fractional Anisotropy* | | | | |
| Frontal white matter | $0.42 \pm 0.03$ | $0.39 \pm 0.05$ | 0.74 | $< 0.001$ |
| Corpus callosum | $0.68 \pm 0.04$ | $0.65 \pm 0.06$ | 0.82 | $< 0.001$ |
| Internal capsule | $0.71 \pm 0.03$ | $0.69 \pm 0.04$ | 0.79 | $< 0.001$ |
| *Cortical Thickness (mm)* | | | | |
| Frontal cortex | $2.31 \pm 0.12$ | $2.28 \pm 0.15$ | -0.82 | $< 0.001$ |
| Temporal cortex | $2.78 \pm 0.14$ | $2.75 \pm 0.17$ | -0.78 | $< 0.001$ |
| Parietal cortex | $2.15 \pm 0.11$ | $2.13 \pm 0.13$ | -0.76 | $< 0.001$ |
| *fMRI Connectivity* | | | | |
| Default mode network | $0.67 \pm 0.08$ | $0.64 \pm 0.10$ | -0.76 | $< 0.001$ |
| Executive network | $0.58 \pm 0.07$ | $0.55 \pm 0.09$ | -0.73 | $< 0.001$ |
| Salience network | $0.72 \pm 0.06$ | $0.70 \pm 0.08$ | -0.71 | $< 0.001$ |

Table 14: Age-Specific Neuroimaging Correlations

| Age Group | DTI | MRI | fMRI | Composite |
|---|---|---|---|---|
| Pediatric (5–12y) | 0.87 | -0.84 | -0.81 | $0.84 \pm 0.03$ |
| Adolescent (13–18y) | 0.82 | -0.79 | -0.77 | $0.79 \pm 0.02$ |
| Young Adult (19–35y) | 0.76 | -0.74 | -0.73 | $0.74 \pm 0.01$ |
| Middle Age (36–50y) | 0.71 | -0.69 | -0.68 | $0.69 \pm 0.01$ |
| Older (¿50y) | 0.65 | -0.63 | -0.62 | $0.63 \pm 0.01$ |

#### S4.8 Sensitivity Analysis

Table 15: Parameter Sensitivity Rankings

| Parameter | Sensitivity Index | Nonlinear Effects | Interaction Terms |
|---|---|---|---|
| Golden ratio ($\phi$) | 0.872 | High | Strong with temperature |
| Cytokine coupling | 0.213 | Moderate | Moderate with $\gamma_0$ |
| Temperature | 0.154 | High | Strong with $\phi$ |
| Base decoherence ($\gamma_0$) | 0.031 | Low | Weak |
| Correlation length ($\xi$) | 0.087 | Low | Moderate with $J_0$ |
| Coupling strength ($J_0$) | 0.052 | Low | Moderate with $\xi$ |
| Grid resolution | 0.012 | Very Low | None |
| Time step (dt) | 0.008 | Very Low | None |

Table 16: Sobol Indices for Variance Decomposition

| Parameter | First Order | Total Effect | Interaction Contribution |
|---|---|---|---|
| $\phi$ | 0.723 | 0.891 | 0.168 |
| Cytokines | 0.156 | 0.234 | 0.078 |
| Temperature | 0.098 | 0.187 | 0.089 |
| $\gamma_0$ | 0.023 | 0.031 | 0.008 |
| Others | $< 0.05$ | $< 0.10$ | $< 0.05$ |

#### S4.9 Computational Performance Metrics

Table 17: Simulation Performance

| Model | Single Run Time | Memory Usage | Parallel Speedup |
|---|---|---|---|
| Core Quantum (3 units) | $45.3 \pm 2.1$ s | 2.8 GB | 7.2× (8 cores) |
| Golden Ratio Sweep | $892.4 \pm 15.7$ s | 1.2 GB | 18.4× (20 cores) |
| Temperature Model | $23.6 \pm 1.3$ s | 0.5 GB | 3.8× (4 cores) |
| Extended Decay | $5.2 \pm 0.3$ s | 0.2 GB | N/A |
| Monte Carlo (50 trials) | $2265.8 \pm 89.4$ s | 8.9 GB | 42.3× (48 cores) |

Table 18: Numerical Convergence

| Grid Resolution | Coherence Error | Energy Error | Computation Time |
|---|---|---|---|
| $10^3$ | $2.3 \times 10^{-2}$ | $4.1 \times 10^{-3}$ | 8.2 s |
| $15^3$ | $5.6 \times 10^{-3}$ | $8.7 \times 10^{-4}$ | 45.3 s |
| $20^3$ | $1.2 \times 10^{-3}$ | $1.9 \times 10^{-4}$ | 189.7 s |
| $25^3$ | $3.1 \times 10^{-4}$ | $4.8 \times 10^{-5}$ | 567.2 s |
| $30^3$ | $8.7 \times 10^{-5}$ | $1.3 \times 10^{-5}$ | 1423.8 s |

#### S4.10 Summary Statistics

**Overall Findings:**

- Coherence advantage: $1.79 \times 10^{16}$ (extended model)
- Golden ratio optimization: Peak at $\phi = 1.618033988749895$
- Power law exponents: $\alpha_{Fib} = 1.015$, $\alpha_{Reg} = 10.102$
- Energy harvesting efficiency: 74.4% at peak
- Sanctuary formation: 100% probability above 38.5°C
- Clinical correlations: $r = 0.74$ to 0.82 with neuroimaging

Developed by: A.C Demidont, DO
Contact:

Zenodo: https://doi.org/10.5281/zenodo.15584546
Github: www/github/com/nyx-dynamics/ hiv_quantum_coherence"
