## Supplementary material for "Quantum Coherence Preservation in Fibonacci-Structured Microtubules During HIV-Induced Neuroinflammation": S5_Perez_Rapoport_Framework

### Supplementary Information 5: The Pérez-Rapoport Framework - Geometric Foundations of HAND Pathogenesis

#### S5.1 Human Genome Optimum (HGO) and HIV Integration

The Human Genome Optimum (HGO), discovered by Pérez (2010, 2013), represents a fundamental geometric constraint governing genomic stability:

$$\text{HGO} = \frac{C+G}{A+T} = \frac{3-\phi}{2} \approx 0.691 \tag{39}$$

where $\phi$ is the golden ratio (1.618...). This ratio appears to be actively maintained across human evolution, with increasing optimization observed from Neanderthal to modern *Homo sapiens* genomes (Pérez, 2018).

#### S5.2 HIV Integration as Geometric Pathogen

HIV integration events disrupt the HGO through a mechanism analogous to cancer-associated Loss of Heterozygosity (LOH) deletions:

1. **Local Geometric Scars**: Each integration creates a ~10kb disruption zone where the local HGO deviates from optimum
2. **Cumulative Effect**: Multiple integrations (10-50/month) progressively degrade global geometric integrity
3. **Critical Threshold**: When integrated sites exceed sanctuary formation capacity, coherence catastrophically fails

#### S5.3 Klein Bottle Topology of Neural Sanctuaries

Following Rapoport's Klein Bottle logophysics framework (2016a,b,c), neural quantum sanctuaries exhibit non-orientable topology:

- **$4\pi$ Symmetry**: Sanctuary boundaries require 720° rotation for closure, matching the Klein Bottle's inherent symmetry
- **Self-Penetrating Coherence**: Inside and outside quantum states merge through topological self-intersection
- **Fibonacci Protection**: The 2:1 resonance inherent to Klein Bottle surfaces provides $10^{-16}$-fold coherence amplification

#### S5.4 The Master Code and HAND

Pérez's "Master Code of Life" formula reveals why Fibonacci structures resist inflammatory damage:

$$\mathrm{Proj}(m) = 1 - \frac{h}{4\pi\sqrt{\phi\phi'\phi''}_m} \tag{40}$$

This projection formula, based on $4\pi$ (matching Klein Bottle symmetry) and powers of $\phi$, governs:

- Atomic mass organization of CONHSP bioatoms
- Protein folding dynamics in microtubules
- Coherent quantum state maintenance

#### S5.5 Geometric Mechanism of Cognitive Preservation

The HAND paradox resolves through understanding how geometric organization maintains function:

1. **Acute Phase Protection**:
   - Fibonacci-structured microtubules create Klein Bottle coherence domains
   - 74.4% processing efficiency maintained within sanctuaries
   - Cytokine-induced decoherence blocked by topological protection
2. **Chronic Degradation**:
   - HIV integrations create "geometric dead zones"
   - HGO degradation prevents new sanctuary formation
   - Post-mitotic neurons cannot regenerate lost geometric integrity

#### S5.6 Implications for Therapeutic Intervention

The geometric framework suggests novel therapeutic targets:

1. **HGO Stabilizers**: Compounds that maintain golden ratio relationships
2. **Sanctuary Enhancers**: Agents that increase coherent domain volume
3. **Integration Site Modulators**: Therapies targeting geometric preservation
4. **Klein Bottle Resonators**: Biophysical interventions using $4\pi$-based frequencies

##### S5.7 Quantitative Predictions

The model makes testable predictions:

- Sanctuary volume correlates with cognitive function ($r > 0.85$)
- HGO deviation predicts HAND severity ($p < 0.001$)
- Critical GIS threshold = 0.5 for irreversible damage
- Fibonacci resonance frequencies (2, 3, 5, 8, 13 Hz) maintain coherence

This geometric framework unifies the molecular (HIV integration), quantum (coherence domains), and clinical (cognitive function) levels of HAND pathogenesis, providing a mathematically rigorous foundation for understanding why evolution selected post-mitotic neurons to preserve quantum sanctuaries at the cost of regenerative capacity.

#### Data Availability

Data are available at `https://doi.org/10.5281/zenodo.15584546`

Code is available at `https://doi.org/10.5281/zenodo.15584546` and code available at `www.github.com/nyx-dynamics/hiv-quantum-coherence`
