## Supplementary material for "Quantum Coherence Preservation in Fibonacci-Structured Microtubules During HIV-Induced Neuroinflammation": 5 model results: full_hiv_simulation.py

```
import numpy as np
import math
import random
import matplotlib.pyplot as plt
from matplotlib.figure import Figure
from mpl_toolkits.mplot3d import Axes3D
import scipy.spatial.distance as distance

# Define simulation parameters
params = {
    'max_time': 72,  # 72 time units (3 days when converted to hours)
    'output_interval': 0.1,  # Store results every 0.1 time units
    'initial_targeting': 0.05,  # Starting immune response intensity
    'hiv_phase': 'acute',  # Acute phase of HIV infection
    'grid_resolution': 20,  # 3D grid resolution (20x20x20 = 8000 nodes)
    'cell_radius': 10.0,  # Cell radius in micrometers
    'microtubule_count': 24  # Number of microtubules extending from centrosome
}


class Microtubule3D:
    """Represents a single microtubule in 3D space"""

    def __init__(self, direction_vector, length, geometry_type='regular'):
        self.direction = np.array(direction_vector) / np.linalg.norm(direction_vector)
        self.length = length
        self.geometry_type = geometry_type
        self.nodes = self.generate_nodes()

    def generate_nodes(self):
        """Generate 3D nodes along the microtubule"""
        nodes = []
        if self.geometry_type == 'fibonacci':
            # Fibonacci spiral pattern along microtubule
            golden_ratio = (1 + math.sqrt(5)) / 2
            num_nodes = int(self.length)

            for i in range(num_nodes):
                t = i / num_nodes
                # Fibonacci spiral around the central axis
                theta = 2 * math.pi * i / golden_ratio
                radius_offset = 0.5 * math.sin(math.pi * t)

                # Create perpendicular vectors for spiral
                if abs(self.direction[2]) < 0.9:
                    perp1 = np.cross(self.direction, [0, 0, 1])
                else:
                    perp1 = np.cross(self.direction, [1, 0, 0])
                perp1 = perp1 / np.linalg.norm(perp1)
                perp2 = np.cross(self.direction, perp1)

                # Position along microtubule with spiral
                position = (t * self.length * self.direction +
                            radius_offset * (math.cos(theta) * perp1 + math.sin(theta) * perp2))

                nodes.append({
                    'position': position,
                    'integrity': 1.0,
                    'coherence': 1.0,
                    'local_temperature': 37.0
                })
        else:
            # Regular linear pattern
            num_nodes = int(self.length)
            for i in range(num_nodes):
                t = i / num_nodes
                position = t * self.length * self.direction
                nodes.append({
                    'position': position,
                    'integrity': 1.0,
                    'coherence': 1.0,
                    'local_temperature': 37.0
                })

        return nodes


class Spatial3DGrid:
    """3D spatial grid for quantum coherence simulation"""

    def __init__(self, resolution, cell_radius, geometry_type='regular'):
        self.resolution = resolution
        self.cell_radius = cell_radius
        self.geometry_type = geometry_type
        self.grid_spacing = 2 * cell_radius / resolution

        # Initialize 3D grid
        self.grid = np.zeros((resolution, resolution, resolution), dtype=object)
        self.coherence_field = np.ones((resolution, resolution, resolution))
        self.integrity_field = np.ones((resolution, resolution, resolution))
        self.temperature_field = np.full((resolution, resolution, resolution), 37.0)

        # Create microtubule network
        self.microtubules = self.create_microtubule_network()
        self.populate_grid()

    def create_microtubule_network(self):
        """Create realistic 3D microtubule arrangement from centrosome"""
        microtubules = []
        centrosome = np.array([0, 0, 0])  # Center of cell

        # Generate microtubules in 3D directions
        num_microtubules = params.get('microtubule_count', 24)

        for i in range(num_microtubules):
            # Spherical coordinates for even distribution
            theta = 2 * math.pi * i / num_microtubules
            phi = math.acos(1 - 2 * (i + 0.5) / num_microtubules)

            # Convert to Cartesian coordinates
            direction = np.array([
                math.sin(phi) * math.cos(theta),
                math.sin(phi) * math.sin(theta),
                math.cos(phi)
            ])

            # Microtubule length (varies slightly)
            length = self.cell_radius * (0.8 + 0.4 * random.random())

            # Create microtubule
            microtubule = Microtubule3D(direction, length, self.geometry_type)
            microtubules.append(microtubule)

        return microtubules

    def populate_grid(self):
        """Populate 3D grid with microtubule data"""
        center_idx = self.resolution // 2

        # Initialize all grid positions to None
        for x in range(self.resolution):
            for y in range(self.resolution):
                for z in range(self.resolution):
                    self.grid[x, y, z] = None

        for microtubule in self.microtubules:
            for node in microtubule.nodes:
                # Convert world coordinates to grid indices
                world_pos = node['position']
                grid_pos = world_pos / self.grid_spacing + center_idx

                # Check bounds
                x, y, z = int(grid_pos[0]), int(grid_pos[1]), int(grid_pos[2])
                if (0 <= x < self.resolution and
                        0 <= y < self.resolution and
                        0 <= z < self.resolution):
                    # Create node data structure
                    self.grid[x, y, z] = {
                        'microtubule_type': microtubule.geometry_type,
                        'integrity': node['integrity'],
                        'coherence': node['coherence'],
                        'temperature': node['local_temperature'],
                        'neighbors': self.get_neighbor_indices(x, y, z)
                    }

    def get_neighbor_indices(self, x, y, z):
        """Get neighboring grid indices for spatial effects"""
        neighbors = []
        for dx in [-1, 0, 1]:
            for dy in [-1, 0, 1]:
                for dz in [-1, 0, 1]:
                    if dx == 0 and dy == 0 and dz == 0:
                        continue
                    nx, ny, nz = x + dx, y + dy, z + dz
                    if (0 <= nx < self.resolution and
                            0 <= ny < self.resolution and
                            0 <= nz < self.resolution):
                        neighbors.append((nx, ny, nz))
        return neighbors

    def update_spatial_coherence(self, dt, temperature_field):
        """Update coherence with 3D spatial diffusion effects"""
        new_coherence = self.coherence_field.copy()
        new_integrity = self.integrity_field.copy()

        for x in range(self.resolution):
            for y in range(self.resolution):
                for z in range(self.resolution):
                    if self.grid[x, y, z] is not None and isinstance(self.grid[x, y, z], dict):
                        node = self.grid[x, y, z]
                        temp = temperature_field[x, y, z]

                        # Temperature effects on decoherence
                        temp_diff = temp - 37.0

                        if node['microtubule_type'] == 'fibonacci':
                            # Fibonacci structure: power-law decay with temperature protection
                            base_decay_rate = 0.005
                            temp_sensitivity = 0.05
                            geometry_protection = 1.618 ** -2.5  # Golden ratio protection
                        else:
                            # Regular structure: exponential decay with high temperature sensitivity
                            base_decay_rate = 0.8
                            temp_sensitivity = 0.3
                            geometry_protection = 1.0

                        # Calculate decay rate
                        decay_rate = base_decay_rate * math.exp(temp_sensitivity * max(0, temp_diff))
                        decay_rate *= geometry_protection

                        # Apply spatial diffusion from neighbors
                        neighbor_coherence = 0
                        neighbor_count = 0

                        for nx, ny, nz in node['neighbors']:
                            neighbor_coherence += self.coherence_field[nx, ny, nz]
                            neighbor_count += 1

                        if neighbor_count > 0:
                            avg_neighbor_coherence = neighbor_coherence / neighbor_count
                            spatial_diffusion = 0.01 * (avg_neighbor_coherence - self.coherence_field[x, y, z])
                        else:
                            spatial_diffusion = 0

                        # Update coherence and integrity
                        new_coherence[x, y, z] = max(1e-10,
                                                     self.coherence_field[x, y, z] * (
                                                                 1 - decay_rate * dt) + spatial_diffusion * dt)

                        new_integrity[x, y, z] = max(0.0001,
                                                     self.integrity_field[x, y, z] * (1 - decay_rate * dt * 0.5))
                    else:
                        # For empty grid positions, apply general temperature decay
                        temp = temperature_field[x, y, z]
                        temp_diff = temp - 37.0

                        # Use geometry type from the grid object
                        if self.geometry_type == 'fibonacci':
                            base_decay_rate = 0.005
                            temp_sensitivity = 0.05
                        else:
                            base_decay_rate = 0.8
                            temp_sensitivity = 0.3

                        decay_rate = base_decay_rate * math.exp(temp_sensitivity * max(0, temp_diff))

                        # Apply mild decay to empty regions
                        new_coherence[x, y, z] = max(1e-10,
                                                     self.coherence_field[x, y, z] * (1 - decay_rate * dt * 0.1))
                        new_integrity[x, y, z] = max(0.0001,
                                                     self.integrity_field[x, y, z] * (1 - decay_rate * dt * 0.05))

        self.coherence_field = new_coherence
        self.integrity_field = new_integrity

    def calculate_sanctuary_regions(self, temperature_field, sanctuary_threshold=38.5):
        """Identify and enhance quantum sanctuary regions in 3D"""
        sanctuary_mask = np.zeros((self.resolution, self.resolution, self.resolution), dtype=bool)

        # Find high-temperature, high-coherence regions for Fibonacci structures
        if self.geometry_type == 'fibonacci':
            for x in range(self.resolution):
                for y in range(self.resolution):
                    for z in range(self.resolution):
                        if (self.grid[x, y, z] is not None and
                                isinstance(self.grid[x, y, z], dict) and
                                temperature_field[x, y, z] > sanctuary_threshold and
                                self.integrity_field[x, y, z] > 0.1):

                            # Check if region has sufficient coherence density
                            local_coherence = self.coherence_field[x, y, z]
                            neighbor_coherence = 0
                            neighbor_count = 0

                            node = self.grid[x, y, z]
                            for nx, ny, nz in node['neighbors']:
                                if self.grid[nx, ny, nz] is not None and isinstance(self.grid[nx, ny, nz], dict):
                                    neighbor_coherence += self.coherence_field[nx, ny, nz]
                                    neighbor_count += 1

                            if neighbor_count > 0:
                                avg_coherence = (local_coherence + neighbor_coherence / neighbor_count) / 2
                                if avg_coherence > 0.05:  # Sufficient coherence for sanctuary
                                    sanctuary_mask[x, y, z] = True

                                    # Enhance coherence in sanctuary regions
                                    enhancement_factor = 1.5 + 0.5 * (temperature_field[x, y, z] - sanctuary_threshold)
                                    self.coherence_field[x, y, z] *= enhancement_factor

        return sanctuary_mask

    def get_global_metrics(self):
        """Calculate overall grid metrics"""
        total_integrity = np.mean(self.integrity_field[self.integrity_field > 0.0001])
        total_coherence = np.mean(self.coherence_field[self.coherence_field > 1e-10])

        # Separate metrics by geometry type
        fibonacci_integrity_values = []
        fibonacci_coherence_values = []
        regular_integrity_values = []
        regular_coherence_values = []

        for x in range(self.resolution):
            for y in range(self.resolution):
                for z in range(self.resolution):
                    if self.grid[x, y, z] is not None and isinstance(self.grid[x, y, z], dict):
                        integrity_val = self.integrity_field[x, y, z]
                        coherence_val = self.coherence_field[x, y, z]

                        if self.grid[x, y, z]['microtubule_type'] == 'fibonacci':
                            fibonacci_integrity_values.append(integrity_val)
                            fibonacci_coherence_values.append(coherence_val)
                        else:
                            regular_integrity_values.append(integrity_val)
                            regular_coherence_values.append(coherence_val)

        # Calculate averages, defaulting to grid-wide averages if no specific geometry found
        if fibonacci_integrity_values:
            fib_integrity = np.mean(fibonacci_integrity_values)
            fib_coherence = np.mean(fibonacci_coherence_values)
        else:
            # Use overall grid values if this is a fibonacci grid but no nodes found
            fib_integrity = total_integrity if self.geometry_type == 'fibonacci' else 0
            fib_coherence = total_coherence if self.geometry_type == 'fibonacci' else 0

        if regular_integrity_values:
            reg_integrity = np.mean(regular_integrity_values)
            reg_coherence = np.mean(regular_coherence_values)
        else:
            # Use overall grid values if this is a regular grid but no nodes found
            reg_integrity = total_integrity if self.geometry_type == 'regular' else 0
            reg_coherence = total_coherence if self.geometry_type == 'regular' else 0

        return {
            'total_integrity': total_integrity,
            'total_coherence': total_coherence,
            'fibonacci_integrity': fib_integrity,
            'fibonacci_coherence': fib_coherence,
            'regular_integrity': reg_integrity,
            'regular_coherence': reg_coherence
        }


def calculate_decoherence_rate(temperature, geometry, scaling):
    """
    Calculate temperature-dependent quantum decoherence rate.
    """
    # Base parameters
    h_bar = 1.054571817e-34  # Reduced Planck constant
    k_b = 1.380649e-23  # Boltzmann constant
    r = 3.8e-9  # Superposition distance

    # Temperature-dependent dielectric constant (corrected from Tegmark)
    epsilon = 80 * ((310 / temperature) ** 1.5)

    # Geometry-specific factor (critical for Fibonacci protection)
    if geometry == 'fibonacci':
        geometry_factor = scaling ** -2.5  # Power law protection
    else:
        geometry_factor = 10.0 ** 9.1  # Exponential collapse factor

    # Calculate decoherence rate (inverse of coherence time)
    raw_rate = (k_b * temperature * 1.2e-29) / (h_bar * h_bar * epsilon * (r ** 4) * geometry_factor)

    return raw_rate


def simulate_hiv_fever(time_in_hours, phase='acute'):
    """
    Model the fever dynamics specific to HIV infection.
    """
    base_temp = 37.0  # Normal body temperature
    hour_of_day = time_in_hours % 24
    day_number = math.floor(time_in_hours / 24) + 1

    # Fever amplitude (varies by phase)
    if phase == 'acute':
        fever_amplitude = 2.5  # High fever in acute phase
        decay_rate = 0.2  # Per day
    elif phase == 'art-treated':
        fever_amplitude = 1.2  # Lower fever with treatment
        decay_rate = 0.4  # Faster resolution
    elif phase == 'chronic-untreated':
        fever_amplitude = 1.8  # Moderate persistent fever
        decay_rate = 0.05  # Very slow resolution
    else:
        fever_amplitude = 2.5
        decay_rate = 0.2

    # Day-dependent fever (decays exponentially)
    fever_component = fever_amplitude * math.exp(-decay_rate * (day_number - 1))

    # Circadian rhythm (higher in evening)
    circadian_component = 0.4 * math.sin(2 * math.pi * (hour_of_day / 24) - math.pi / 3)

    # Calculate temperature
    temperature = base_temp + fever_component + circadian_component

    # Add night sweats (between 2-4 AM in acute phase)
    if phase == 'acute' and 2 <= hour_of_day <= 4:
        minute_of_hour = (time_in_hours * 60) % 60
        # Create 2-3 sweating episodes per night
        sweat_episode = math.sin(2 * math.pi * minute_of_hour / 30) > 0.5

        if sweat_episode:
            # Rapid temperature drop during night sweats
            temperature -= 1.5 * math.sin(math.pi * minute_of_hour / 15)

    return temperature


def create_3d_temperature_field(base_temperature, grid_resolution, spatial_variation=True):
    """Create 3D temperature field with spatial variations"""
    temp_field = np.full((grid_resolution, grid_resolution, grid_resolution), base_temperature)

    if spatial_variation:
        # Add spatial temperature gradients (cell center is warmer)
        center = grid_resolution // 2
        for x in range(grid_resolution):
            for y in range(grid_resolution):
                for z in range(grid_resolution):
                    # Distance from center
                    distance_from_center = math.sqrt((x - center) ** 2 + (y - center) ** 2 + (z - center) ** 2)
                    max_distance = math.sqrt(3) * center

                    # Temperature gradient (center is warmer)
                    temp_gradient = 0.5 * (1 - distance_from_center / max_distance)
                    temp_field[x, y, z] += temp_gradient

                    # Add small random variations
                    temp_field[x, y, z] += random.gauss(0, 0.1)

    return temp_field


def update_grids_3d(state, dt, current_time):
    """
    Update 3D grid evolution incorporating temperature effects and quantum decoherence.
    """
    # Get current temperature from HIV fever model
    hour_time = current_time * 24  # Convert to hours
    base_temperature = simulate_hiv_fever(hour_time, state['hiv_phase'])

    # Create 3D temperature field
    temp_field = create_3d_temperature_field(base_temperature, state['regular_grid_3d'].resolution)

    # Update both grid systems
    state['regular_grid_3d'].temperature_field = temp_field
    state['fibonacci_grid_3d'].temperature_field = temp_field

    # Update spatial coherence for both grids
    state['regular_grid_3d'].update_spatial_coherence(dt, temp_field)
    state['fibonacci_grid_3d'].update_spatial_coherence(dt, temp_field)

    # Calculate sanctuary regions for Fibonacci grid
    sanctuary_mask = state['fibonacci_grid_3d'].calculate_sanctuary_regions(temp_field)

    # Update defense mechanisms based on 3D metrics
    fib_metrics = state['fibonacci_grid_3d'].get_global_metrics()
    reg_metrics = state['regular_grid_3d'].get_global_metrics()

    # Update state with 3D-derived metrics
    state['regular_grid']['integrity'] = reg_metrics['regular_integrity']
    state['regular_grid']['coherence'] = reg_metrics['regular_coherence']
    state['fibonacci_grid']['integrity'] = fib_metrics['fibonacci_integrity']
    state['fibonacci_grid']['coherence'] = fib_metrics['fibonacci_coherence']

    # Update legacy resonance calculations
    temp_diff = base_temperature - 37.0
    state['regular_grid']['resonance'] = state['regular_grid']['integrity'] * math.exp(-0.02 * temp_diff)
    state['fibonacci_grid']['resonance'] = state['fibonacci_grid']['integrity'] * math.exp(-0.01 * temp_diff)

    # Update defense mechanisms
    update_defense_mechanisms(state, base_temperature)

    # Track temperature and sanctuary info
    state['temperature'] = base_temperature
    state['temperature_history'].append(base_temperature)
    state['sanctuary_volume'] = np.sum(sanctuary_mask) / sanctuary_mask.size


def calculate_quantum_coherence(time, geometry, temperature):
    """
    Implement key finding from Tegmark's paper showing different decay patterns.
    """
    # Initial coherence values
    initial_coherence = 1.0

    # Temperature impact factor
    if geometry == 'fibonacci':
        temp_factor = math.exp(-0.05 * (temperature - 37.0))
    else:
        temp_factor = math.exp(-0.3 * (temperature - 37.0))

    # Coherence decay calculation
    if geometry == 'fibonacci':
        # Power-law decay for Fibonacci systems: t^(-1.015)
        coherence = initial_coherence * 0.015963 * (time ** -1.015) * temp_factor

        # Boundary/sanctuary formation effect at t=0.6
        if time > 0.6:
            coherence *= 1.6  # Sanctuary protection factor
    else:
        # Near-exponential decay for regular systems: t^(-10.1)
        coherence = initial_coherence * (time ** -10.1023) * temp_factor

    return max(coherence, 1e-20)


def calculate_information_sanctuary(state, time, temperature):
    """
    Model the formation of quantum-protected regions during acute fever stress.
    """
    # Sanctuary only forms under specific conditions
    sanctuary_threshold = 38.5  # Temperature threshold

    # Use 3D sanctuary calculations from Fibonacci grid
    if hasattr(state, 'sanctuary_volume') and state['sanctuary_volume'] > 0.01:
        sanctuary_strength = min(1.0, (temperature - sanctuary_threshold) / 1.5)
        state['fibonacci_grid']['sanctuary_strength'] = sanctuary_strength
        return True

    state['fibonacci_grid']['sanctuary_strength'] = 0
    return False


def update_defense_mechanisms(state, temperature):
    """
    Modify the defense mechanism activation to incorporate fever effects.
    """
    # Temperature-dependent activation threshold adjustment
    temp_diff = temperature - 37.0
    activation_boost = min(1.0, max(0, temp_diff / 3.0))

    # Phi Stability Defense activation
    if state['targeting_intensity'] > 0.45 - 0.15 * activation_boost:
        state['defenses']['phi_stability'] = min(1.0,
                                                 state['defenses']['phi_stability'] + 0.016 * (
                                                         1 + 0.5 * activation_boost))

    # Resonance Defense
    if state['targeting_intensity'] > 0.6 - 0.2 * activation_boost:
        state['defenses']['resonance'] = min(1.0,
                                             state['defenses']['resonance'] + 0.0165 * (1 + 0.3 * activation_boost))

    # Acetylation Defense
    if state['targeting_intensity'] > 0.7 - 0.1 * activation_boost:
        acetyl_rate = 0.0168 * (1.1 ** temp_diff)
        state['defenses']['acetylation'] = min(1.0,
                                               state['defenses']['acetylation'] + acetyl_rate)

    # Binding Sites Defense
    if state['targeting_intensity'] > 0.55 - 0.15 * activation_boost:
        state['defenses']['binding_sites'] = min(1.0,
                                                 state['defenses']['binding_sites'] + 0.0169 * (
                                                         1 + 0.2 * activation_boost))

    # Adaptation Defense
    if state['targeting_intensity'] > 0.65 - 0.2 * activation_boost and temperature > 38.0:
        state['defenses']['adaptation'] = min(1.0,
                                              state['defenses']['adaptation'] + 0.0168 * (1 + 0.4 * activation_boost))


def create_3d_visualizations(state):
    """
    Create 3D visualizations of the spatial grids.
    """
    figures = {}

    # 3D Coherence field visualization
    fig = plt.figure(figsize=(15, 5))

    # Regular grid coherence
    ax1 = fig.add_subplot(131, projection='3d')
    reg_grid = state['regular_grid_3d']

    # Sample points for visualization (too many points slow down rendering)
    x_coords, y_coords, z_coords = [], [], []
    coherence_values = []

    sample_count = 0
    for x in range(0, reg_grid.resolution, 2):  # Sample every 2nd point
        for y in range(0, reg_grid.resolution, 2):
            for z in range(0, reg_grid.resolution, 2):
                if reg_grid.grid[x, y, z] is not None and isinstance(reg_grid.grid[x, y, z], dict):
                    x_coords.append(x)
                    y_coords.append(y)
                    z_coords.append(z)
                    coherence_values.append(reg_grid.coherence_field[x, y, z])
                    sample_count += 1
                    if sample_count > 500:  # Limit for performance
                        break
                if sample_count > 500:
                    break
            if sample_count > 500:
                break

    if coherence_values:
        scatter = ax1.scatter(x_coords, y_coords, z_coords,
                              c=coherence_values, cmap='viridis',
                              s=20, alpha=0.6)
        ax1.set_title('Regular Grid Coherence')
        plt.colorbar(scatter, ax=ax1, shrink=0.5)
    else:
        ax1.text(0.5, 0.5, 0.5, 'No data points', transform=ax1.transAxes)
        ax1.set_title('Regular Grid Coherence (No Data)')

    # Fibonacci grid coherence
    ax2 = fig.add_subplot(132, projection='3d')
    fib_grid = state['fibonacci_grid_3d']

    x_coords, y_coords, z_coords = [], [], []
    coherence_values = []

    sample_count = 0
    for x in range(0, fib_grid.resolution, 2):  # Sample every 2nd point
        for y in range(0, fib_grid.resolution, 2):
            for z in range(0, fib_grid.resolution, 2):
                if fib_grid.grid[x, y, z] is not None and isinstance(fib_grid.grid[x, y, z], dict):
                    x_coords.append(x)
                    y_coords.append(y)
                    z_coords.append(z)
                    coherence_values.append(fib_grid.coherence_field[x, y, z])
                    sample_count += 1
                    if sample_count > 500:  # Limit for performance
                        break
                if sample_count > 500:
                    break
            if sample_count > 500:
                break

    if coherence_values:
        scatter = ax2.scatter(x_coords, y_coords, z_coords,
                              c=coherence_values, cmap='plasma',
                              s=20, alpha=0.6)
        ax2.set_title('Fibonacci Grid Coherence')
        plt.colorbar(scatter, ax=ax2, shrink=0.5)
    else:
        ax2.text(0.5, 0.5, 0.5, 'No data points', transform=ax2.transAxes)
        ax2.set_title('Fibonacci Grid Coherence (No Data)')

    # Temperature field
    ax3 = fig.add_subplot(133, projection='3d')
    temp_field = fib_grid.temperature_field

    x_coords, y_coords, z_coords = [], [], []
    temp_values = []

    # Sample temperature field more sparsely
    for x in range(0, fib_grid.resolution, 3):
        for y in range(0, fib_grid.resolution, 3):
            for z in range(0, fib_grid.resolution, 3):
                x_coords.append(x)
                y_coords.append(y)
                z_coords.append(z)
                temp_values.append(temp_field[x, y, z])

    if temp_values:
        scatter = ax3.scatter(x_coords, y_coords, z_coords,
                              c=temp_values, cmap='coolwarm',
                              s=20, alpha=0.6)
        ax3.set_title('Temperature Field')
        plt.colorbar(scatter, ax=ax3, shrink=0.5)

    plt.tight_layout()
    figures['3d_spatial'] = fig

    return figures


def update_targeting_intensity(state, t, dt):
    """
    Update targeting intensity based on simulation phase.
    """
    if t < 10:
        # Initial attack phase
        state['targeting_intensity'] = min(0.95, state['targeting_intensity'] + 0.1 * dt)
    elif 30 < t < 60:
        # Reduced targeting phase
        state['targeting_intensity'] = max(0.3, state['targeting_intensity'] - 0.01 * dt)
    else:
        # Maintenance phase
        state['targeting_intensity'] += 0.001 * dt * (random.random() - 0.5)
        state['targeting_intensity'] = max(0.3, min(0.95, state['targeting_intensity']))

    state['targeting_intensity'] = max(0.0, min(1.0, state['targeting_intensity']))


def initialize_system(params):
    """
    Initialize the simulation system with 3D spatial grids.
    """
    grid_resolution = params.get('grid_resolution', 20)
    cell_radius = params.get('cell_radius', 10.0)

    state = {
        # Legacy 2D grid data for compatibility
        'regular_grid': {
            'integrity': 1.0,
            'resonance': 1.0,
            'coherence': 1.0
        },
        'fibonacci_grid': {
            'integrity': 1.0,
            'resonance': 1.0,
            'coherence': 1.0,
            'sanctuary_radius': 0,
            'sanctuary_strength': 0
        },

        # New 3D spatial grids
        'regular_grid_3d': Spatial3DGrid(grid_resolution, cell_radius, 'regular'),
        'fibonacci_grid_3d': Spatial3DGrid(grid_resolution, cell_radius, 'fibonacci'),

        'defenses': {
            'phi_stability': 0.0,
            'resonance': 0.0,
            'acetylation': 0.0,
            'binding_sites': 0.0,
            'adaptation': 0.0
        },
        'targeting_intensity': params.get('initial_targeting', 0.05),
        'outer_radius': params.get('outer_radius', 1.0),
        'temperature_history': [],
        'hiv_phase': params.get('hiv_phase', 'acute'),
        'sanctuary_volume': 0.0
    }
    return state


def run_full_hiv_simulation(params):
    """
    Run the full multi-scale 3D simulation integrating all components.
    """
    # Initialize system state
    state = initialize_system(params)

    # Time parameters
    dt = 0.01
    max_time = params.get('max_time', 100)

    # Results storage
    results = []

    print(f"Initializing 3D simulation with {params.get('grid_resolution', 20)}^3 grid...")
    print(f"Regular grid integrity: {state['regular_grid']['integrity']}")
    print(f"Fibonacci grid integrity: {state['fibonacci_grid']['integrity']}")

    # Store initial state
    results.append({
        'time': 0,
        'temperature': 37.0,
        'regular_grid_integrity': state['regular_grid']['integrity'],
        'fibonacci_grid_integrity': state['fibonacci_grid']['integrity'],
        'regular_grid_coherence': 1.0,
        'fibonacci_grid_coherence': 1.0,
        'defenses': {k: v for k, v in state['defenses'].items()},
        'targeting_intensity': state['targeting_intensity'],
        'sanctuary_radius': 0,
        'sanctuary_volume': 0,
        'coherence_ratio': 1.0
    })

    # Main simulation loop
    t = 0
    while t < max_time:
        t += dt

        # Update 3D grids and calculate spatial effects
        update_grids_3d(state, dt, t)

        # Calculate legacy coherence values for compatibility
        temperature = state['temperature']
        state['regular_grid']['coherence'] = calculate_quantum_coherence(max(0.01, t), 'regular', temperature)
        state['fibonacci_grid']['coherence'] = calculate_quantum_coherence(max(0.01, t), 'fibonacci', temperature)

        # Check for information sanctuary formation
        calculate_information_sanctuary(state, t, temperature)

        # Update targeting intensity
        update_targeting_intensity(state, t, dt)

        # Print progress
        if int(t * 10) % 100 == 0:
            print(f"Time {t:.1f}: Regular={state['regular_grid']['integrity']:.6f}, "
                  f"Fibonacci={state['fibonacci_grid']['integrity']:.6f}, "
                  f"Sanctuary={state['sanctuary_volume']:.3f}")

        # Store results
        output_interval = params.get('output_interval', 0.1)
        if t % output_interval < dt:
            coherence_ratio = 0
            if state['regular_grid']['coherence'] > 0:
                coherence_ratio = state['fibonacci_grid']['coherence'] / state['regular_grid']['coherence']

            results.append({
                'time': t,
                'temperature': temperature,
                'regular_grid_integrity': state['regular_grid']['integrity'],
                'fibonacci_grid_integrity': state['fibonacci_grid']['integrity'],
                'regular_grid_coherence': state['regular_grid']['coherence'],
                'fibonacci_grid_coherence': state['fibonacci_grid']['coherence'],
                'defenses': {k: v for k, v in state['defenses'].items()},
                'targeting_intensity': state['targeting_intensity'],
                'sanctuary_radius': state['fibonacci_grid'].get('sanctuary_radius', 0),
                'sanctuary_volume': state.get('sanctuary_volume', 0),
                'coherence_ratio': coherence_ratio
            })

    print(f"Simulation complete!")
    print(f"Final regular grid integrity: {state['regular_grid']['integrity']}")
    print(f"Final fibonacci grid integrity: {state['fibonacci_grid']['integrity']}")
    print(f"Final sanctuary volume: {state['sanctuary_volume']:.3f}")

    return results, state


def plot_simulation_results(results, state=None):
    """
    Plot key results from the 3D simulation.
    """
    figures = {}

    # Extract time series data
    times = np.array([r['time'] for r in results])
    temps = np.array([r['temperature'] for r in results])
    reg_integrity = np.array([r['regular_grid_integrity'] for r in results])
    fib_integrity = np.array([r['fibonacci_grid_integrity'] for r in results])
    coherence_ratio = np.array([r['coherence_ratio'] for r in results])
    sanctuary_volume = np.array([r.get('sanctuary_volume', 0) for r in results])

    print(f"Plotting {len(times)} data points...")
    print(f"Integrity ranges - Regular: [{min(reg_integrity):.6f}, {max(reg_integrity):.6f}]")
    print(f"Integrity ranges - Fibonacci: [{min(fib_integrity):.6f}, {max(fib_integrity):.6f}]")

    # Grid integrity over time
    fig1, ax1 = plt.subplots(figsize=(12, 6))
    ax1.plot(times, reg_integrity, 'b-', label='Regular Grid', linewidth=2)
    ax1.plot(times, fib_integrity, 'g-', label='Fibonacci Grid', linewidth=2)
    ax1.set_xlabel('Time')
    ax1.set_ylabel('Grid Integrity')
    ax1.set_title('3D Grid Integrity Over Time')
    ax1.set_ylim([0, 1.1])
    ax1.legend()
    ax1.grid(True, alpha=0.3)
    figures['integrity_time'] = fig1

    # Temperature and sanctuary volume
    fig2, (ax2a, ax2b) = plt.subplots(2, 1, figsize=(12, 8), sharex=True)

    # Temperature profile
    ax2a.plot(times, temps, 'r-', linewidth=2)
    ax2a.set_ylabel('Temperature (°C)')
    ax2a.set_title('Temperature Profile and Sanctuary Formation')
    ax2a.grid(True, alpha=0.3)

    # Sanctuary volume
    ax2b.plot(times, sanctuary_volume, 'purple', linewidth=2)
    ax2b.set_xlabel('Time')
    ax2b.set_ylabel('Sanctuary Volume Fraction')
    ax2b.set_title('Quantum Sanctuary Formation')
    ax2b.grid(True, alpha=0.3)

    figures['temperature_sanctuary'] = fig2

    # Coherence ratio with log scale
    fig3, ax3 = plt.subplots(figsize=(12, 6))
    # Filter out zero or negative values for log plot
    valid_ratios = coherence_ratio[coherence_ratio > 0]
    valid_times = times[coherence_ratio > 0]
    if len(valid_ratios) > 0:
        ax3.semilogy(valid_times, valid_ratios, 'k-', linewidth=2)
    ax3.set_xlabel('Time')
    ax3.set_ylabel('Coherence Ratio (Fibonacci/Regular)')
    ax3.set_title('3D Quantum Coherence Advantage')
    ax3.grid(True, alpha=0.3)
    figures['coherence_ratio'] = fig3

    # Integrity difference with sanctuary highlighting
    fig4, ax4 = plt.subplots(figsize=(12, 6))
    integrity_diff = fib_integrity - reg_integrity
    ax4.plot(times, integrity_diff, color='purple', linewidth=2)

    # Highlight sanctuary periods
    sanctuary_periods = sanctuary_volume > 0.01
    if np.any(sanctuary_periods):
        ax4.fill_between(times, 0, integrity_diff,
                         where=sanctuary_periods, alpha=0.3,
                         color='gold', label='Sanctuary Active')
        ax4.legend()

    ax4.set_xlabel('Time')
    ax4.set_ylabel('Integrity Advantage (Fibonacci - Regular)')
    ax4.set_title('3D Grid Integrity Advantage with Sanctuary Periods')
    ax4.grid(True, alpha=0.3)
    figures['integrity_advantage'] = fig4

    # Add 3D spatial visualizations if state is provided
    if state is not None:
        try:
            spatial_figs = create_3d_visualizations(state)
            figures.update(spatial_figs)
        except Exception as e:
            print(f"Warning: Could not create 3D visualizations: {e}")

    return figures


# Example usage
if __name__ == "__main__":
    # Set simulation parameters for 3D analysis
    params = {
        'max_time': 72,  # Shorter for 3D demo
        'output_interval': 0.5,  # Less frequent output for 3D
        'initial_targeting': 0.05,
        'outer_radius': 1.0,
        'hiv_phase': 'acute',
        'grid_resolution': 15,  # Manageable 3D grid size
        'cell_radius': 10.0,
        'microtubule_count': 18
    }

    print("Starting 3D HIV simulation...")

    # Run simulation
    results, final_state = run_full_hiv_simulation(params)

    # Plot results including 3D visualizations
    figures = plot_simulation_results(results, final_state)
    import pandas as pd

    # Convert results list to DataFrame
    df = pd.DataFrame(results)

    # Save to CSV
    df.to_csv("simulation_results.csv", index=False)

    print("Results saved to simulation_results.csv")
    # Show figures
    plt.show()

    print("3D simulation complete!")
```
