## Supplementary material for "Quantum Coherence Preservation in Fibonacci-Structured Microtubules During HIV-Induced Neuroinflammation": 5 model results: monte_carlo_analysis_report 2.pdf

### Monte Carlo Analysis of HIV Quantum Coherence Simulation

Analysis performed: 2025-05-14 22:05:27

#### Summary

This report presents the results of Monte Carlo analysis and sensitivity analysis performed on the quantum coherence simulation data for different HIV states. The analysis provides insights into the robustness of the simulation results and the sensitivity of the model to various parameters.

#### Monte Carlo Results

Monte Carlo simulations were run with 1000 iterations for each HIV state, varying key model parameters according to their estimated distributions. The table below shows the mean values and 95% confidence intervals for key metrics: ## Monte Carlo Results

Monte Carlo simulations were run with 1000 iterations for each HIV state, varying key model parameters according to their estimated distributions. The table below shows the mean values and 95% confidence intervals for key metrics:

| State | Metric | Mean | Median | Std | 9[illegible] |
|---|---|---|---|---|---|
| acute_hiv | max_ratio | 2.666822e+03 | 2.665296e+03 | 118.652507 | (2427.0065, 2899.[illegible] |
| acute_hiv | half_life_ratio | 3.259996e-01 | 3.279969e-01 | 0.046735 | (0.2258, 0.[illegible] |
| acute_hiv | sanctuary_ratio | 1.177778e-01 | 1.000689e-01 | 0.075680 | (0.0250, 0.[illegible] |
| acute_hiv | final_ratio | 5.292700e-01 | 5.287608e-01 | 0.019377 | (0.4923, 0.[illegible] |
| acute_hiv | auc_ratio | 6.505101e-06 | 4.864360e-07 | 0.000041 | (0.0000, 0.[illegible] |
| art-controlled_hiv | max_ratio | 2.595759e+03 | 2.661351e+03 | 333.268311 | (1454.4810, 2886.[illegible] |
| art-controlled_hiv | half_life_ratio | 2.172638e-01 | 2.075472e-01 | 0.029533 | (0.1833, 0.[illegible] |
| art-controlled_hiv | sanctuary_ratio | 4.418110e-01 | 3.821905e-01 | 0.251483 | (0.1272, 1.[illegible] |
| art-controlled_hiv | final_ratio | 5.296193e-01 | 5.290960e-01 | 0.018945 | (0.4940, 0.[illegible] |
| art-controlled_hiv | auc_ratio | 2.197731e-03 | 2.126198e-04 | 0.011494 | (0.0000, 0.[illegible] |
| chronic_untreated_hiv | max_ratio | 2.661203e+03 | 2.667527e+03 | 116.782708 | (2430.3175, 2892.[illegible] |
| chronic_untreated_hiv | half_life_ratio | 3.942680e-01 | 3.943662e-01 | 0.126246 | (0.2698, 0.[illegible] |
| chronic_untreated_hiv | sanctuary_ratio | 5.481490e-02 | 4.103754e-02 | 0.048218 | (0.0077, 0.[illegible] |
| chronic_untreated_hiv | final_ratio | 5.286736e-01 | 5.291373e-01 | 0.020022 | (0.4895, 0.[illegible] |
| chronic_untreated_hiv | auc_ratio | 7.300902e-07 | 9.034290e-09 | 0.000005 | (0.0000, 0.[illegible] |
| study_volunteer | max_ratio | 2.324078e+02 | 1.972181e+02 | 139.927140 | (71.7156, 611.[illegible] |
| study_volunteer | half_life_ratio | 4.197598e-01 | 3.928571e-01 | 0.117909 | (0.2821, 0.[illegible] |
| study_volunteer | sanctuary_ratio | 2.547384e+00 | 2.484645e+00 | 0.670179 | (1.4518, 4.[illegible] |
| study_volunteer | final_ratio | 5.293341e-01 | 5.291931e-01 | 0.019808 | (0.4922, 0.[illegible] |
| study_volunteer | auc_ratio | 6.102047e-01 | 5.278066e-01 | 0.398408 | (0.0780, 1.[illegible] |

#### Sensitivity Analysis

Sensitivity analysis was performed to identify which model parameters have the greatest influence on the coherence metrics for each HIV state.

The parameters with the strongest correlations for each HIV state are:

Acute Hiv: - For max_ratio: fib_exponent ($r = 1.000$) - For half_life_ratio: state_factor ($r = 0.990$)

Art Controlled Hiv: - For max_ratio: fib_exponent ($r = 1.000$) - For half_life_ratio: reg_exponent ($r = 0.987$)

Chronic Untreated Hiv: - For max_ratio: fib_exponent ($r = 1.000$) - For half_life_ratio: state_factor ($r = 0.994$)

Study Volunteer: - For max_ratio: fib_exponent ($r = 1.000$) - For half_life_ratio: reg_exponent ($r = 0.986$)

For detailed sensitivity results, see the sensitivity plots in the output directory.

#### Statistical Tests

These results indicate: - Max Ratio shows significant differences across HIV states ($p = 0.0000$) - Half Life Ratio shows significant differences across HIV states ($p = 0.0000$) - Sanctuary Ratio shows significant differences across HIV states ($p = 0.0000$) - Auc Ratio shows significant differences across HIV states ($p = 0.0000$)

#### Conclusion

The Monte Carlo analysis confirms the robustness of the observed differences in coherence metrics across HIV states. The model is most sensitive to the state factors, power law exponents, and maximum ratio parameters. This provides support for the hypothesis that Fibonacci-structured systems offer varying degrees of coherence advantage depending on the inflammatory state.
