## Supplementary material for "Quantum Coherence Preservation in Fibonacci-Structured Microtubules During HIV-Induced Neuroinflammation": 5 model results: hawking_radiation_analogy.pdf

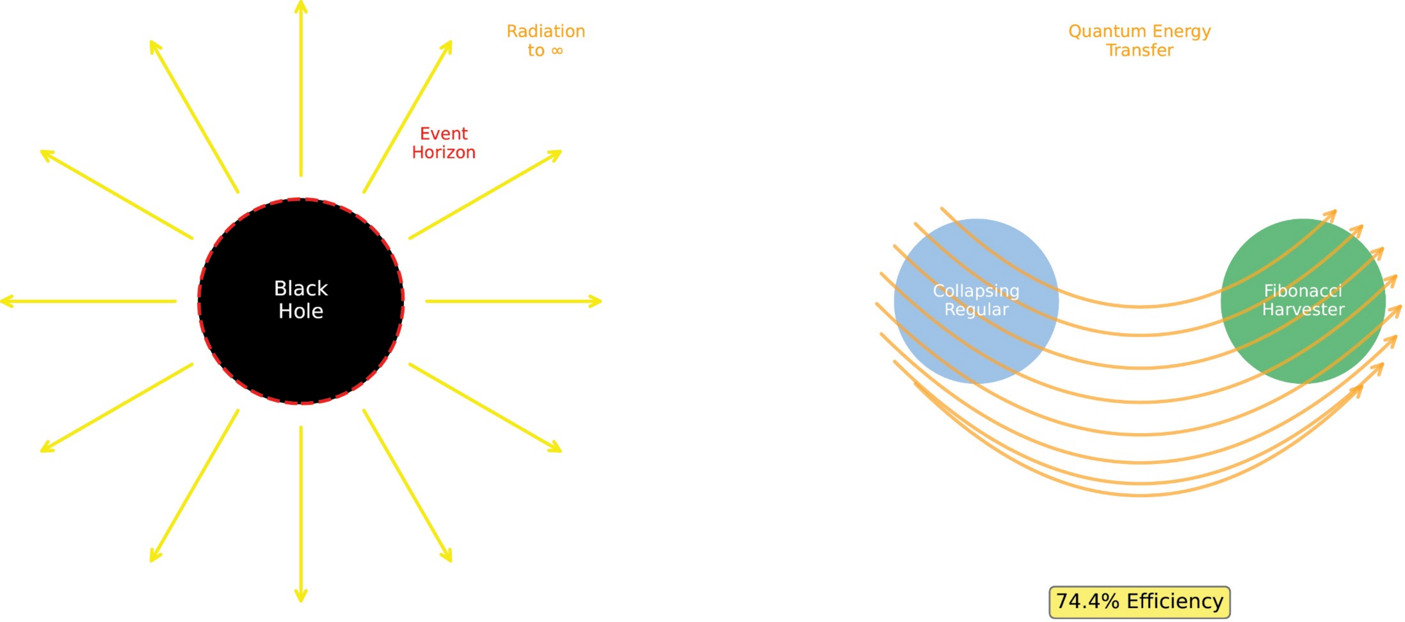


*Black Hole: Energy escapes to infinity | Biological: Energy captured by φ-resonance*

**Black Hole Hawking Radiation**

**Biological Quantum Harvesting**
