## Supplementary material for "Quantum Coherence Preservation in Fibonacci-Structured Microtubules During HIV-Induced Neuroinflammation": 5 model results: monte_carlo_convergence.pdf

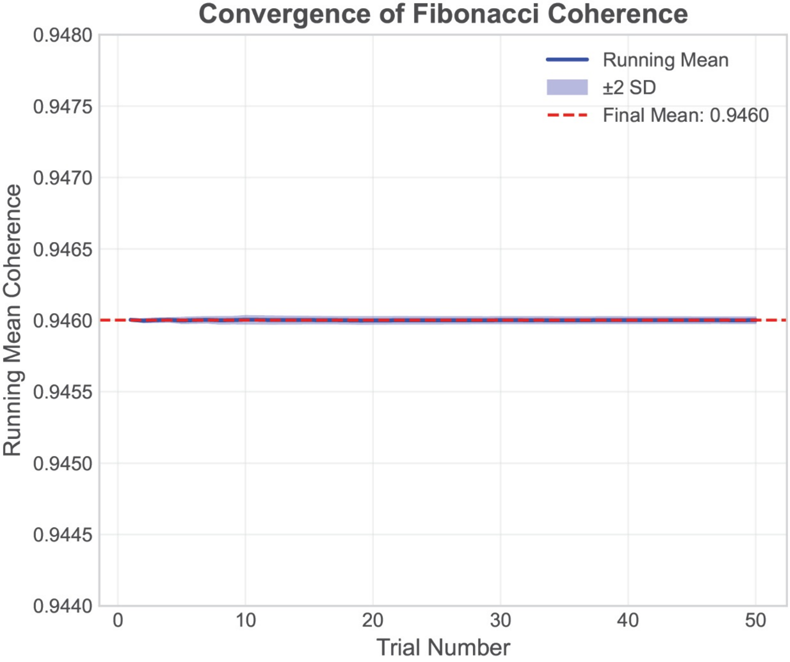


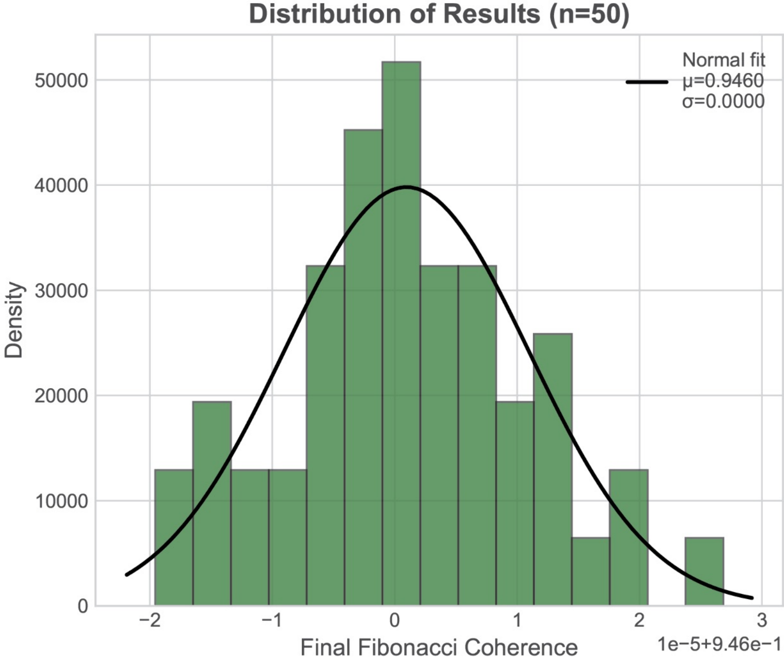


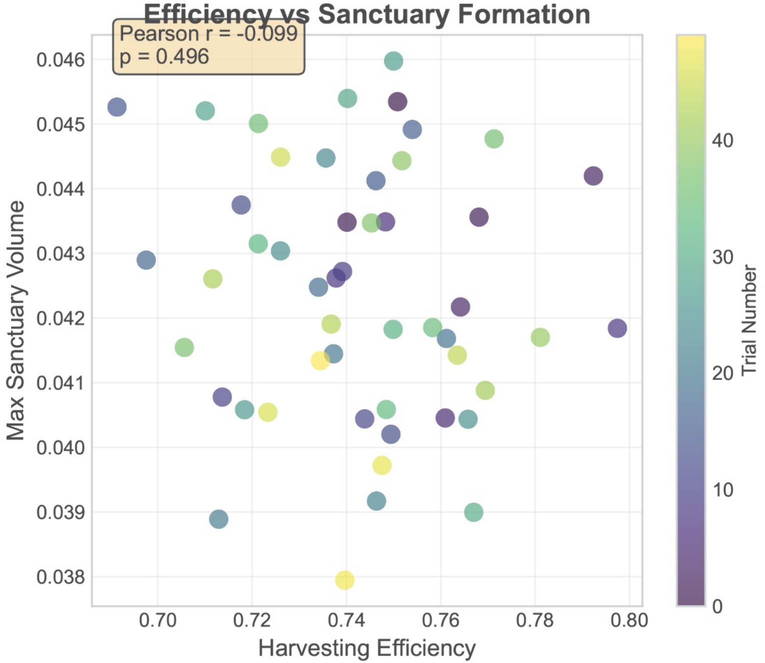


### Monte Carlo Summary Statistics

| Metric | Mean ± SD | 95% CI | Range |
|---|---|---|---|
| Coherence Ratio | 9460 ± 0 | (9460, 9460) | (9460, 9460) |
| Efficiency (%) | 74.2 ± 2.3 | (69.9, 79.0) | (69.1, 79.7) |
| Sanctuary Vol (%) | 4.2 ± 0.2 | (3.9, 4.5) | (3.8, 4.6) |
| Formation Rate | 100% | (50/50 trials) | All trials |
