## Supplementary material for "Quantum Coherence Preservation in Fibonacci-Structured Microtubules During HIV-Induced Neuroinflammation": 5 model results: phi_battle.py

```
import numpy as np
import matplotlib.pyplot as plt
from matplotlib.gridspec import GridSpec
import os
import pandas as pd
import json
from datetime import datetime


# First, let's copy your visualization function
def visualize_phi_targeting_battle(simulation_results, timeline, save_dir="phi_battle_results"):
    """
    Create visualization of how HIV proteins target the golden ratio
    structure of microtubules over time and save the results.

    Args:
        simulation_results (dict): Dictionary containing simulation results
        timeline (np.array): Array of time points
        save_dir (str): Directory to save results

    Returns:
        fig: Matplotlib figure object
    """
    # Create output directory if it doesn't exist
    os.makedirs(save_dir, exist_ok=True)
    timestamp = datetime.now().strftime("%Y%m%d_%H%M%S")

    try:
        # Save raw data first
        # Convert numpy arrays to lists for JSON serialization
        json_data = {
            "timeline": timeline.tolist() if isinstance(timeline, np.ndarray) else timeline,
            "regular_grid_integrity": simulation_results['regular_grid_integrity'].tolist()
            if isinstance(simulation_results['regular_grid_integrity'], np.ndarray)
            else simulation_results['regular_grid_integrity'],
            "fibonacci_grid_integrity": simulation_results['fibonacci_grid_integrity'].tolist()
            if isinstance(simulation_results['fibonacci_grid_integrity'], np.ndarray)
            else simulation_results['fibonacci_grid_integrity'],
            "regular_resonance": simulation_results['regular_resonance'].tolist()
            if isinstance(simulation_results['regular_resonance'], np.ndarray)
            else simulation_results['regular_resonance'],
            "fibonacci_resonance": simulation_results['fibonacci_resonance'].tolist()
            if isinstance(simulation_results['fibonacci_resonance'], np.ndarray)
            else simulation_results['fibonacci_resonance'],
            "targeting_intensity": simulation_results['targeting_intensity'].tolist()
            if isinstance(simulation_results['targeting_intensity'], np.ndarray)
            else simulation_results['targeting_intensity'],
            "metadata": {
                "timestamp": timestamp,
                "description": "HIV phi-targeting battle simulation results"
            }
        }

        # Save defense activation data separately due to its structure
        if 'defense_activation' in simulation_results:
            defense_data = {}
            defense_mechanisms = ['Phi Stability', 'Resonance',
                                  'Acetylation', 'Binding Sites', 'Adaptation']

            for i, mechanism in enumerate(defense_mechanisms):
                defense_data[mechanism] = simulation_results['defense_activation'][i].tolist() \
                    if isinstance(simulation_results['defense_activation'][i], np.ndarray) \
                    else simulation_results['defense_activation'][i]

            json_data["defense_activation"] = defense_data

        # Save structure heatmap data
        if 'structure_heatmap' in simulation_results:
            if isinstance(simulation_results['structure_heatmap'], np.ndarray):
                # Save as separate NPY file for large arrays
                heatmap_path = os.path.join(save_dir, f"structure_heatmap_{timestamp}.npy")
                np.save(heatmap_path, simulation_results['structure_heatmap'])
                json_data["structure_heatmap_file"] = heatmap_path
            else:
                # If it's small enough, include in JSON
                json_data["structure_heatmap"] = simulation_results['structure_heatmap']

        # Save JSON data
        json_path = os.path.join(save_dir, f"phi_battle_data_{timestamp}.json")
        with open(json_path, 'w') as f:
            json.dump(json_data, f, indent=2)

        # Also save as CSV for easier analysis
        csv_data = {
            "Time": timeline,
            "Regular_Grid_Integrity": simulation_results['regular_grid_integrity'],
            "Fibonacci_Grid_Integrity": simulation_results['fibonacci_grid_integrity'],
            "Regular_Resonance": simulation_results['regular_resonance'],
            "Fibonacci_Resonance": simulation_results['fibonacci_resonance'],
            "Targeting_Intensity": simulation_results['targeting_intensity']
        }

        # Add defense mechanisms if available
        if 'defense_activation' in simulation_results:
            defense_mechanisms = ['Phi Stability', 'Resonance',
                                  'Acetylation', 'Binding Sites', 'Adaptation']
            for i, mechanism in enumerate(defense_mechanisms):
                csv_data[f"Defense_{mechanism.replace(' ', '_')}"] = simulation_results['defense_activation'][i]

        # Save CSV
        df = pd.DataFrame(csv_data)
        csv_path = os.path.join(save_dir, f"phi_battle_data_{timestamp}.csv")
        df.to_csv(csv_path, index=False)

        print(f"Saved simulation data to:\n- {json_path}\n- {csv_path}")

        # Now create the visualization
        fig = plt.figure(figsize=(15, 12))
        gs = GridSpec(3, 2, figure=fig)

        # 1. Microtubule structural integrity over time
        ax1 = fig.add_subplot(gs[0, 0])
        ax1.plot(timeline, simulation_results['regular_grid_integrity'],
                 'r-', label='Regular Grid')
        ax1.plot(timeline, simulation_results['fibonacci_grid_integrity'],
                 'b-', label='Fibonacci Grid')
        ax1.set_title('Microtubule Structural Integrity During HIV Attack')
        ax1.set_ylabel('Integrity (%)')
        ax1.legend()

        # 2. Phi resonance preservation
        ax2 = fig.add_subplot(gs[0, 1])
        ax2.plot(timeline, simulation_results['regular_resonance'],
                 'r-', label='Regular Grid')
        ax2.plot(timeline, simulation_results['fibonacci_resonance'],
                 'b-', label='Fibonacci Grid')
        ax2.set_title('Golden Ratio Resonance Preservation')
        ax2.set_ylabel('Resonance Strength')
        ax2.legend()

        # 3. HIV targeting intensity (varies with disease progression)
        ax3 = fig.add_subplot(gs[1, 0])
        ax3.plot(timeline, simulation_results['targeting_intensity'], 'g-')
        ax3.axhline(y=0.5, color='orange', linestyle='--',
                    label='Critical Threshold')
        ax3.set_title('HIV Phi-Targeting Intensity')
        ax3.set_ylabel('Attack Intensity')
        ax3.legend()

        # 4. Defense mechanism activation
        ax4 = fig.add_subplot(gs[1, 1])
        defense_mechanisms = ['Phi Stability', 'Resonance',
                              'Acetylation', 'Binding Sites', 'Adaptation']

        if 'defense_activation' in simulation_results:
            for i, mechanism in enumerate(defense_mechanisms):
                if i < len(simulation_results['defense_activation']):
                    ax4.plot(timeline, simulation_results['defense_activation'][i],
                             label=mechanism)
            ax4.set_title('Defense Mechanism Activation')
            ax4.set_ylabel('Activation Level')
            ax4.legend(loc='upper center', bbox_to_anchor=(0.5, -0.15), ncol=3)
        else:
            ax4.text(0.5, 0.5, "Defense activation data not available",
                     ha='center', va='center', transform=ax4.transAxes)

        # 5. Microtubule structural heatmap (2D visualization)
        ax5 = fig.add_subplot(gs[2, :])

        if 'structure_heatmap' in simulation_results:
            # Create a heatmap showing phi-preservation in microtubule structure
            im = ax5.imshow(simulation_results['structure_heatmap'],
                            aspect='auto', cmap='viridis')
            ax5.set_title('Microtubule Phi-Structure Preservation Over Time')
            ax5.set_xlabel('Time')
            ax5.set_ylabel('Microtubule Section')
            plt.colorbar(im, ax=ax5, label='Phi-Preservation')
        else:
            ax5.text(0.5, 0.5, "Heatmap data not available",
                     ha='center', va='center', transform=ax5.transAxes)

        # Add timestamp and metadata to figure
        fig.text(0.98, 0.01, f"Generated: {timestamp}",
                 ha='right', va='bottom', fontsize=8)

        plt.tight_layout()

        # Save visualization
        fig_path = os.path.join(save_dir, f"phi_battle_visualization_{timestamp}.png")
        plt.savefig(fig_path, dpi=300, bbox_inches='tight')

        # Also save in vector format for publication
        fig_path_pdf = os.path.join(save_dir, f"phi_battle_visualization_{timestamp}.pdf")
        plt.savefig(fig_path_pdf, bbox_inches='tight')

        print(f"Saved visualization to:\n- {fig_path}\n- {fig_path_pdf}")

        return fig

    except Exception as e:
        print(f"Error in visualization or data saving: {e}")
        import traceback
        print(traceback.format_exc())

        # Try to save error information
        error_file = os.path.join(save_dir, f"error_log_{timestamp}.txt")
        with open(error_file, 'w') as f:
            f.write(f"Error occurred during visualization: {e}\n\n")
            f.write(traceback.format_exc())
            f.write("\n\nPartial simulation data:\n")
            f.write(str(simulation_results))

        return None


# Now let's add the simulation function to generate data for our visualization
def simulate_phi_targeting_battle(duration=100, time_steps=1000, hiv_progression_type="acute"):
    """
    Simulate the battle between HIV proteins and microtubule phi structures.

    Args:
        duration: Duration of simulation in arbitrary time units
        time_steps: Number of time steps for simulation
        hiv_progression_type: Type of HIV progression ('acute', 'chronic', 'controlled')

    Returns:
        simulation_results: Dictionary containing simulation results
        timeline: Array of time points
    """
    # Create timeline
    timeline = np.linspace(0, duration, time_steps)

    # Parameters based on HIV progression type
    if hiv_progression_type == "acute":
        # Rapid increase in targeting intensity, severe damage
        attack_rate = 0.05
        max_intensity = 0.9
        recovery_ability = 0.01
        defense_activation_rate = 0.02
    elif hiv_progression_type == "chronic":
        # Gradual increase in targeting, sustained damage
        attack_rate = 0.02
        max_intensity = 0.7
        recovery_ability = 0.005
        defense_activation_rate = 0.01
    elif hiv_progression_type == "controlled":
        # Limited targeting with ART, partial recovery
        attack_rate = 0.03
        max_intensity = 0.5
        recovery_ability = 0.02
        defense_activation_rate = 0.03
    else:
        # Default parameters
        attack_rate = 0.03
        max_intensity = 0.7
        recovery_ability = 0.01
        defense_activation_rate = 0.02

    # Initialize arrays
    targeting_intensity = np.zeros(time_steps)
    regular_grid_integrity = np.ones(time_steps)
    fibonacci_grid_integrity = np.ones(time_steps)
    regular_resonance = np.ones(time_steps)
    fibonacci_resonance = np.ones(time_steps)

    # Defense mechanisms (5 different types)
    defense_activation = np.zeros((5, time_steps))

    # Microtubule structure heatmap (spatial x temporal)
    structure_sections = 50
    structure_heatmap = np.ones((structure_sections, time_steps))

    # Simulation parameters
    phi_advantage = 1.618  # Golden ratio advantage
    threshold = 0.5
    resonance_decay_factor = 0.1
    fibonacci_resilience = 3.0

    # Run simulation
    for t in range(1, time_steps):
        # Update targeting intensity based on progression model
        if hiv_progression_type == "acute":
            # Quick ramp up with early peak
            if timeline[t] < duration * 0.2:
                targeting_intensity[t] = min(max_intensity,
                                             targeting_intensity[t - 1] + attack_rate * (
                                                         1 - targeting_intensity[t - 1]))
            else:
                # Slight decline due to immune response
                targeting_intensity[t] = max(0.4,
                                             targeting_intensity[t - 1] - 0.002 * (targeting_intensity[t - 1] - 0.4))
        elif hiv_progression_type == "chronic":
            # Gradual increase to plateau
            targeting_intensity[t] = min(max_intensity,
                                         targeting_intensity[t - 1] + attack_rate * (
                                                     1 - targeting_intensity[t - 1] / max_intensity))
        elif hiv_progression_type == "controlled":
            # Increase then decrease due to ART
            if timeline[t] < duration * 0.3:
                targeting_intensity[t] = min(max_intensity,
                                             targeting_intensity[t - 1] + attack_rate * (
                                                         1 - targeting_intensity[t - 1]))
            else:
                # Decline due to ART
                targeting_intensity[t] = max(0.2,
                                             targeting_intensity[t - 1] - 0.01 * (targeting_intensity[t - 1] - 0.2))

        # Update defense mechanisms activation
        for d in range(5):
            # Different defense mechanisms activate at different rates and times
            activation_threshold = 0.1 + d * 0.05
            activation_delay = int(time_steps * (0.05 + d * 0.02))

            if t > activation_delay and targeting_intensity[t] > activation_threshold:
                defense_rate = defense_activation_rate * (1 + d * 0.1)
                defense_activation[d, t] = min(1.0, defense_activation[d, t - 1] +
                                               defense_rate * (targeting_intensity[t] - activation_threshold))
            else:
                defense_activation[d, t] = max(0, defense_activation[d, t - 1] - 0.01)

        # Calculate damage to regular grid
        damage_regular = targeting_intensity[t] * (1 - defense_activation[0, t] * 0.5)
        recovery_regular = recovery_ability * defense_activation[3, t]
        regular_grid_integrity[t] = max(0, regular_grid_integrity[t - 1] -
                                        damage_regular + recovery_regular)

        # Calculate damage to fibonacci grid (more resistant)
        damage_fibonacci = targeting_intensity[t] * (1 - defense_activation[0, t] * 0.8) / fibonacci_resilience
        recovery_fibonacci = recovery_ability * defense_activation[3, t] * phi_advantage
        fibonacci_grid_integrity[t] = max(0, fibonacci_grid_integrity[t - 1] -
                                          damage_fibonacci + recovery_fibonacci)

        # Update resonance
        regular_resonance[t] = regular_grid_integrity[t] * (1 - resonance_decay_factor * targeting_intensity[t])
        fibonacci_resonance[t] = fibonacci_grid_integrity[t] * (1 - resonance_decay_factor *
                                                                targeting_intensity[t] / phi_advantage)

        # Update structure heatmap (different sections affected differently)
        for s in range(structure_sections):
            # Fibonacci sections (every fibonacci number section)
            is_fibonacci_section = s in [1, 2, 3, 5, 8, 13, 21, 34] or s % 5 == 0

            section_resistance = 1.0 + (phi_advantage - 1.0) * is_fibonacci_section
            section_damage = targeting_intensity[t] * (
                        1 - defense_activation[2, t] * (0.3 + 0.5 * is_fibonacci_section))

            # More targeted damage to fibonacci sections when targeting exceeds threshold
            if targeting_intensity[t] > threshold and is_fibonacci_section:
                section_damage *= 1.2

            # Random fluctuations
            noise = np.random.normal(0, 0.02)

            # Update section integrity
            current_integrity = structure_heatmap[s, t - 1]
            damage = section_damage / section_resistance
            repair = recovery_ability * defense_activation[4, t] * (1 + 0.5 * is_fibonacci_section)

            structure_heatmap[s, t] = max(0, min(1, current_integrity - damage + repair + noise))

    # Compile results
    simulation_results = {
        'regular_grid_integrity': regular_grid_integrity,
        'fibonacci_grid_integrity': fibonacci_grid_integrity,
        'regular_resonance': regular_resonance,
        'fibonacci_resonance': fibonacci_resonance,
        'targeting_intensity': targeting_intensity,
        'defense_activation': defense_activation,
        'structure_heatmap': structure_heatmap
    }

    return simulation_results, timeline


# Main function to run simulation and visualization
def run_phi_battle_simulation(progression_type="acute", duration=100, time_steps=1000):
    """
    Run the entire phi targeting battle simulation and visualization.

    Args:
        progression_type: Type of HIV progression ('acute', 'chronic', 'controlled')
        duration: Duration of simulation
        time_steps: Number of time steps

    Returns:
        None
    """
    print(f"Running phi battle simulation for {progression_type} HIV progression...")

    # Run simulation
    simulation_results, timeline = simulate_phi_targeting_battle(
        duration=duration,
        time_steps=time_steps,
        hiv_progression_type=progression_type
    )

    # Create visualization
    output_dir = f"phi_battle_results_{progression_type}"
    fig = visualize_phi_targeting_battle(simulation_results, timeline, save_dir=output_dir)

    if fig:
        print(f"Simulation and visualization completed successfully!")
        plt.show()  # Show the figure (if in interactive environment)
    else:
        print("Simulation completed but visualization encountered errors.")


# Execute the simulation for all progression types
if __name__ == "__main__":
    # Run acute HIV simulation
    run_phi_battle_simulation(progression_type="acute")

    # Run chronic HIV simulation
    run_phi_battle_simulation(progression_type="chronic")

    # Run ART-controlled HIV simulation
    run_phi_battle_simulation(progression_type="controlled")
```
