## Supplementary material for "Quantum Coherence Preservation in Fibonacci-Structured Microtubules During HIV-Induced Neuroinflammation": 5 model results: phi_battle_analysis.py

```
def load_phi_battle_data(data_path):
    """
    Load saved phi battle simulation data for analysis

    Args:
        data_path (str): Path to the JSON or CSV data file

    Returns:
        dict: Loaded simulation data
    """
    import os
    import json
    import pandas as pd
    import numpy as np

    try:
        if data_path.endswith('.json'):
            # Load JSON data
            with open(data_path, 'r') as f:
                data = json.load(f)

            # Check if heatmap is stored separately
            if 'structure_heatmap_file' in data and os.path.exists(data['structure_heatmap_file']):
                data['structure_heatmap'] = np.load(data['structure_heatmap_file'])

            return data

        elif data_path.endswith('.csv'):
            # Load CSV data
            df = pd.read_csv(data_path)

            # Convert to dictionary format
            data = {
                'timeline': df['Time'].values,
                'regular_grid_integrity': df['Regular_Grid_Integrity'].values,
                'fibonacci_grid_integrity': df['Fibonacci_Grid_Integrity'].values,
                'regular_resonance': df['Regular_Resonance'].values,
                'fibonacci_resonance': df['Fibonacci_Resonance'].values,
                'targeting_intensity': df['Targeting_Intensity'].values
            }

            # Check for defense mechanism columns
            defense_mechanisms = ['Phi_Stability', 'Resonance',
                                  'Acetylation', 'Binding_Sites', 'Adaptation']

            defense_data = []
            for mechanism in defense_mechanisms:
                col_name = f"Defense_{mechanism}"
                if col_name in df.columns:
                    defense_data.append(df[col_name].values)

            if defense_data:
                data['defense_activation'] = defense_data

            return data

        else:
            print(f"Unsupported file format: {data_path}")
            return None

    except Exception as e:
        print(f"Error loading data: {e}")
        import traceback
        print(traceback.format_exc())
        return None
```
