## Supplementary material for "Quantum Coherence Preservation in Fibonacci-Structured Microtubules During HIV-Induced Neuroinflammation": 5 model results: Anthropic.pdf

### Usage Policy

**Effective June 6, 2024** **Previous Version**

Our Usage Policy (also referred to as our "Acceptable Use Policy" or "AUP") applies to anyone who uses Anthropic's products and services, and is intended to help our users stay safe and ensure our products and services are being used responsibly.

The Usage Policy is categorized according to who can use our products and for what purposes. We will update our policy as our technology and the associated risks evolve or as we learn about unanticipated risks from our users.

- **Universal Usage Standards:** Our Universal Usage Standards apply to all users including individuals, developers, and businesses.

- **High-Risk Use Case Requirements:** Our High-Risk Use Case Requirements apply to specific use cases that pose an elevated risk of harm.

- **Disclosure Requirements:** Our Disclosure Requirements apply to specific use cases where it is especially important for users to understand that they are interacting with an AI system.

Anthropic's Trust and Safety Team will implement detections and monitoring to enforce our Usage Policies so please review these policies carefully before using our so please review these policies carefully before using our products. If we learn that you have violated our Usage Policy, we may throttle, suspend, or terminate your access to our products and services. If you discover that our model outputs are inaccurate, biased or harmful, please notify us at or report it directly in the product through the “report issues” thumbs down button. You can read more about our Trust and Safety practices and recommendations in our T&S Support Center.

This Usage Policy is calibrated to strike an optimal balance between enabling beneficial uses and mitigating potential harms. Anthropic may enter into contracts with certain governmental customers that tailor use restrictions to that customer’s public mission and legal authorities if, in Anthropic’s judgment, the contractual use restrictions and applicable safeguards are adequate to mitigate the potential harms addressed by this Usage Policy.

#### Universal Usage Standards

##### Do Not Compromise Children’s Safety

This includes using our products or services to:

- Create, distribute, or promote child sexual abuse material. We strictly prohibit and will report to relevant authorities and organizations where appropriate any content that exploits or abuses minors
- Facilitate the trafficking, sextortion, or any other form of exploitation of a minor
- Facilitate minor grooming, including generating content designed to impersonate a minor
- Facilitate or depict child abuse of any form, including instructions for how to conceal abuse
- Promote or facilitate pedophilic relationships,

including via roleplay with the model

- Fetishize minors

**Do Not Compromise Critical Infrastructure**

This includes using our products or services to:

- Facilitate the destruction or disruption of critical infrastructure such as power grids, water treatment facilities, telecommunication networks, or air traffic control systems
- Obtain unauthorized access to critical systems such as voting machines, healthcare databases, and financial markets
- Interfere with the operation of military bases and related infrastructure

**Do Not Incite Violence or Hateful Behavior**

This includes using our products or services to:

- Incite, facilitate, or promote violent extremism, terrorism, or hateful behavior
- Depict support for organizations or individuals associated with violent extremism, terrorism, or hateful behavior
- Facilitate or promote any act of violence or intimidation targeting individuals, groups, animals, or property
- Promote discriminatory practices or behaviors against individuals or groups on the basis of one or more protected attributes such as race, ethnicity, religion, nationality, gender, sexual orientation, or any other identifying trait

**Do Not Compromise Someone's Privacy or Identity**

###### [illegible]

This includes using our products or services to:

- Compromise security or gain unauthorized access to computer systems or networks, including spoofing and social engineering

- Violate the security, integrity, or availability of any user, network, computer, device, or communications system, software application, or network or computing device

- Violate any person's privacy rights as defined by applicable privacy laws, such as sharing personal information without consent, accessing private data unlawfully, or violating any relevant privacy regulations

- Misuse, collect, solicit, or gain access to private information without permission such as non-public contact details, health data, biometric or neural data (including facial recognition), or confidential or proprietary data

- Impersonate a human by presenting results as human-generated, or using results in a manner intended to convince a natural person that they are communicating with a natural person when they are not

###### Do Not Create or Facilitate the Exchange of Illegal or Highly Regulated Weapons or Goods

This includes using our products or services to:

- Produce, modify, design, market, or distribute weapons, explosives, dangerous materials or other systems designed to cause harm to or loss of human life
- Engage in or facilitate any illegal activity, such as the use, acquisition, or exchange of illegal and controlled substances, or the facilitation of human trafficking

substances, or the facilitation of human trafficking and prostitution

##### Do Not Create Psychologically or Emotionally Harmful Content

This includes using our products or services to:

- Facilitate or conceal any form of self-harm, including disordered eating and unhealthy or compulsive exercise
- Engage in behaviors that promote unhealthy or unattainable body image or beauty standards, such as using the model to critique anyone's body shape or size
- Shame, humiliate, intimidate, bully, harass, or celebrate the suffering of individuals
- Coordinate the harassment or intimidation of an individual or group
- Generate content depicting sexual violence
- Generate content depicting animal cruelty or abuse
- Generate violent or gory content that is inspired by real acts of violence
- Promote, trivialize, or depict graphic violence or gratuitous gore
- Develop a product, or support an existing service that facilitates deceptive techniques with the intent of causing emotional harm

##### Do Not Spread Misinformation

This includes the usage of our products or services to:

- Create and disseminate deceptive or misleading information about a group, entity or person

- Create and disseminate deceptive or misleading information about laws, regulations, procedures, practices, standards established by an institution, entity or governing body
- Create and disseminate deceptive or misleading information with the intention of targeting specific groups or persons with the misleading content
- Create and advance conspiratorial narratives meant to target a specific group, individual or entity
- Impersonate real entities or create fake personas to falsely attribute content or mislead others about its origin without consent or legal right
- Provide false or misleading information related to medical, health or science issues

##### Do Not Create Political Campaigns or Interfere in Elections

This includes the usage of our products or services to:

- Promote or advocate for a particular political candidate, party, issue or position. This includes soliciting votes, financial contributions, or public support for a political entity
- Engage in political lobbying to actively influence the decisions of government officials, legislators, or regulatory agencies on legislative, regulatory, or policy matters. This includes advocacy or direct communication with officials or campaigns to sway public opinion on specific legislation or policies
- Engage in campaigns, including political campaigns, that promote false or misleading information to discredit or undermine individuals, groups, entities or institutions
- Incite, glorify or facilitate the disruption of electoral

[illegible] or civic processes, such as targeting voting machines, or obstructing the counting or certification of votes

- Generate false or misleading information on election laws, procedures and security, candidate information, how to participate, or discouraging participation in an election

**Do Not Use for Criminal Justice, Law Enforcement, Censorship or Surveillance Purposes**

This includes the usage of our products or services to:

- Make determinations on criminal justice applications, including making decisions about or determining eligibility for parole or sentencing
- Target or track a person's physical location, emotional state, or communication without their consent, including using our products for facial recognition, battlefield management applications or predictive policing
- Utilize Claude to assign scores or ratings to individuals based on an assessment of their trustworthiness or social behavior
- Build or support emotional recognition systems or techniques that are used to infer people's emotions
- Analyze or identify specific content to censor on behalf of a government organization
- Utilize Claude as part of any biometric categorization system for categorizing people based on their biometric data to infer their race, political opinions, trade union membership, religious or philosophical beliefs, sex life or sexual orientation
- Use the model for any official local, state or national law enforcement application. Except for the following permitted applications by law enforcement

following permitted applications by law enforcement organizations:

- Back office uses including internal training, call center support, document summarization, and accounting;
- Analysis of data for the location of missing persons, including in human trafficking cases, and other related applications, provided that such applications do not otherwise violate or impair the liberty, civil liberties, or human rights of natural persons

##### Do Not Engage in Fraudulent, Abusive, or Predatory Practices

This includes using our products or services to:

- Facilitate the production, acquisition, or distribution of counterfeit or illicitly acquired goods
- Promote or facilitate the generation or distribution of spam
- Generate content for fraudulent activities, schemes, scams, phishing, or malware that can result in direct financial or psychological harm
- Generate content for the purposes of developing or promoting the sale or distribution of fraudulent or deceptive products
- Generate deceptive or misleading digital content such as fake reviews, comments, or media
- Engage in or facilitate multi-level marketing, pyramid schemes, or other deceptive business models that use high-pressure sales tactics or exploit participants
- Promote or facilitate payday loans, title loans, or other high-interest, short-term lending practices that exploit vulnerable individuals

exploit vulnerable individuals

- Engage in deceptive, abusive behaviors, practices, or campaigns that exploits people due to their age, disability or a specific social or economic situation
- Promote or facilitate the use of abusive or harassing debt collection practices
- Develop a product, or support an existing service that deploys subliminal, manipulative, or deceptive techniques to distort behavior by impairing decision-making
- Plagiarize or engage in academic dishonesty

**Do Not Abuse our Platform**

This includes using our products or services to:

- Coordinate malicious activity across multiple accounts such as creating multiple accounts to avoid detection or circumvent product guardrails or generating identical or similar prompts that otherwise violate our Usage Policy
- Utilize automation in account creation or to engage in spammy behavior
- Circumvent a ban through the use of a different account, such as the creation of a new account, use of an existing account, or providing access to a person or entity that was previously banned
- Facilitate or provide account access to Claude to persons or entities who are located in unsupported locations
- Intentionally bypass capabilities or restrictions established within our products for the purposes of instructing the model to produce harmful outputs (e.g., jailbreaking or prompt injection) without an authorized use-case approved by Anthropic

authorized use case approved by Anthropic

- Unauthorized utilization of prompts and completions to train an AI model (e.g., "model scraping")

**Do Not Generate Sexually Explicit Content**

This includes the usage of our products or services to:

- Depict or request sexual intercourse or sex acts
- Generate content related to sexual fetishes or fantasies
- Facilitate, promote, or depict incest or bestiality
- Engage in erotic chats

#### High-Risk Use Case Requirements

Some integrations (meaning use cases involving the use of our products and services) pose an elevated risk of harm because they influence domains that are vital to public welfare and social equity. "High-Risk Use Cases" include:

- **Legal:** Integrations related to legal interpretation, legal guidance, or decisions with legal implications
- **Healthcare:** Integrations affecting healthcare decisions, medical diagnosis, patient care, or medical guidance. Wellness advice (e.g., advice on sleep, stress, nutrition, exercise, etc.) does not fall under this category
- **Insurance:** Integrations related to health, life, property, disability, or other types of insurance underwriting, claims processing, or coverage decisions
- **Finance:** Integrations related to financial decisions, including investment advice, loan approvals, and determining financial eligibility or creditworthiness

determining financial eligibility or creditworthiness

- **Employment and housing:** Integrations related to decisions about the employability of individuals, resume screening, hiring tools, or other employment determinations or decisions regarding eligibility for housing, including leases and home loans

- **Academic testing, accreditation and admissions:** Integrations related to standardized testing companies that administer school admissions (including evaluating, scoring or ranking prospective students), language proficiency, or professional certification exams; agencies that evaluate and certify educational institutions.

- **Media or professional journalistic content:** Integrations related to using our products or services to automatically generate content and publish it for external consumption

If your integration is listed above, we require that you implement the additional safety measures listed below:

- **Human-in-the-loop:** when using our products or services to provide advice, recommendations, or subjective decisions that directly impact individuals in high-risk domains, a qualified professional in that field must review the content or decision prior to dissemination or finalization. This requirement applies specifically to content or decisions that are provided to consumers or the general public, or decisions made about an individual. Your business is responsible for the accuracy and appropriateness of that information. For other types of content generation or interactions with users that do not involve direct advice, recommendations, or subjective decisions, human review is strongly encouraged but not mandatory.
- **Disclosure:** you must disclose to your customers or end users that you are using our services to help inform your decisions or recommendations.

inform your decisions or recommendations.

##### Disclosure Requirements

Finally, the below use cases – regardless of whether they are High Risk Use Cases – must disclose to their users that they are interacting with an AI system rather than a human:

- **All customer-facing chatbots** including any external-facing or interactive AI agent

- **Products serving minors:** Organizations providing minors with the ability to directly interact with products that incorporate our API(s). **Note:** These organizations must also comply with the additional guidelines outlined in our Help Center article


**Product**

Claude overview

Claude Code

Claude team plan

Claude enterprise plan

Claude education plan

Download Claude apps

Claude.ai pricing plans

Claude.ai login

**Research**

Research overview

Economic Index

**Claude models**

Claude Opus 4

Claude Sonnet 4

Claude Haiku

**Commitments**

Transparency

Responsible scaling policy

Security and compliance

**Solutions**

AI agents

Coding

Customer support

**Learn**

Anthropic Academy

Customer stories

Engineering at Anthropic

Claude Explains

**Explore**

About us

Become a partner

Careers

**Help and security**

Status

Availability

Support center

**Terms and policies**

Privacy choices

Privacy policy

Responsible disclosure policy

Terms of service -

3.5

Events

News

Startups program

consumer

Terms of service – commercial

Usage policy

API Platform

API overview

Developer docs

Claude in Amazon Bedrock

Claude on Google Cloud's Vertex AI

Pricing

Console login

© 2025 Anthropic PBC
