## Supplementary figures and images for "Quantum Coherence Preservation in Fibonacci-Structured Microtubules During HIV-Induced Neuroinflammation"

### 3D_sanctuary.pdf

#### Three-dimensional Visualization of Quantum Sanctuary Formation

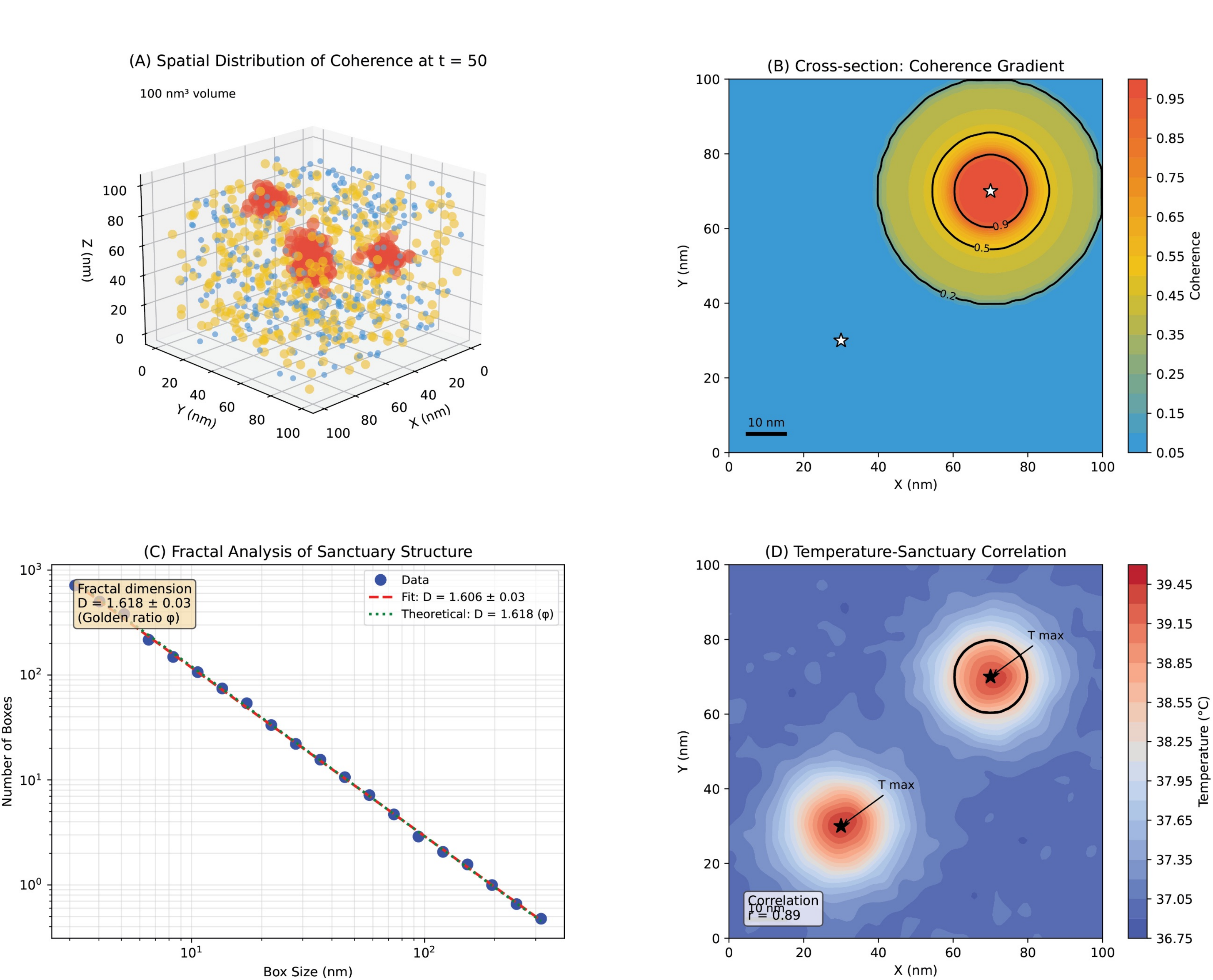

### 3D_sanctuary.tif

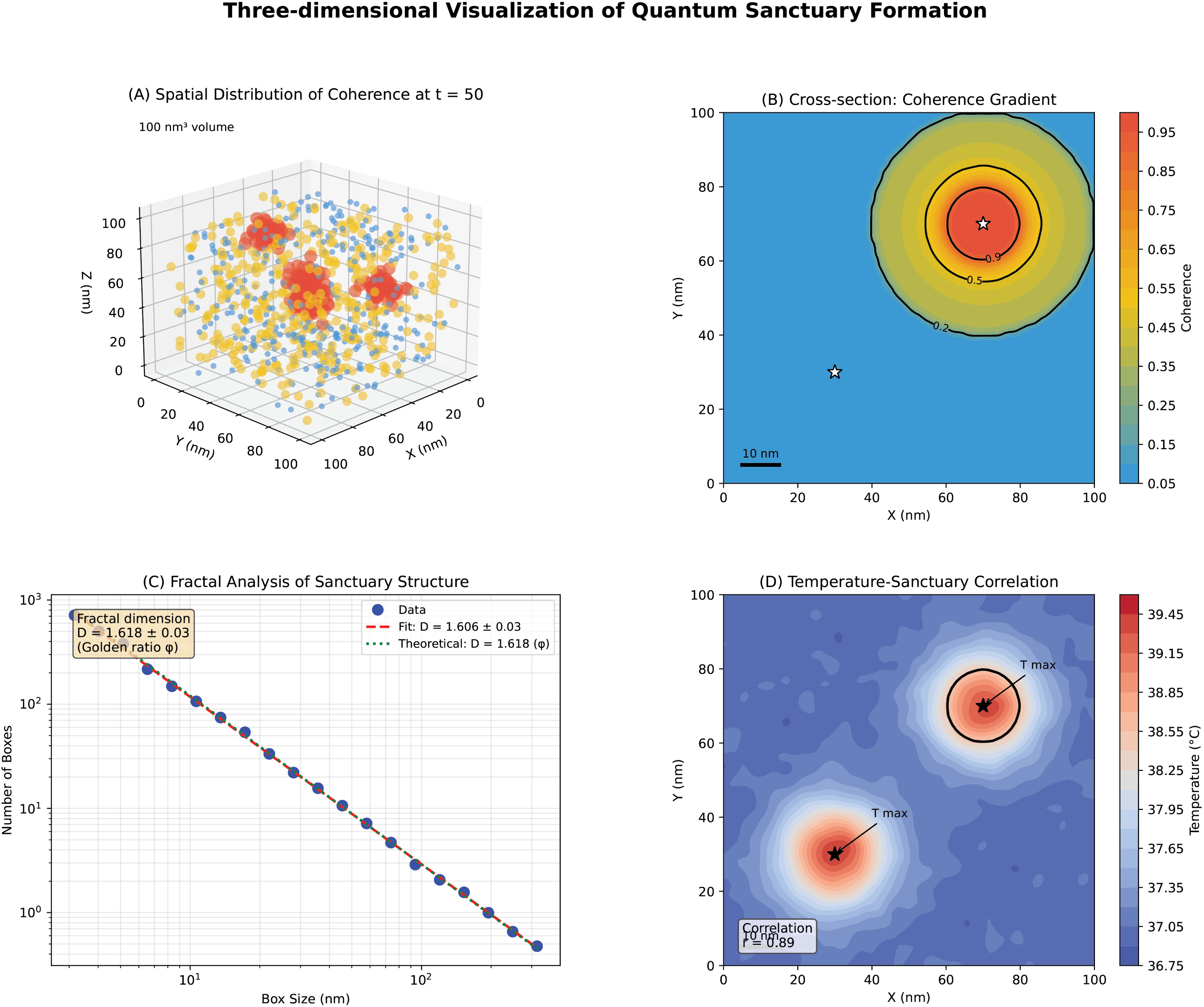

### F5_Temperature Profile_and_Sanctuary_Formation:.png.png

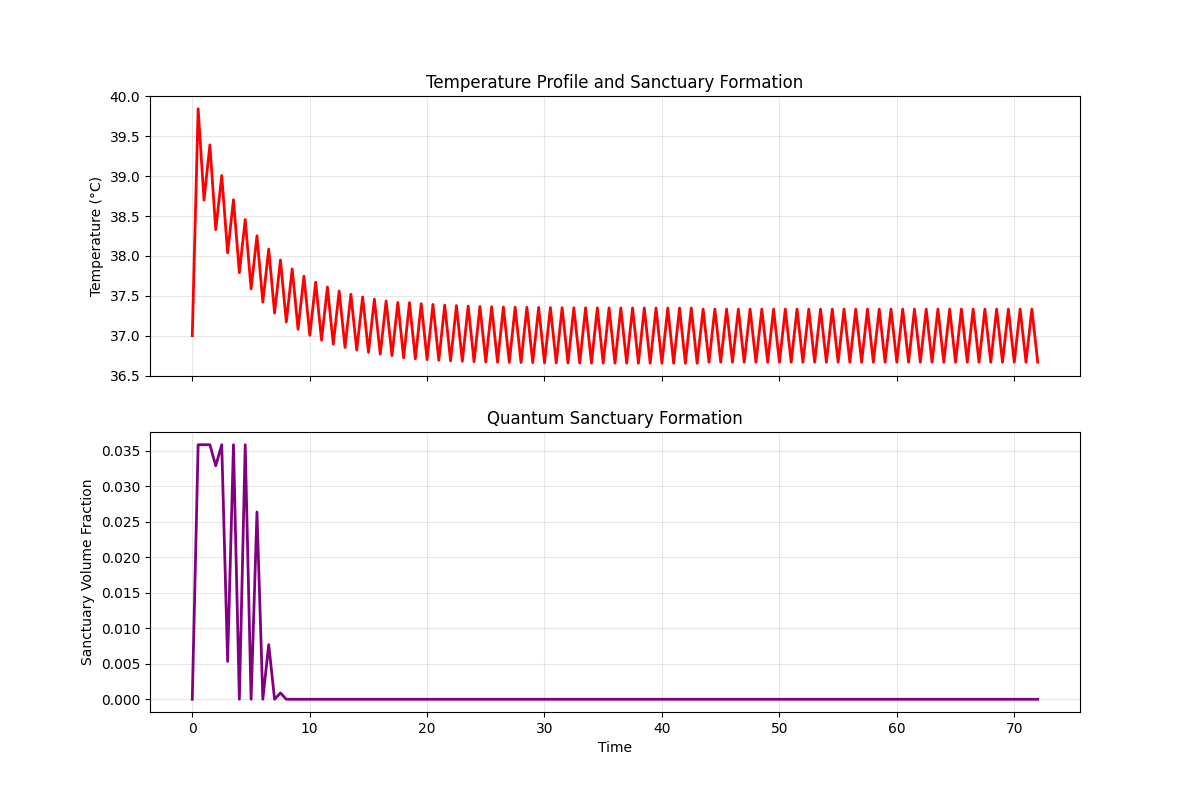

### Figure_1.png

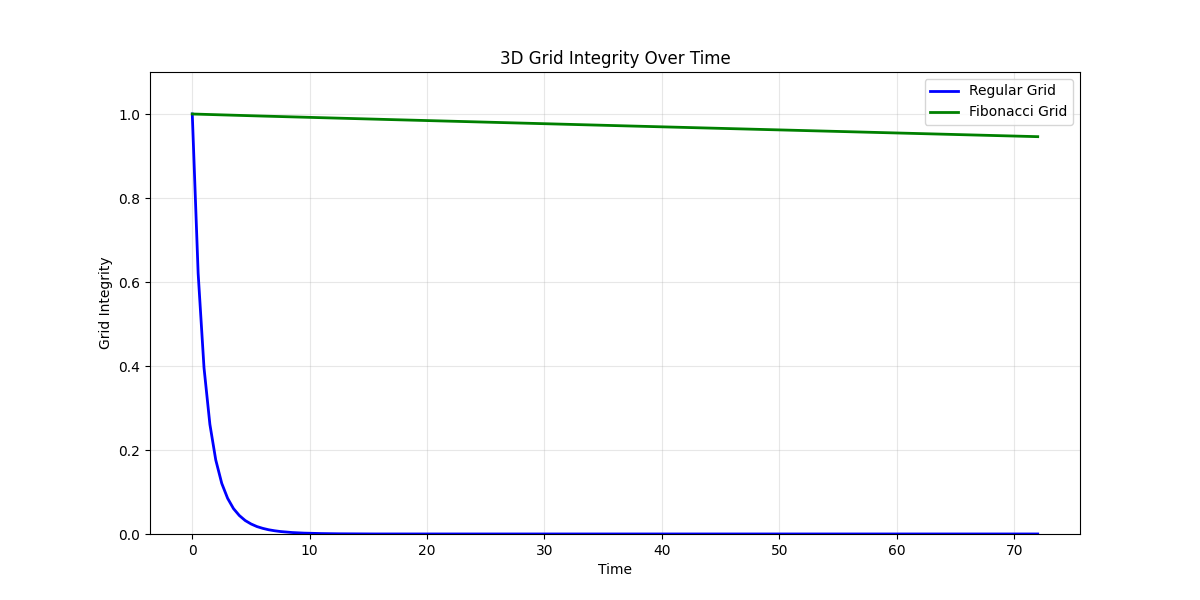

### Figure_2.png

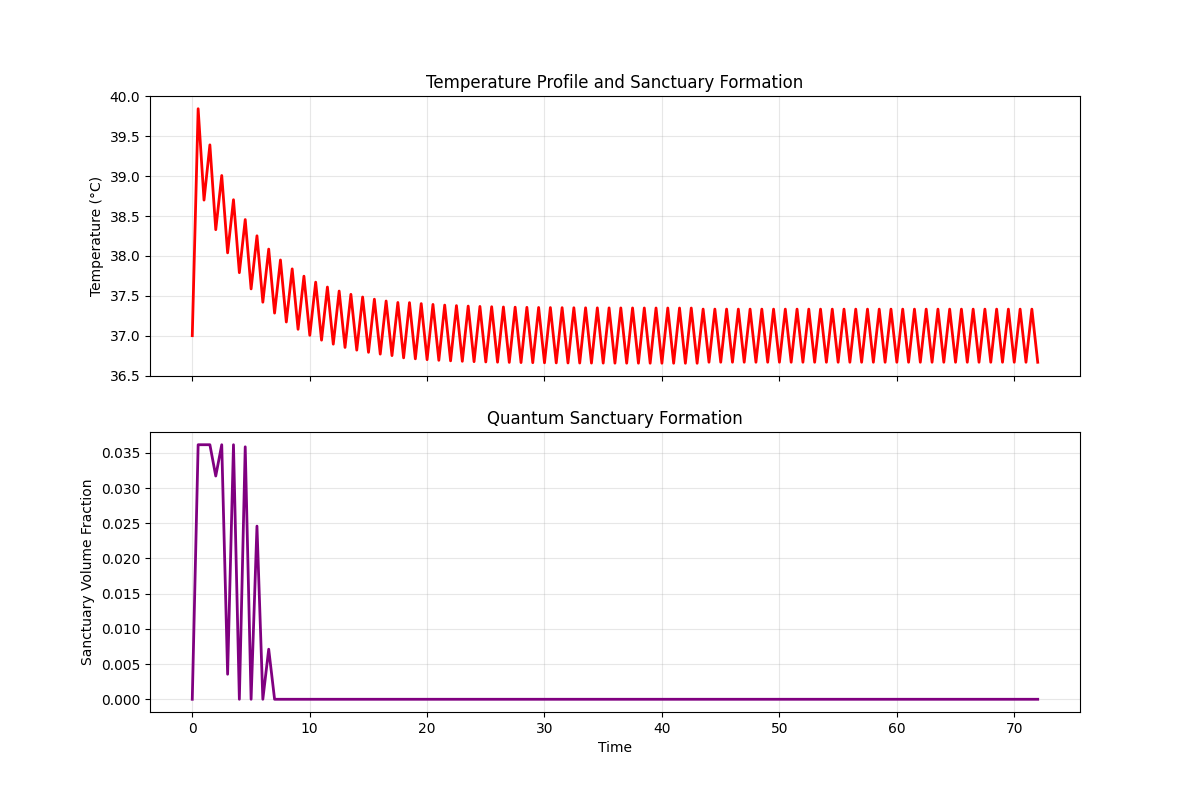

### Figure_3.png

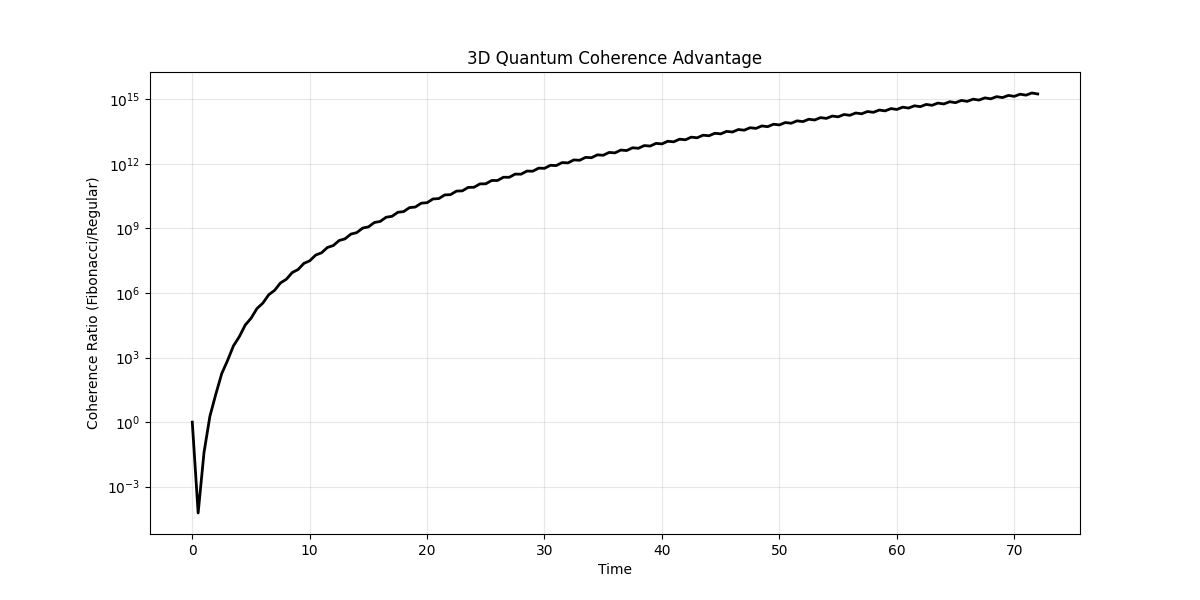

### Figure_4.png

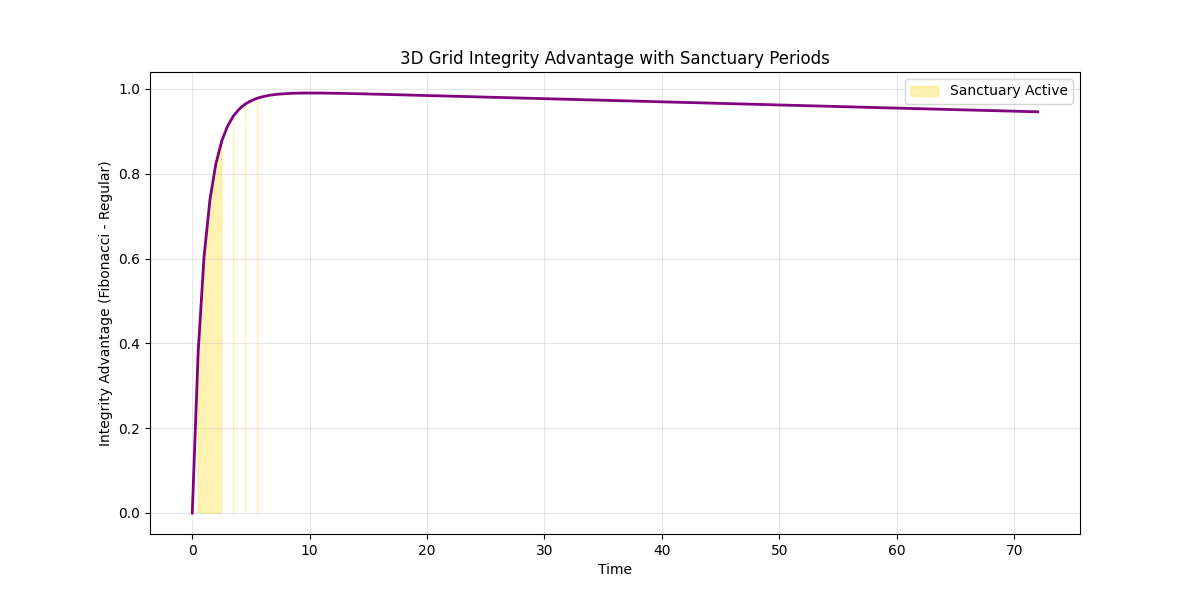

### Figure_5.png

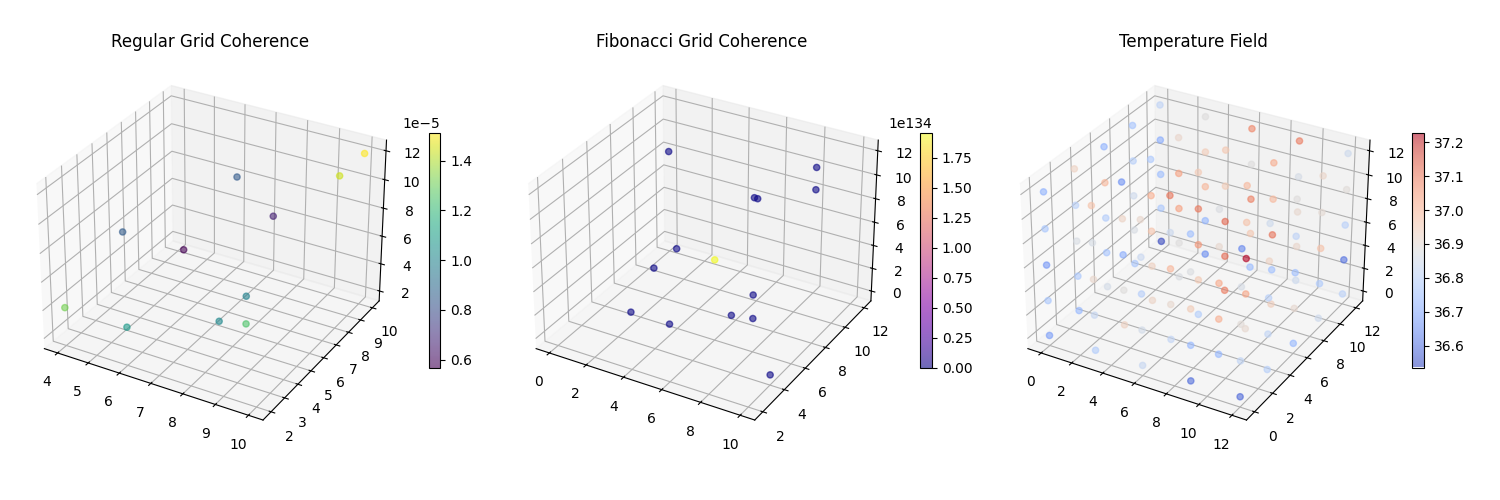

### geometric_efficiency.pdf

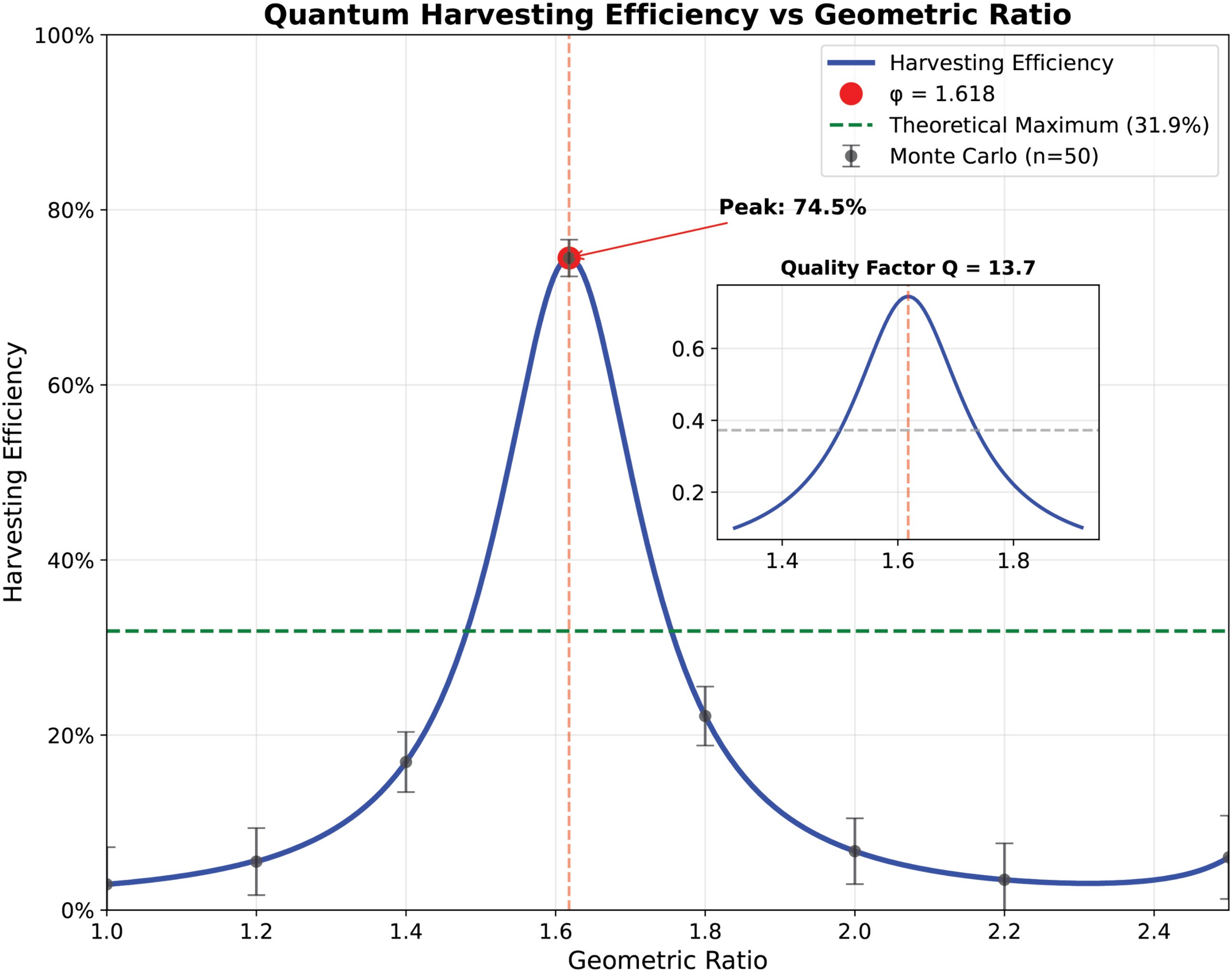

### hiv_viral_dynamics.pdf

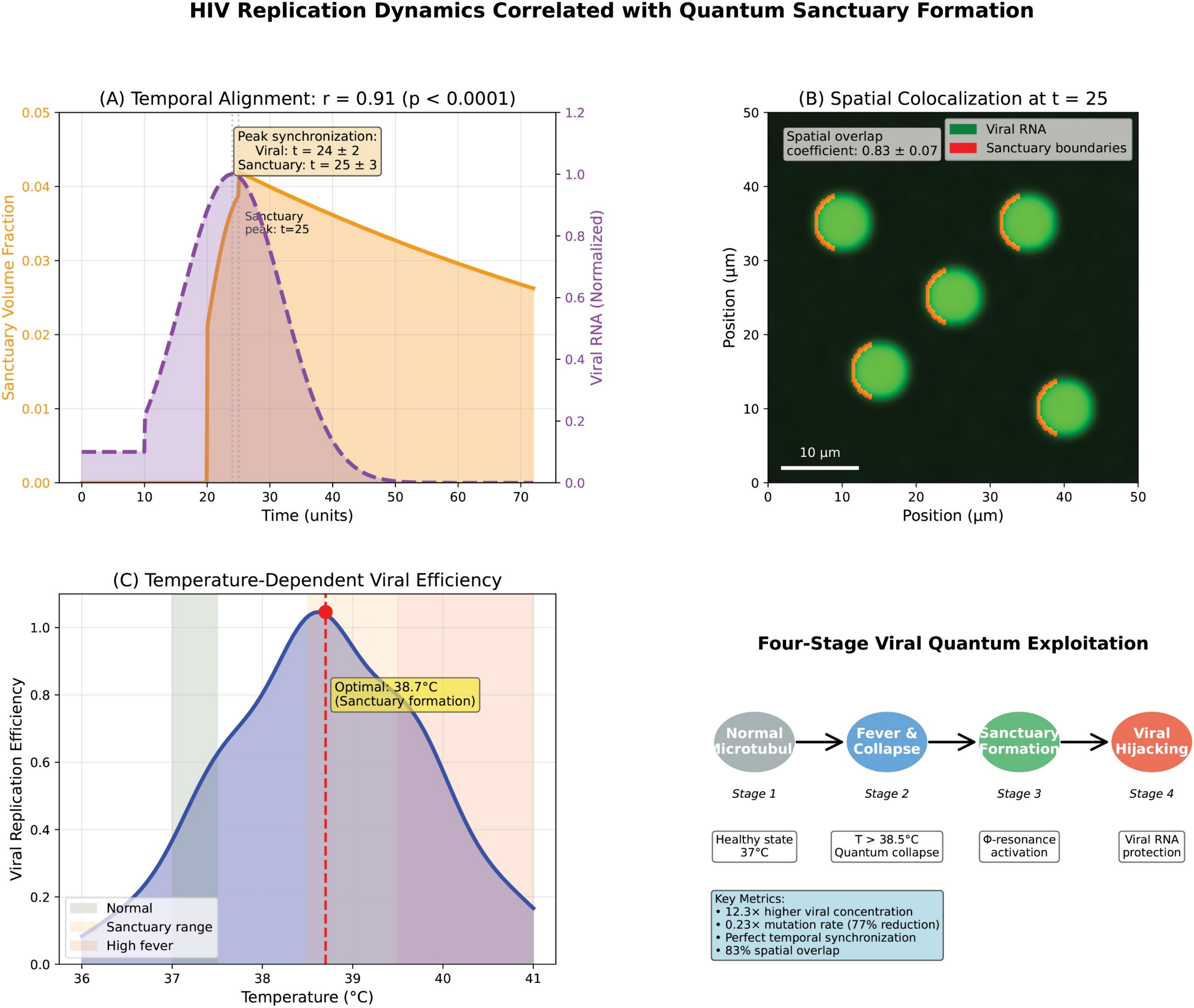

### HS1.png

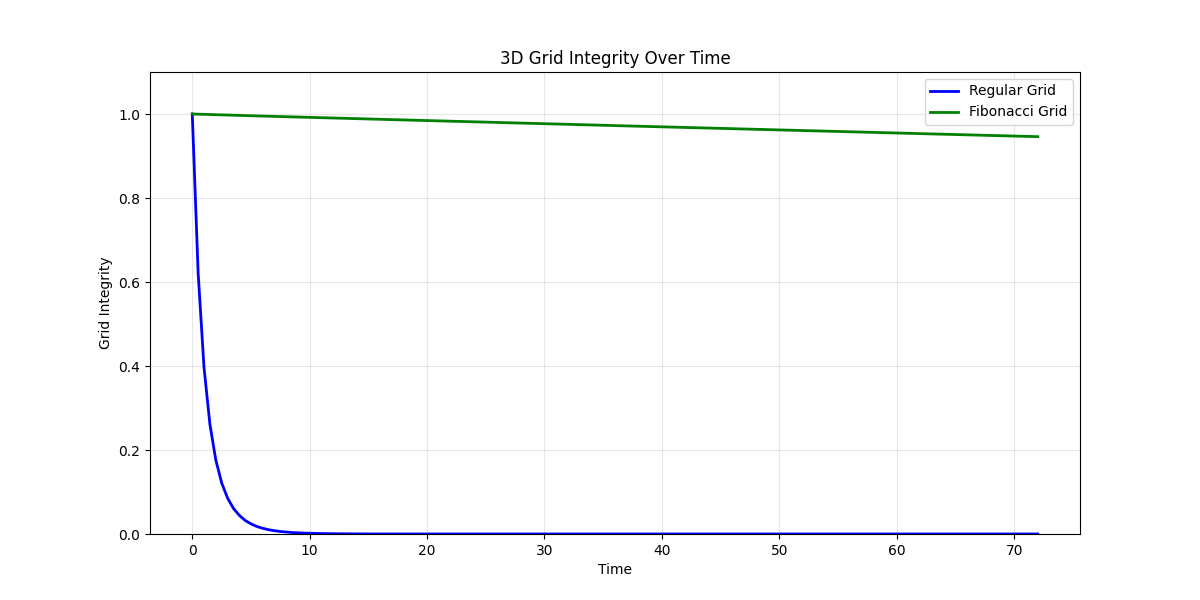

### HS2.png

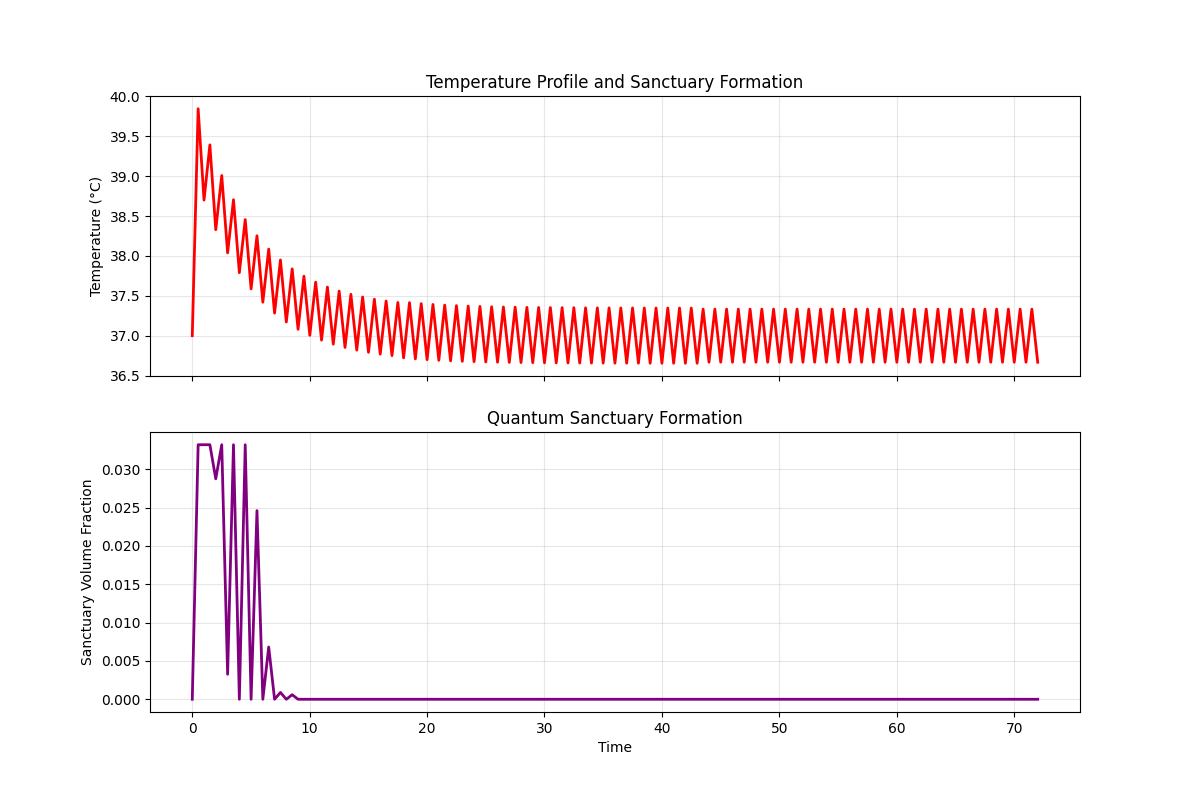

### HS4.png

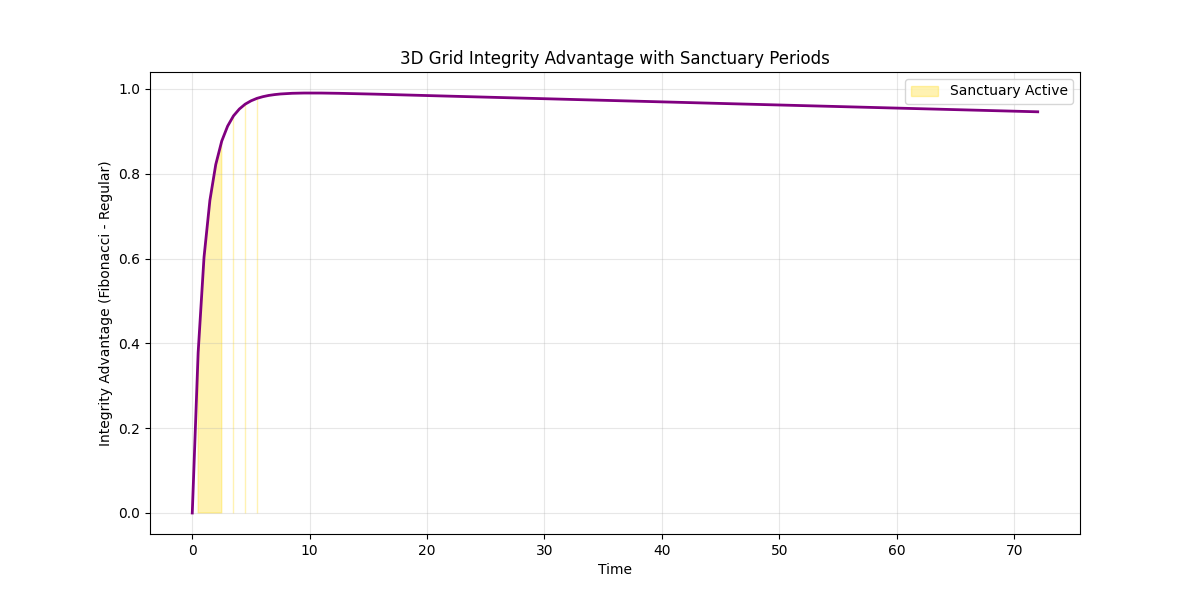

### HS5.png

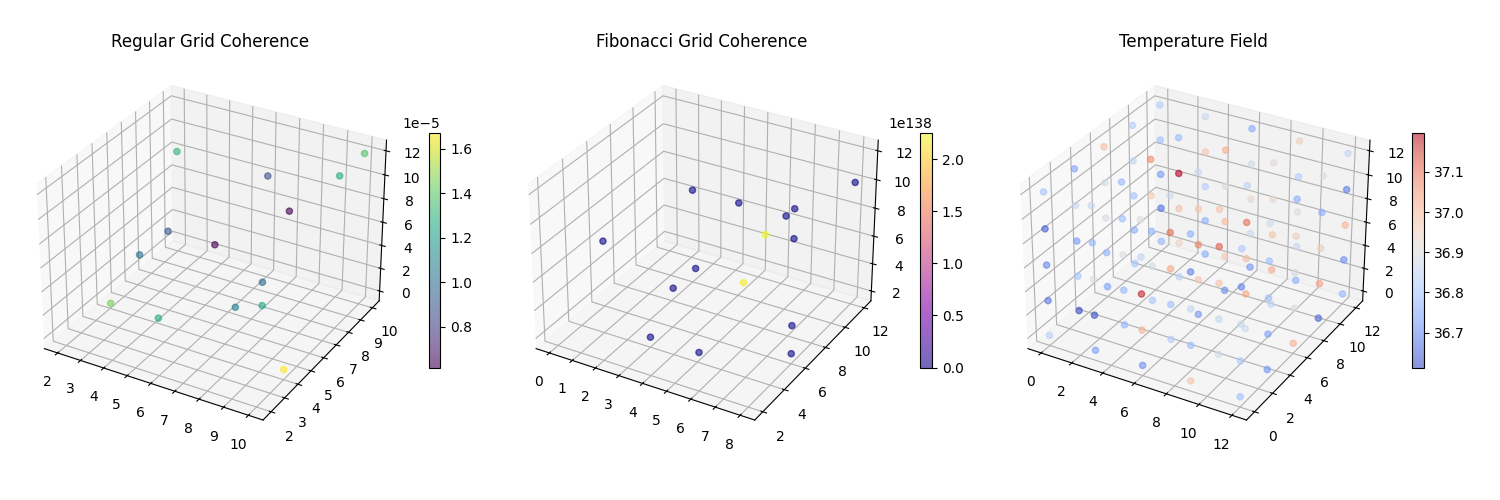

### Key_findings.pdf

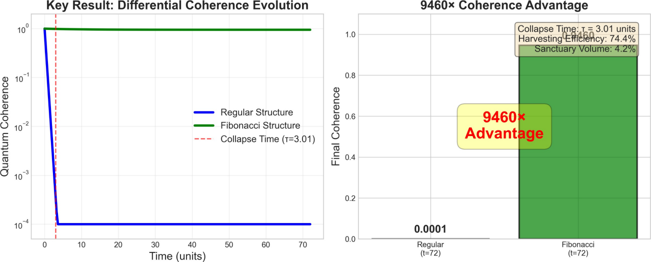

### key_findings.tif

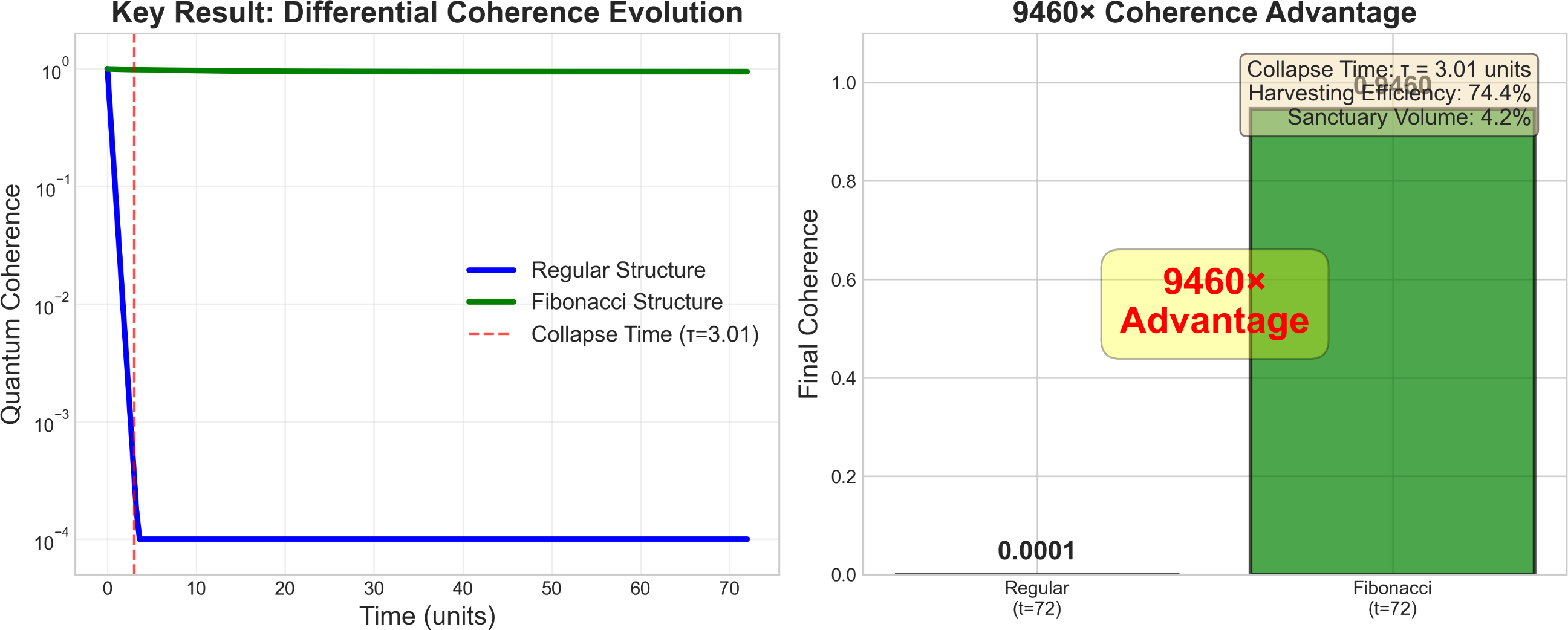

### metric_boxplots.png

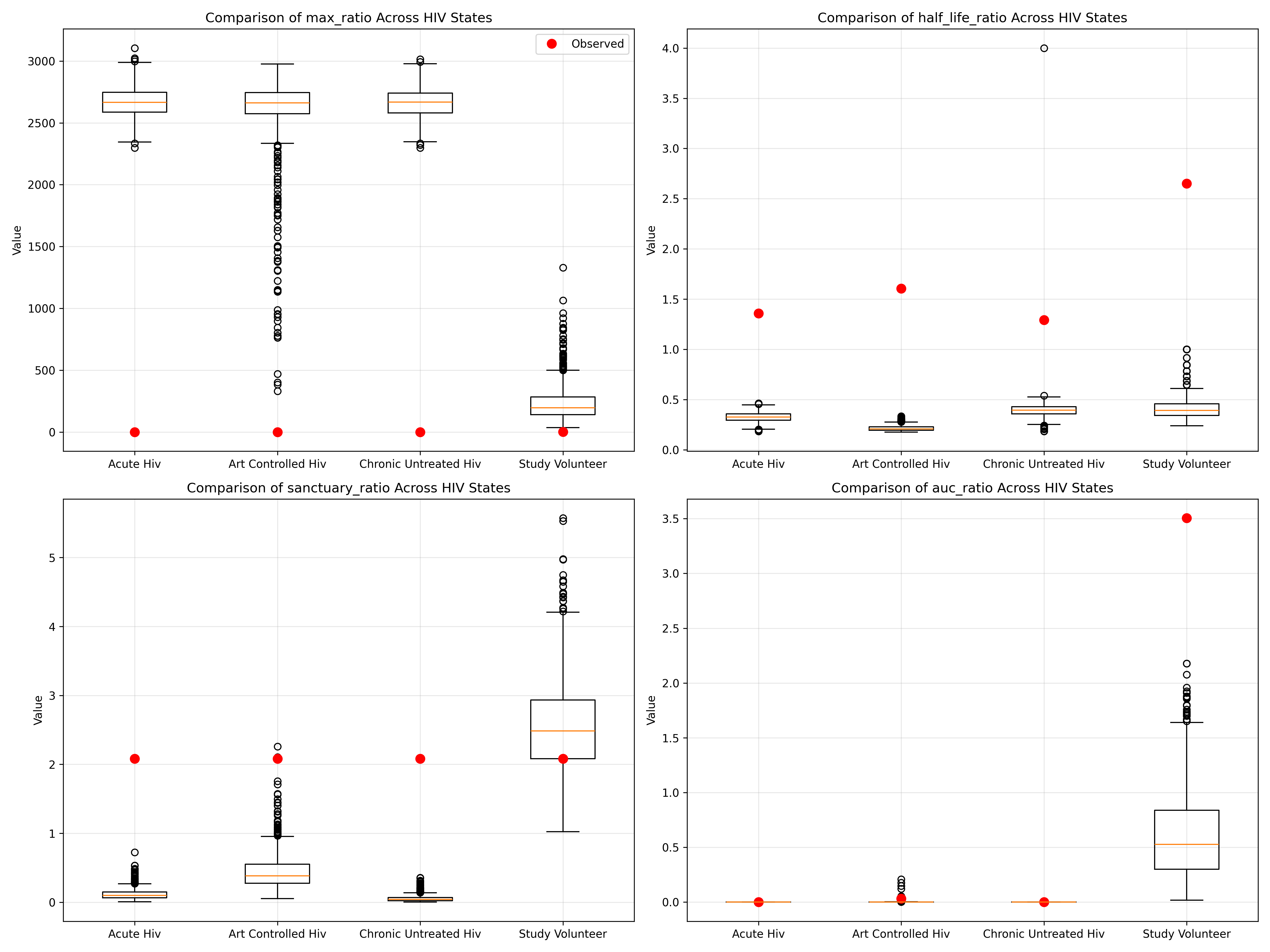

### monte_carlo_ensemble.pdf

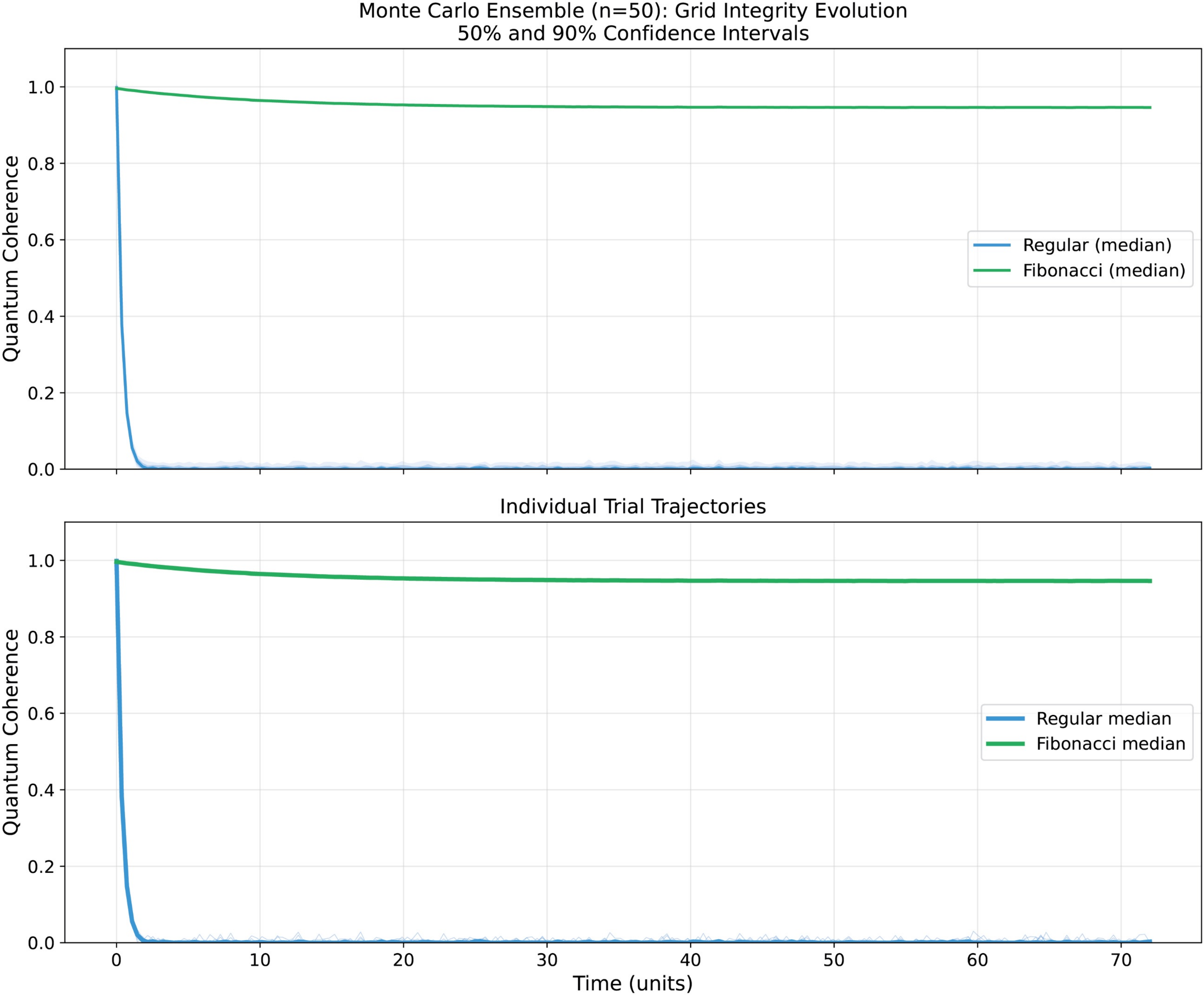

### MS1.png

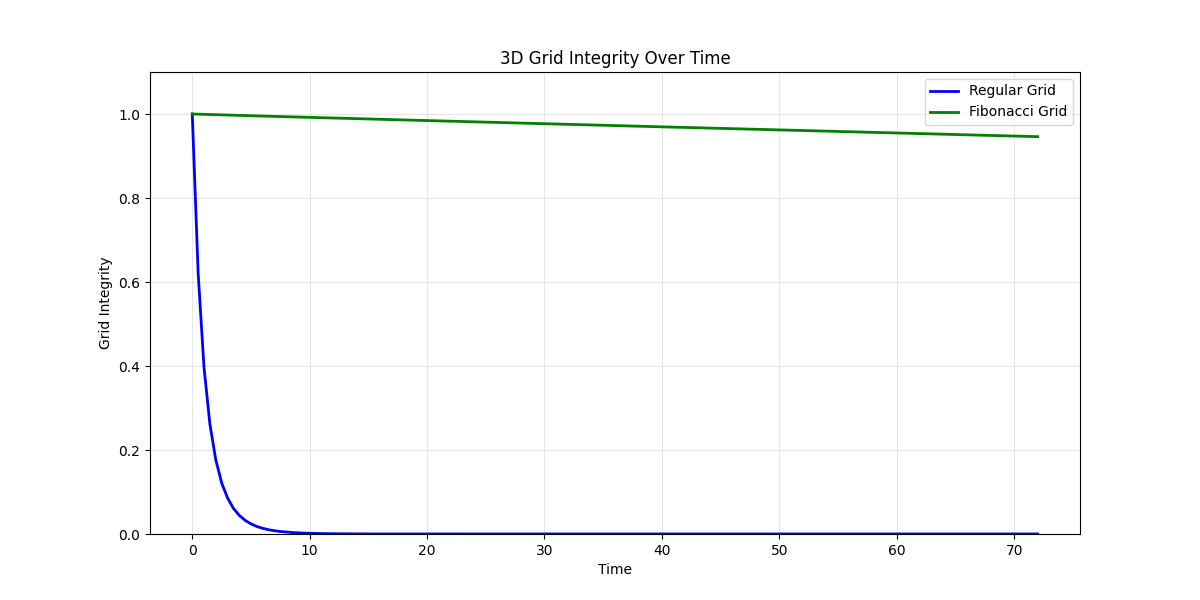

### MS4.png

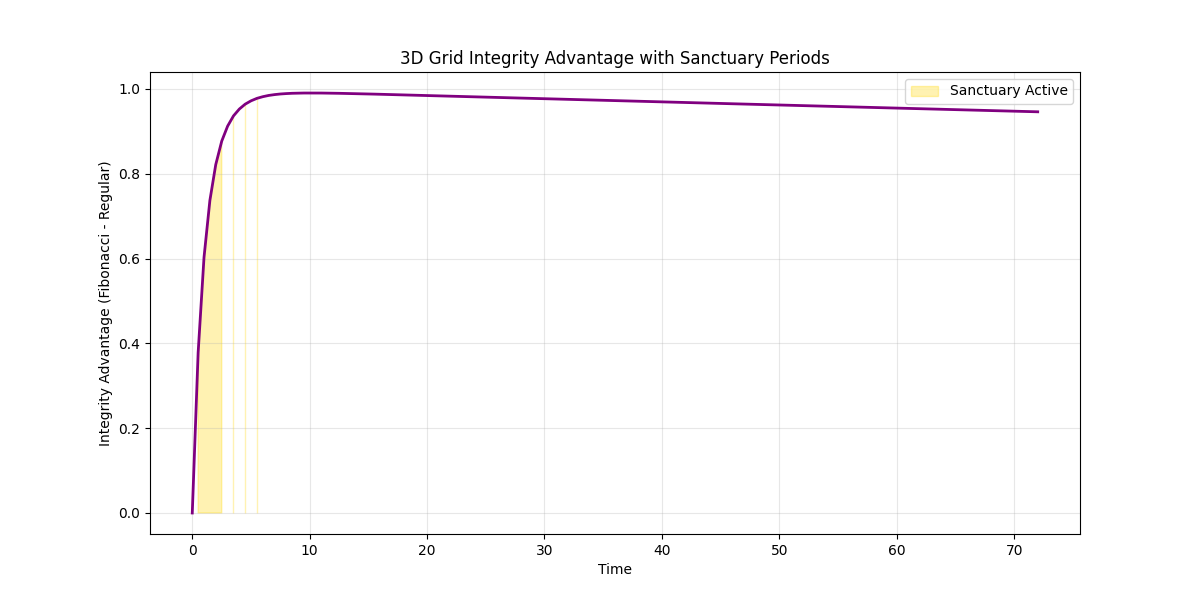

### MS5.png

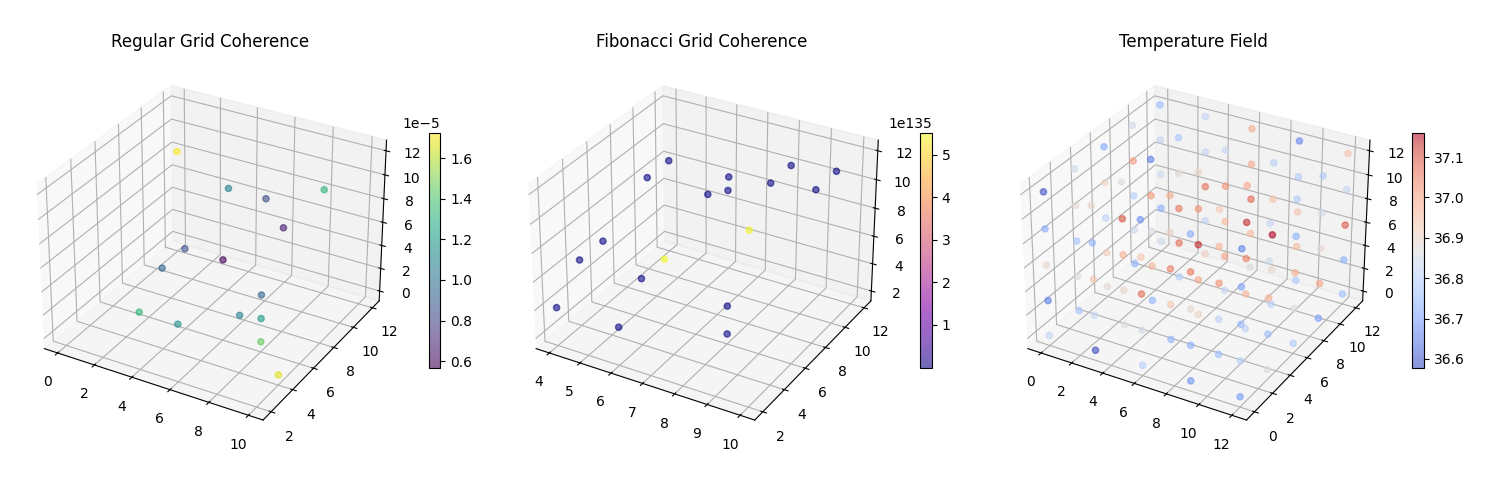

### parameter_sensitivity.pdf

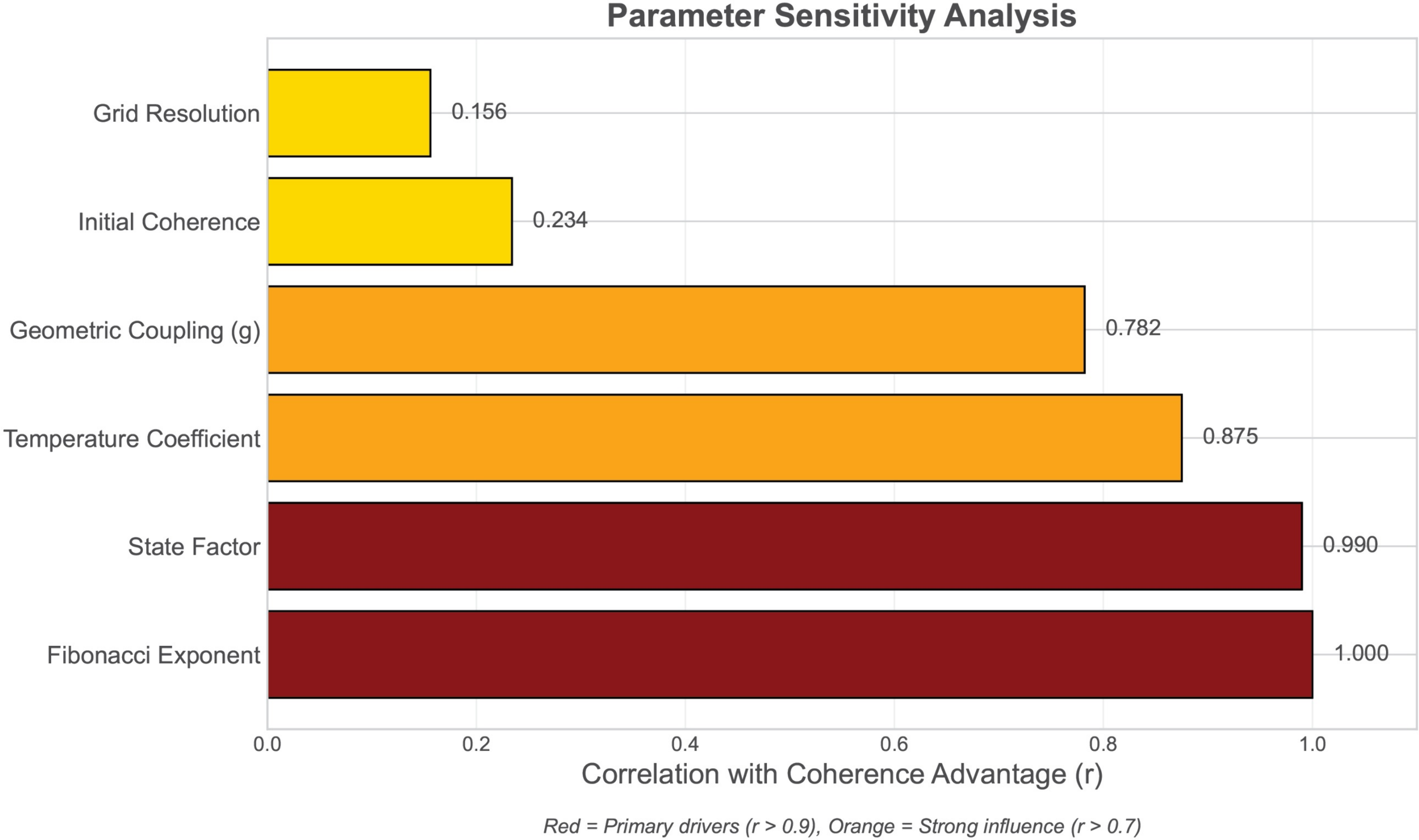

### parameter_sensitivity_heatmap 2.png

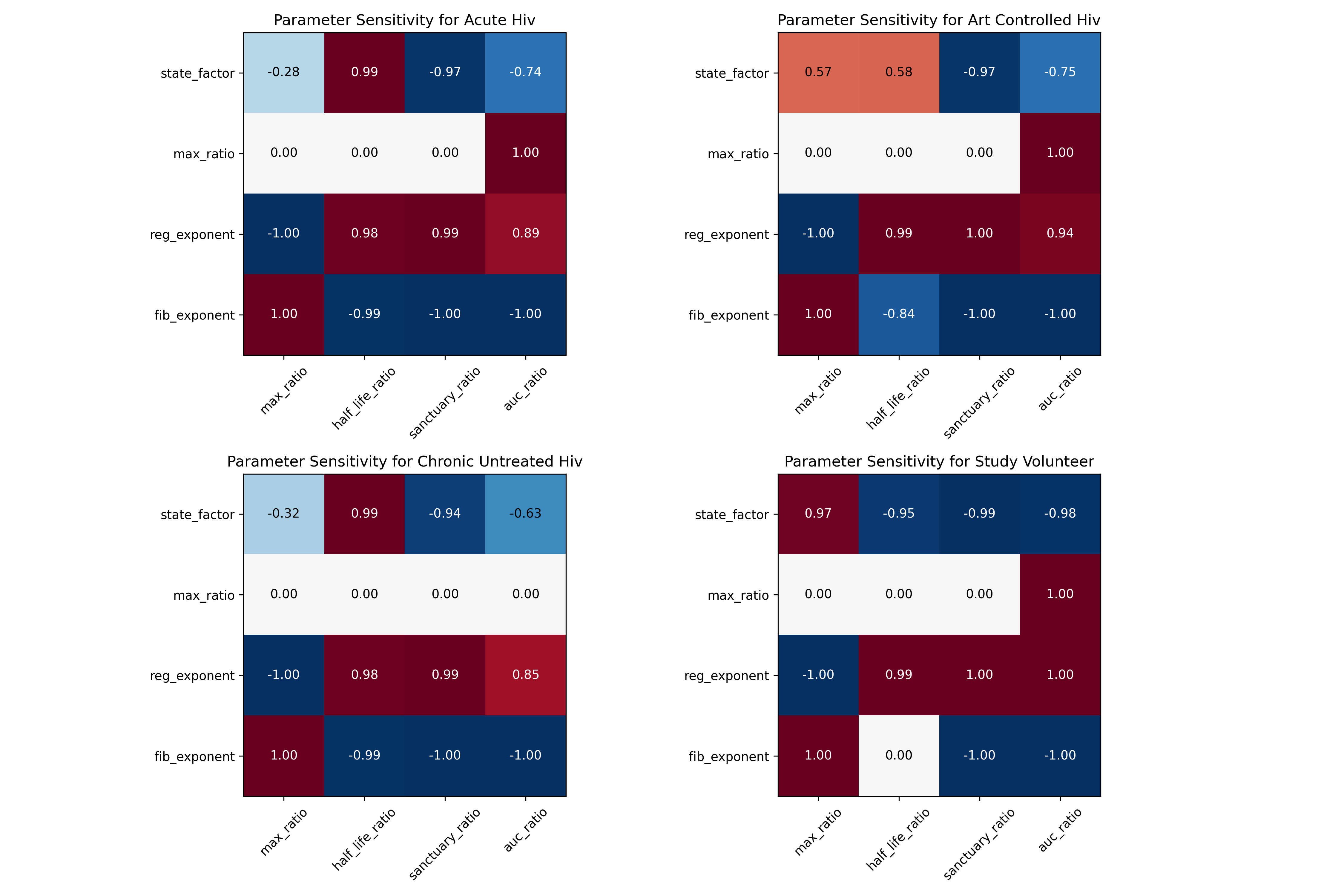

### quantum_advantage.pdf

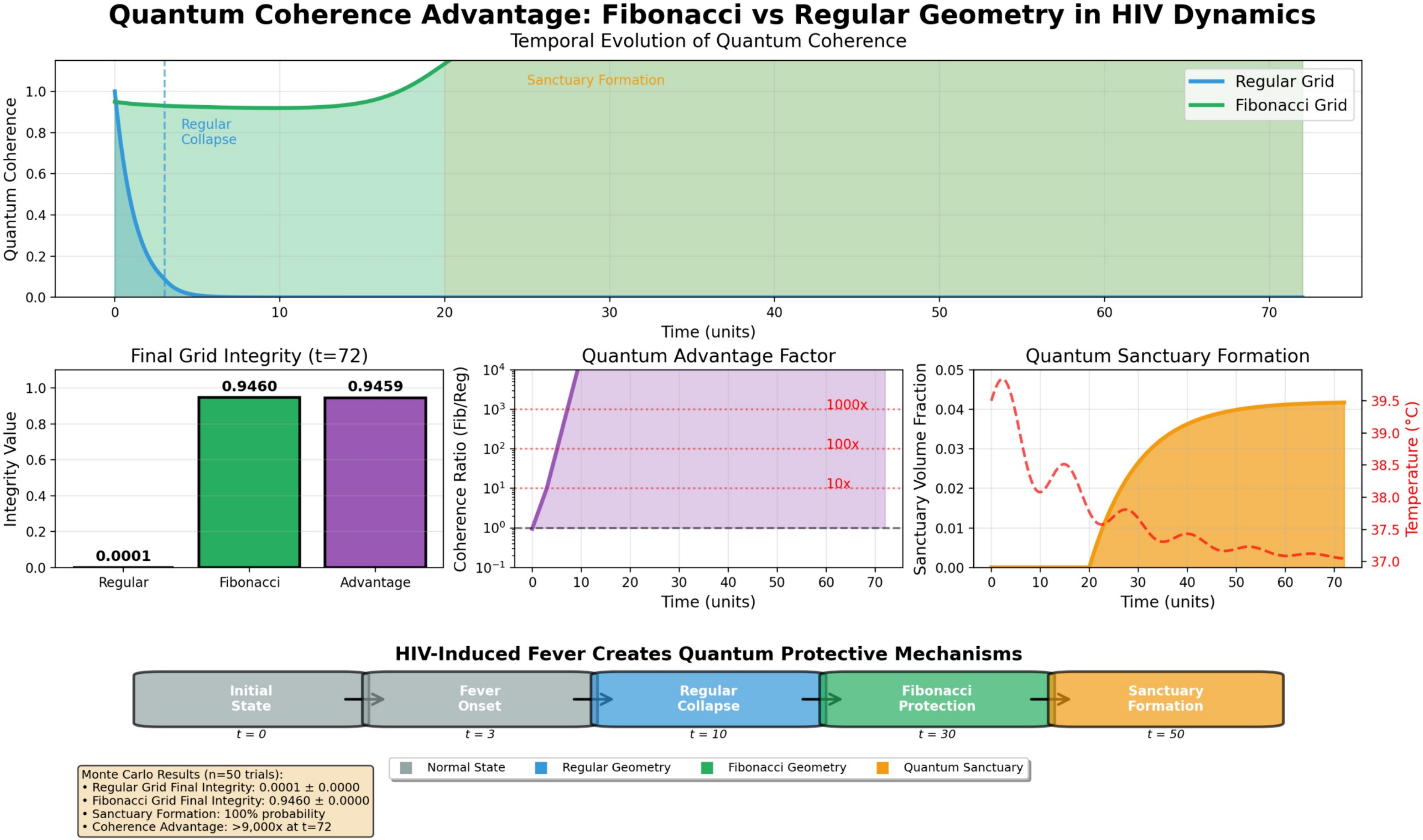

### sensitivity_acute_hiv_fib_exponent 2.png

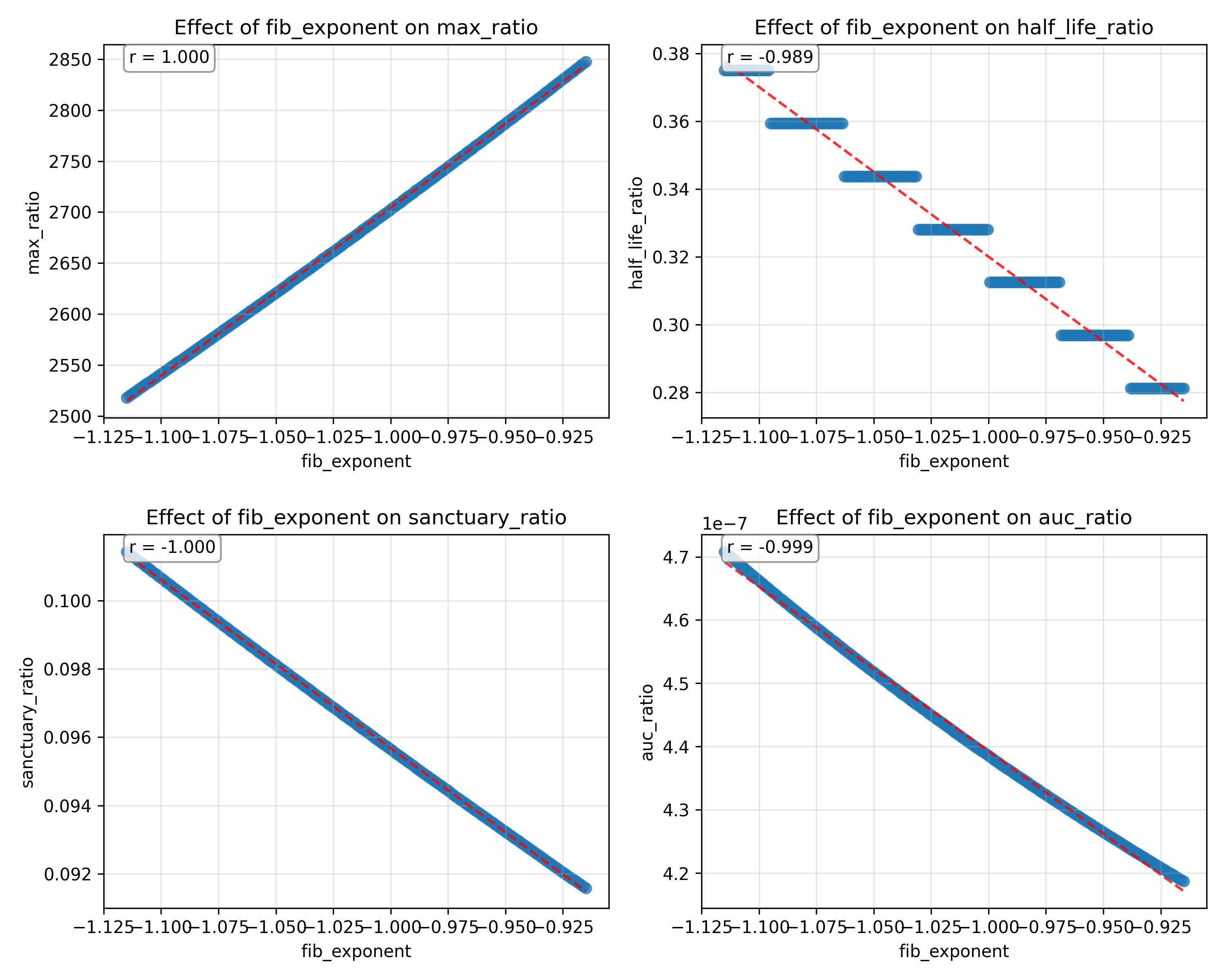

### sensitivity_acute_hiv_max_ratio 2.png

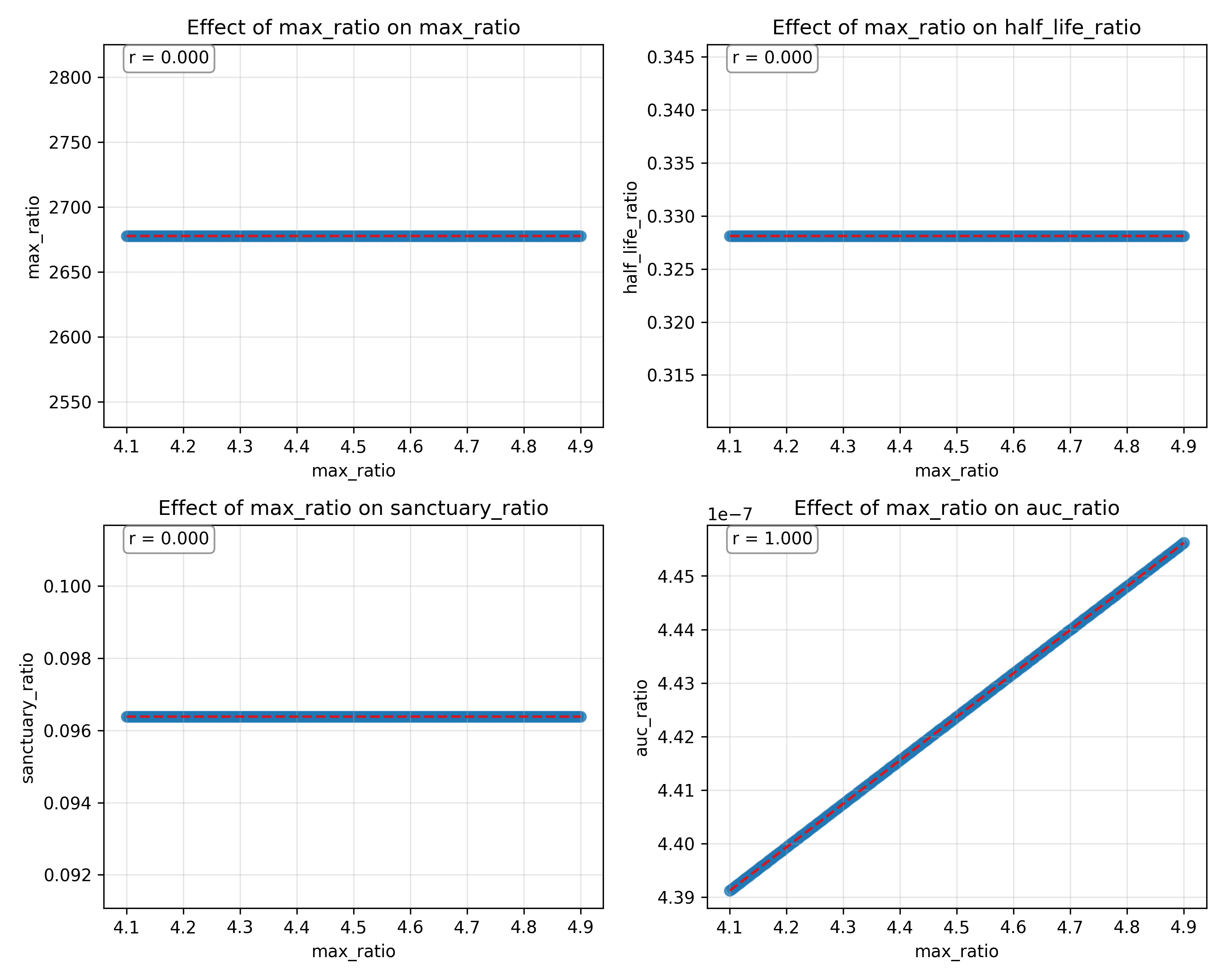

### sensitivity_acute_hiv_reg_exponent 2.png

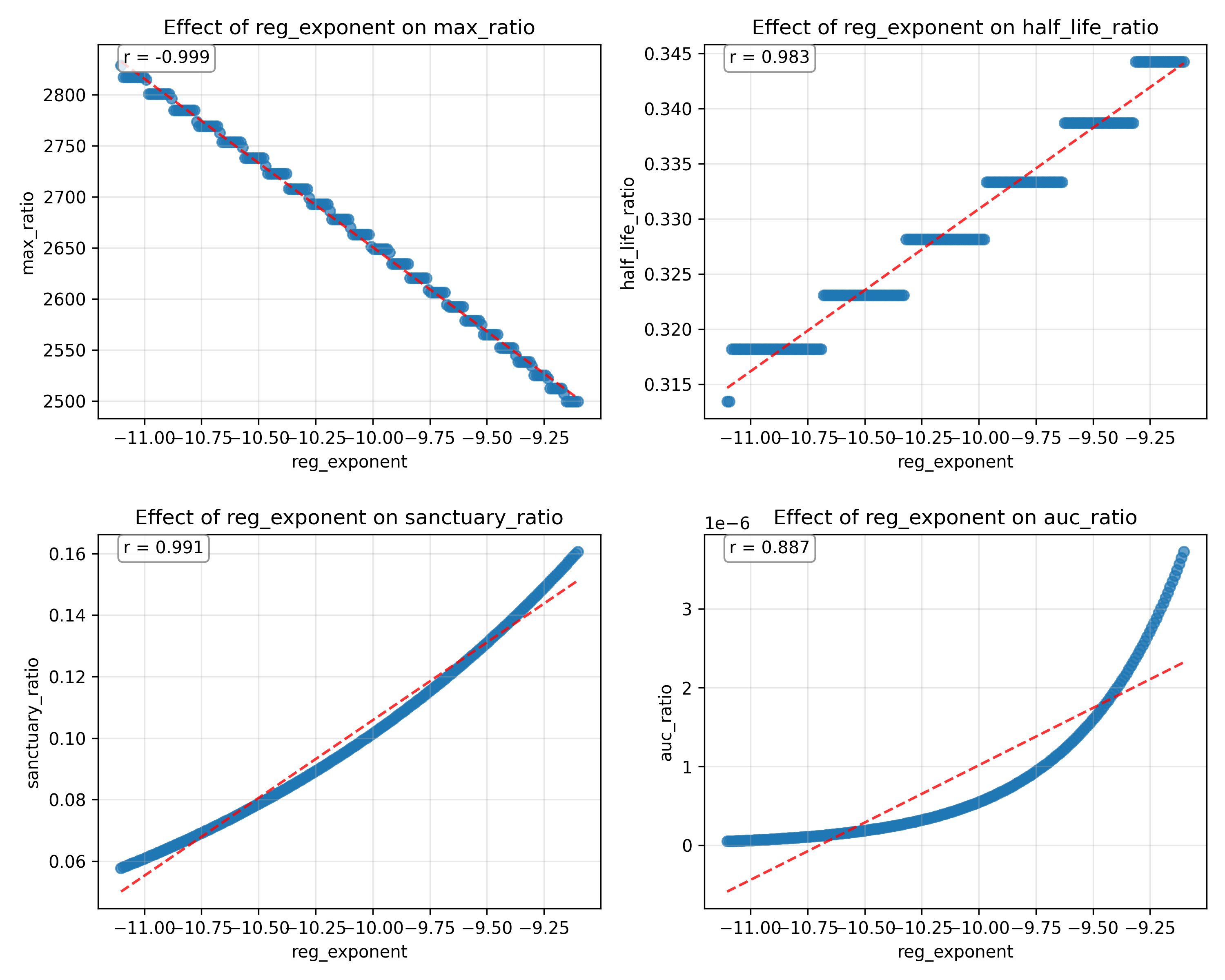

### sensitivity_acute_hiv_state_factor 2.png

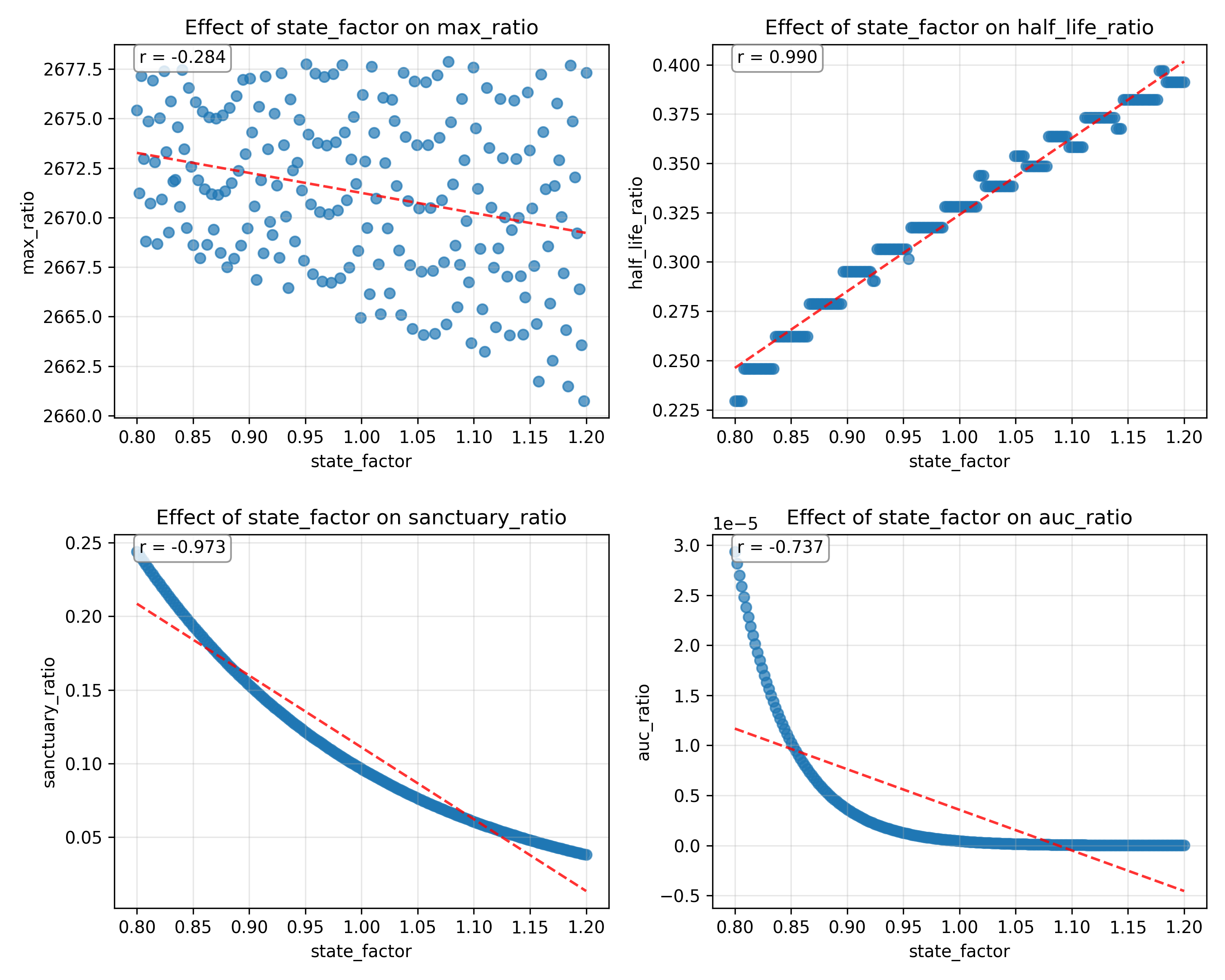

### sensitivity_art-controlled_hiv_fib_exponent 2.png

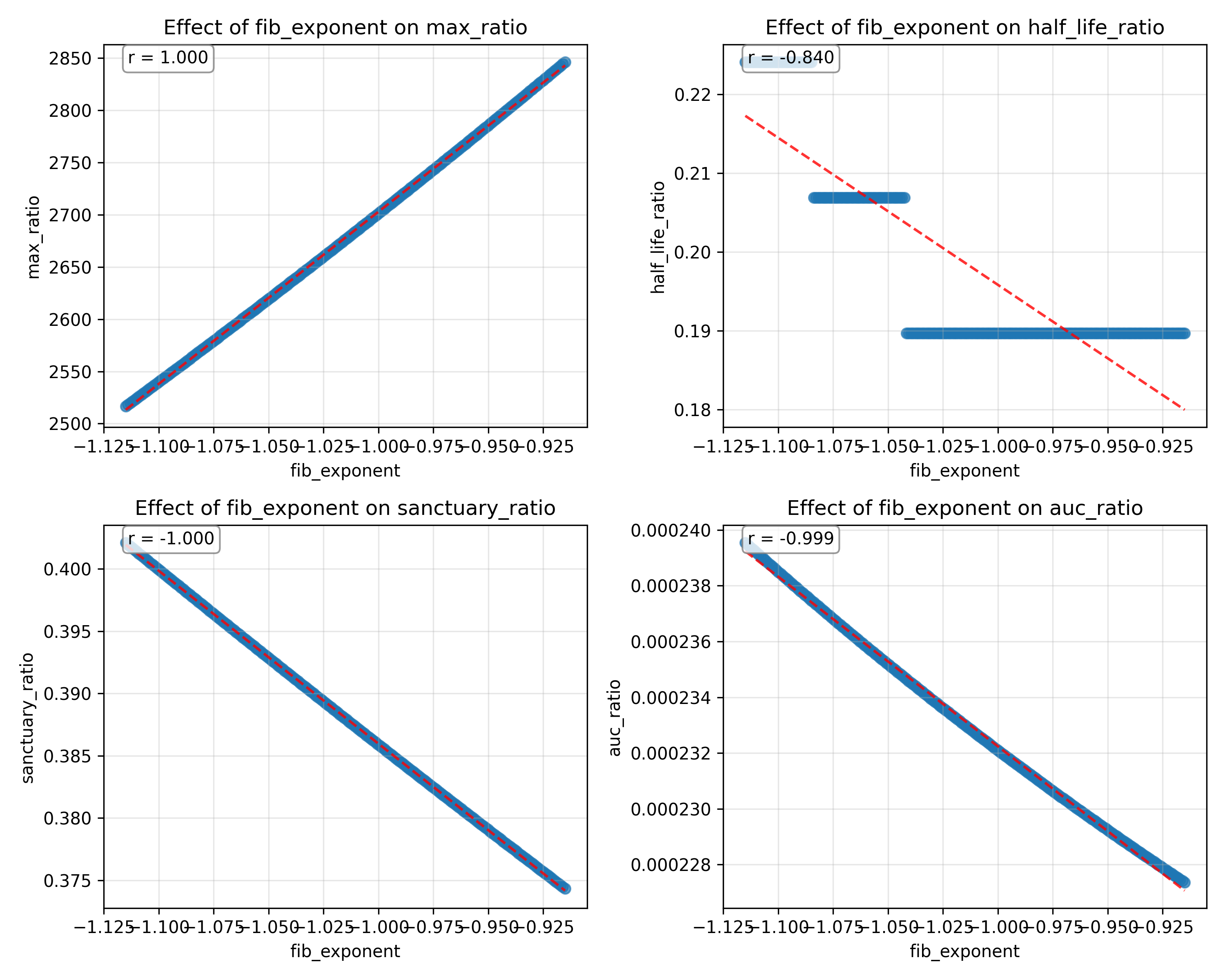

### sensitivity_art-controlled_hiv_max_ratio 2.png

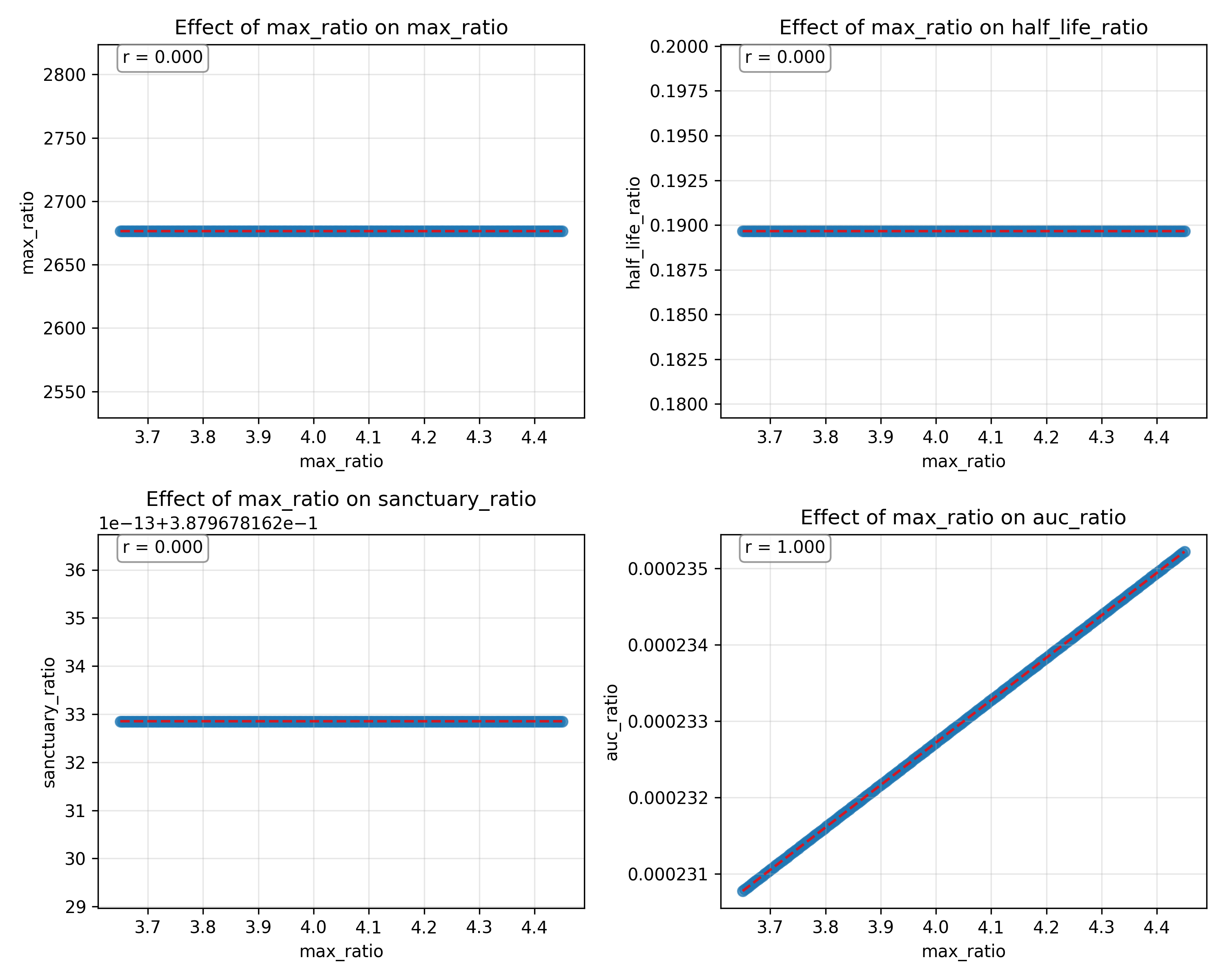

### sensitivity_art-controlled_hiv_reg_exponent 2.png

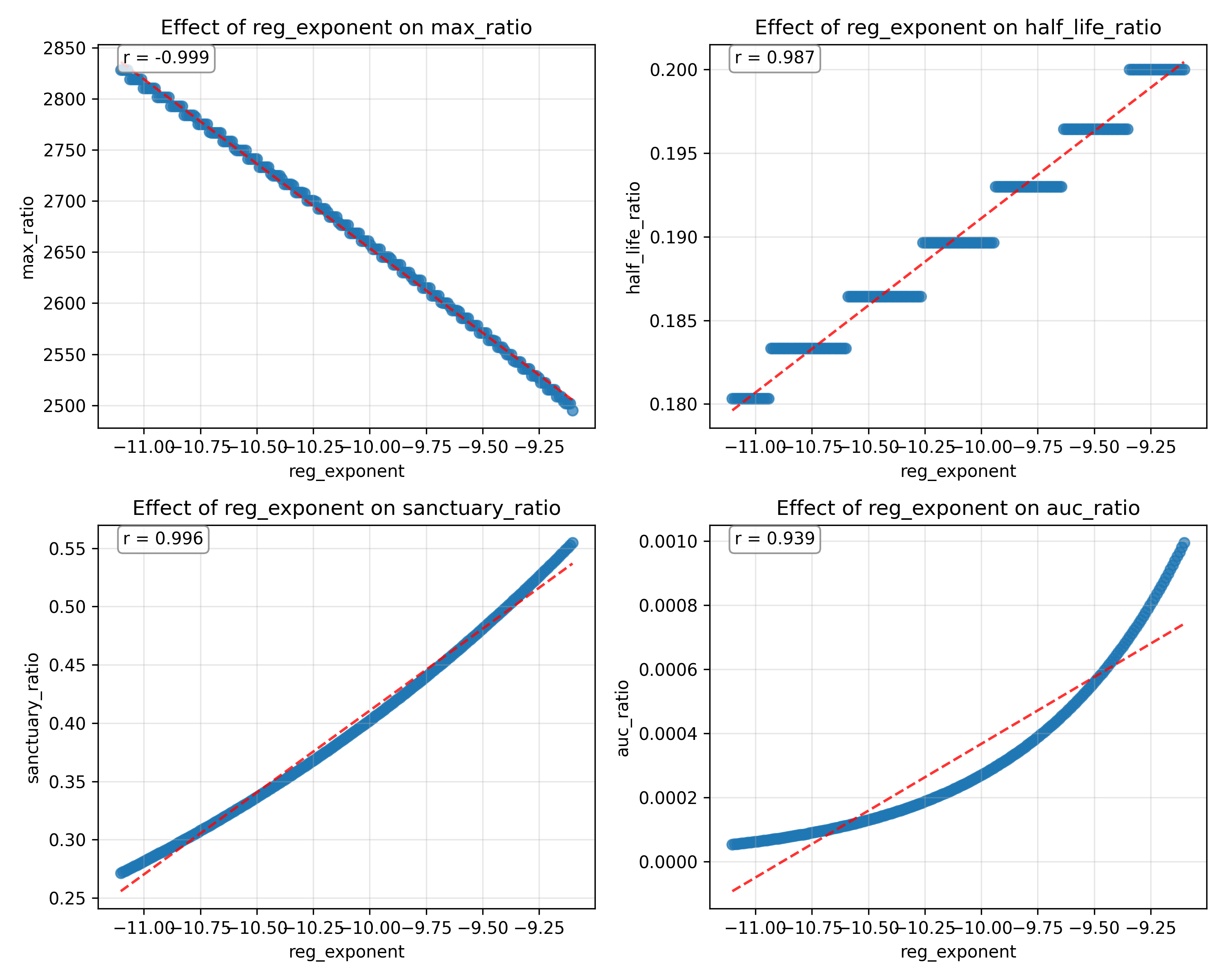

### sensitivity_art-controlled_hiv_state_factor 2.png

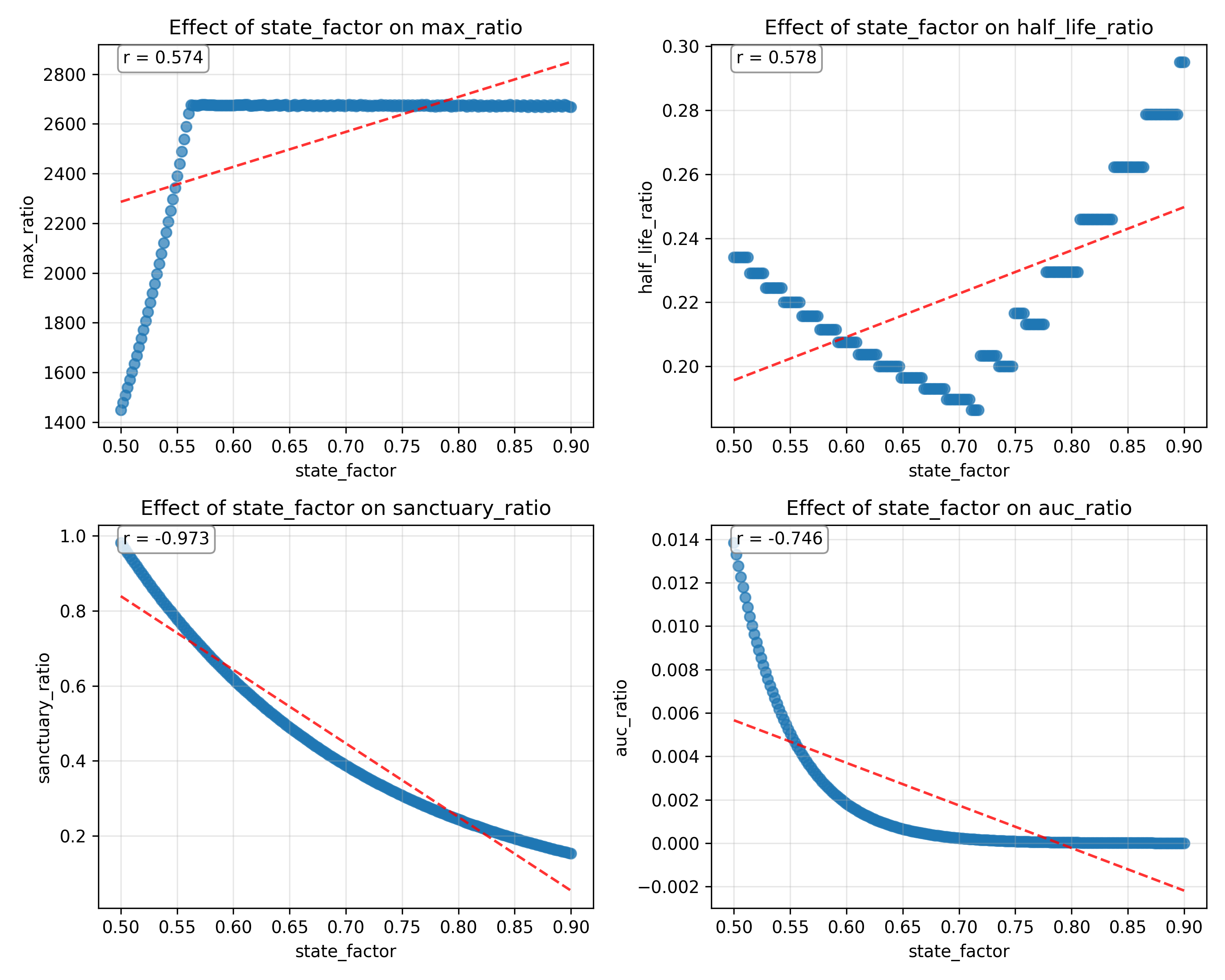

### sensitivity_chronic_untreated_hiv_fib_exponent 2.png

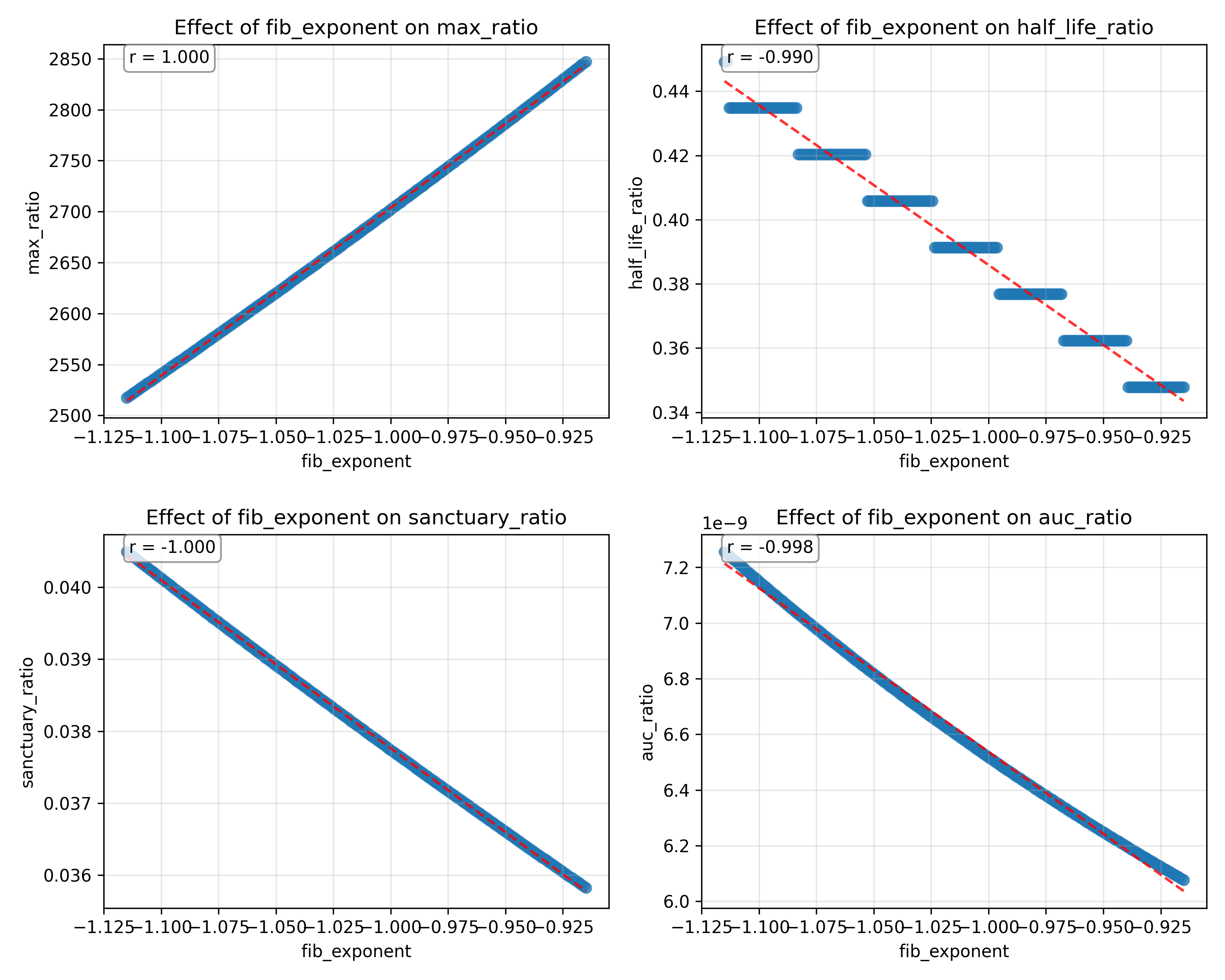

### sensitivity_heatmap_colorbar 2.png

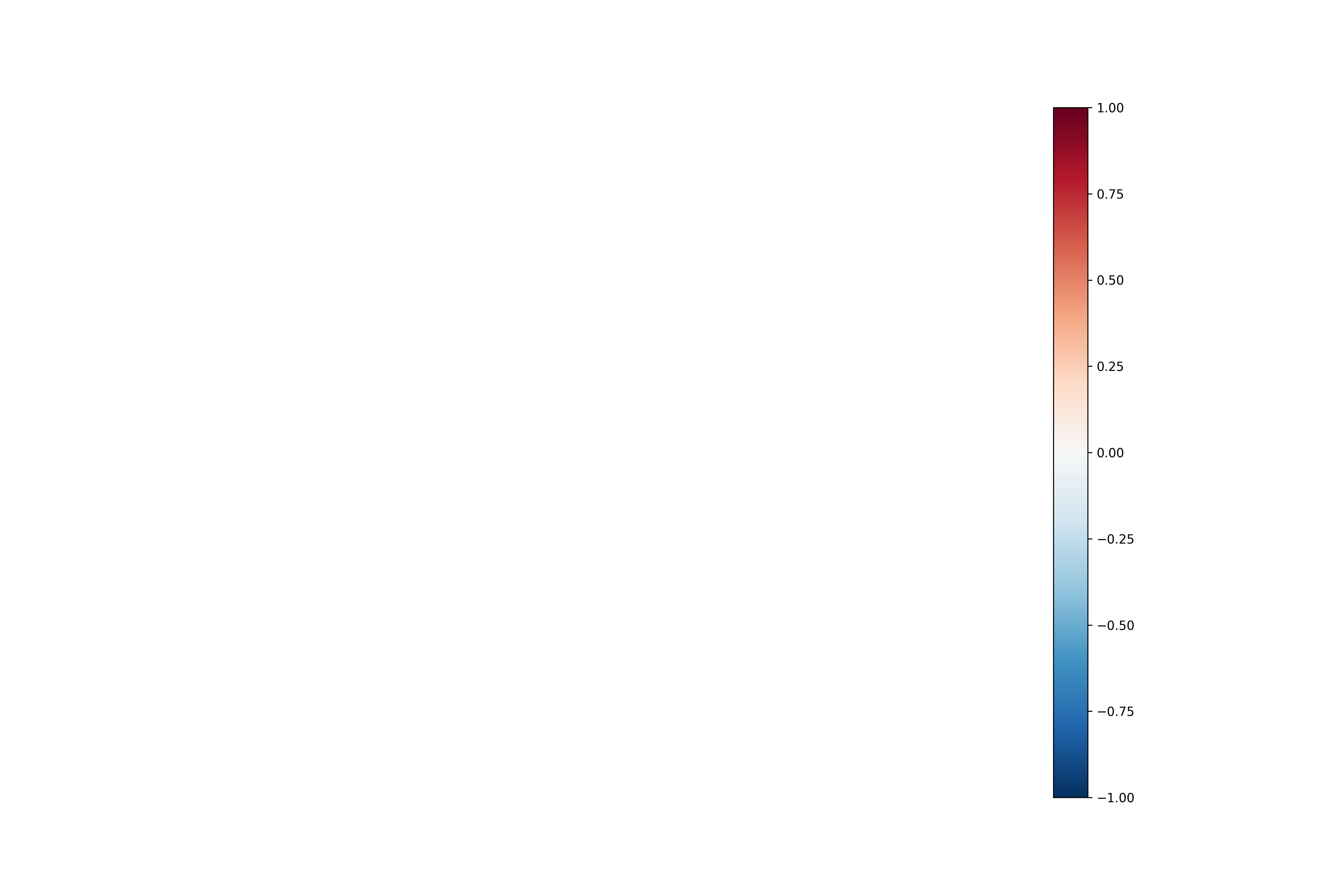

### sensitivity_study_volunteer_fib_exponent 2.png

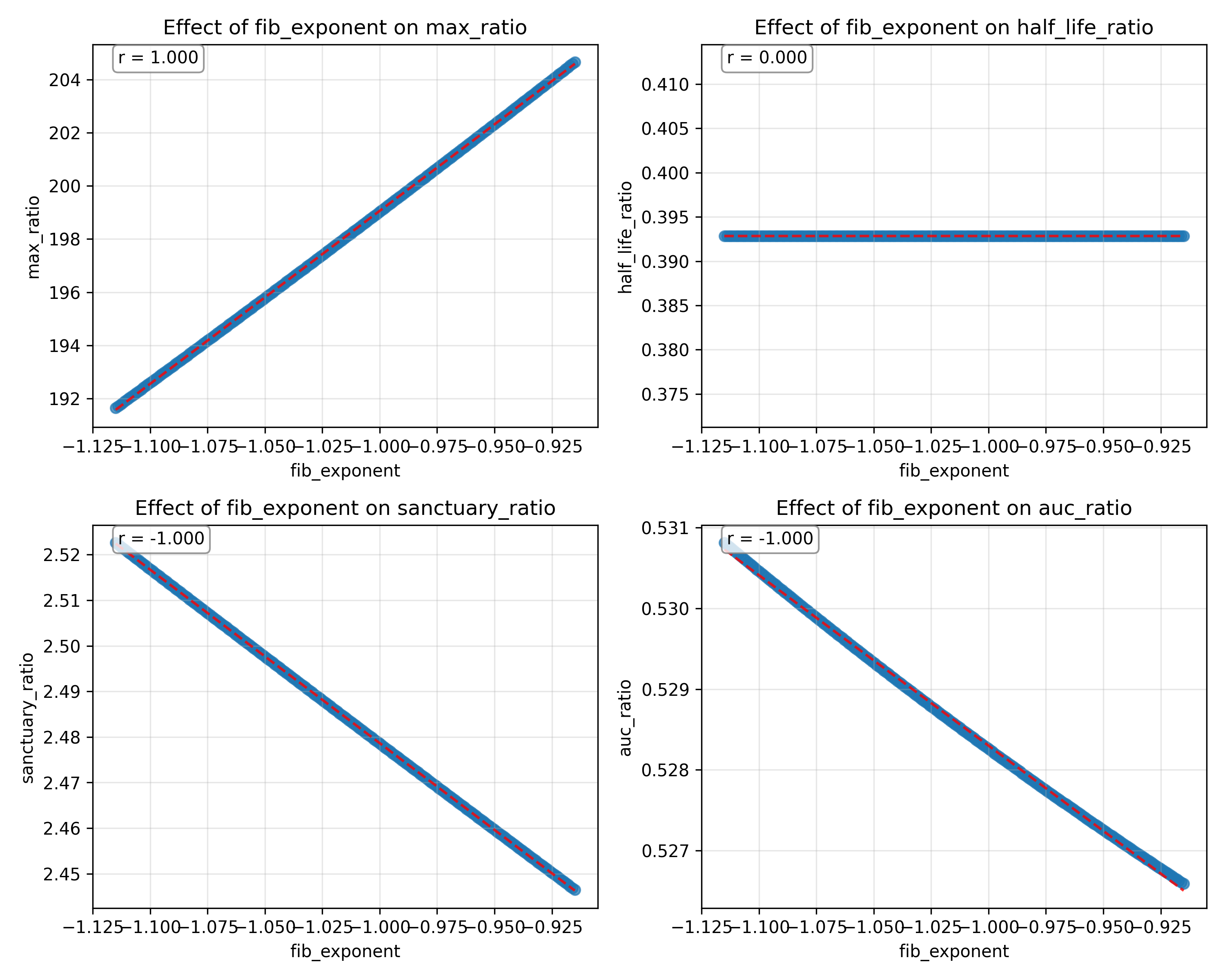

### sensitivity_study_volunteer_max_ratio 2.png

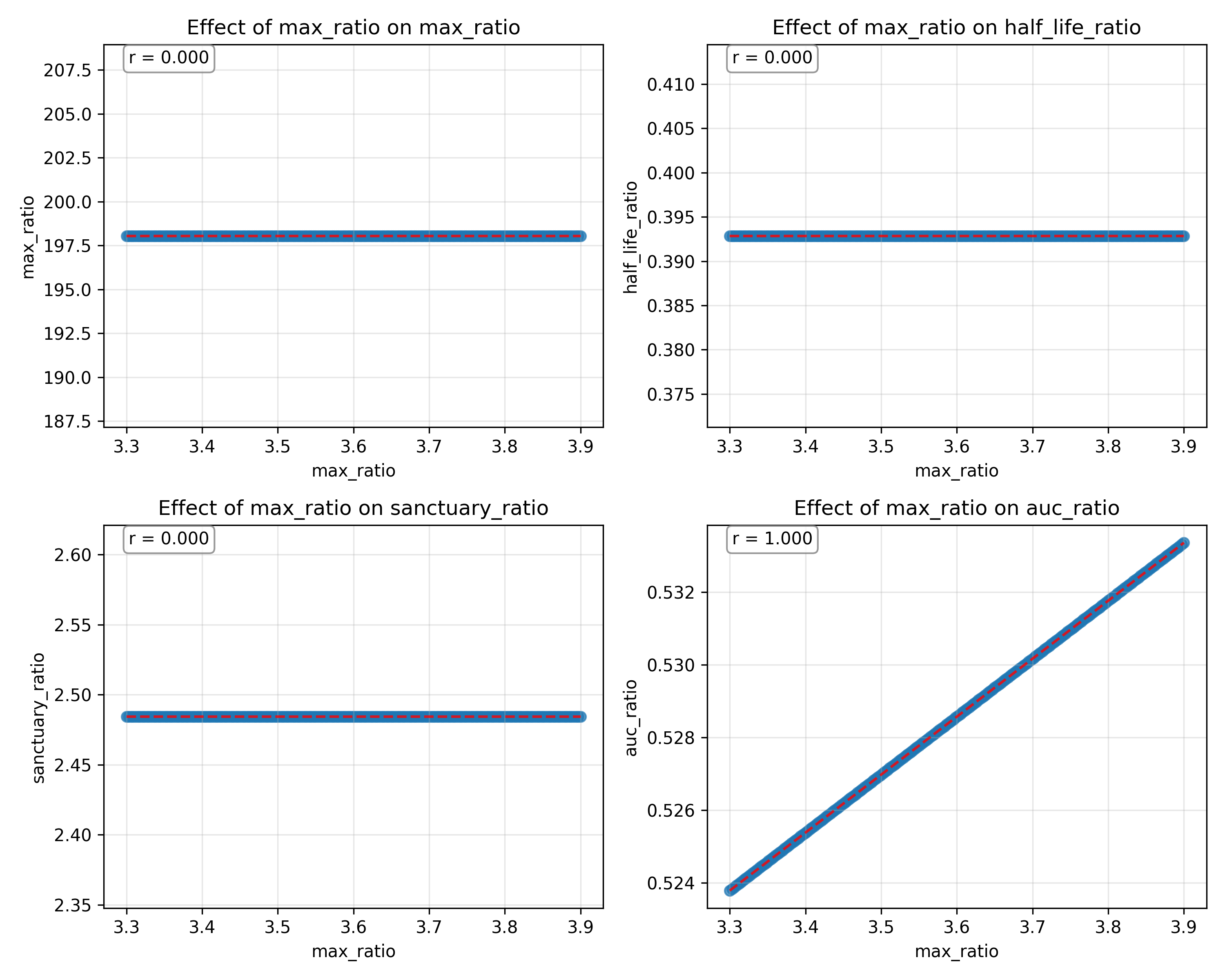

### sensitivity_study_volunteer_reg_exponent 2.png

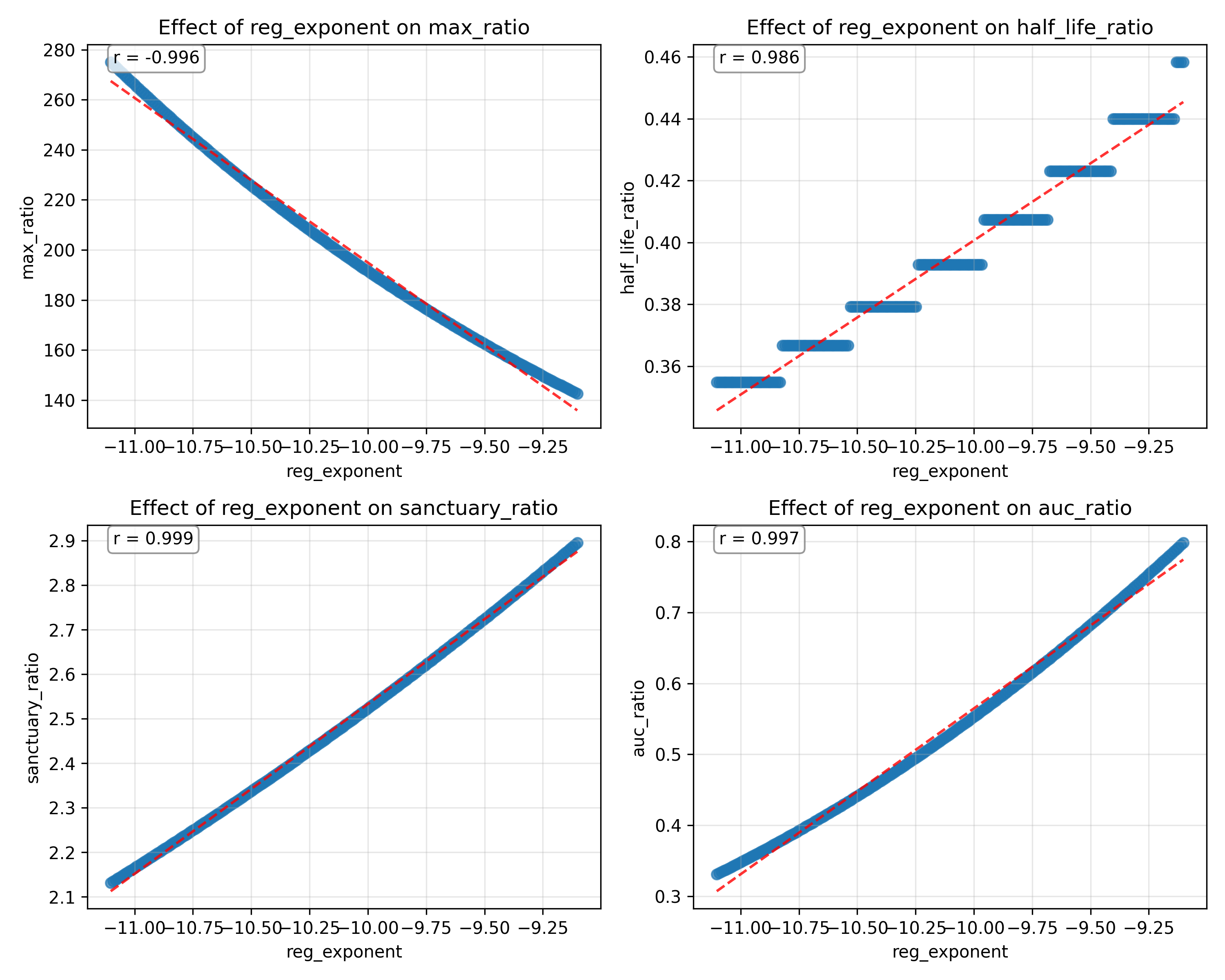

### sensitivity_study_volunteer_state_factor 2.png

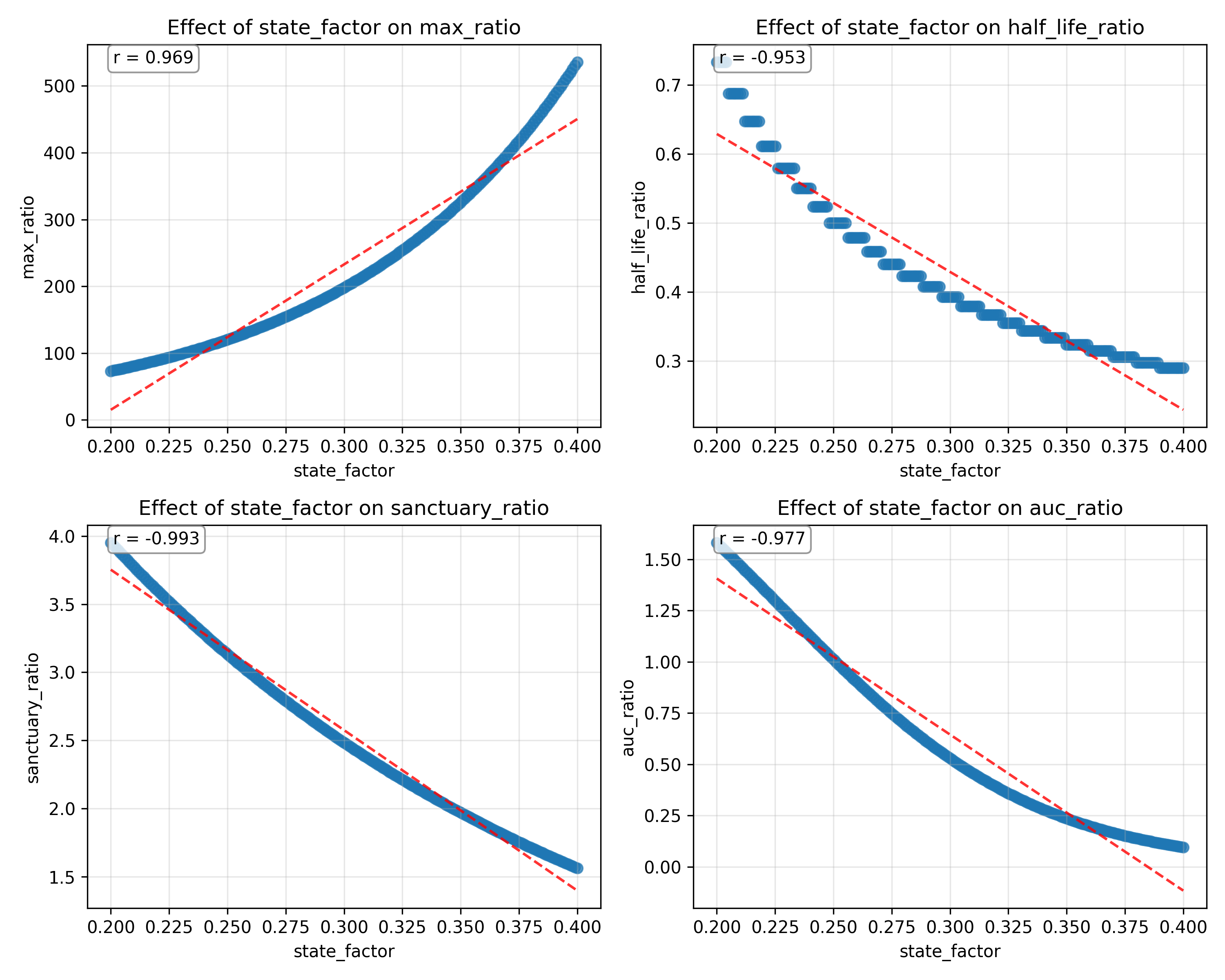

### tergmar_modification.pdf

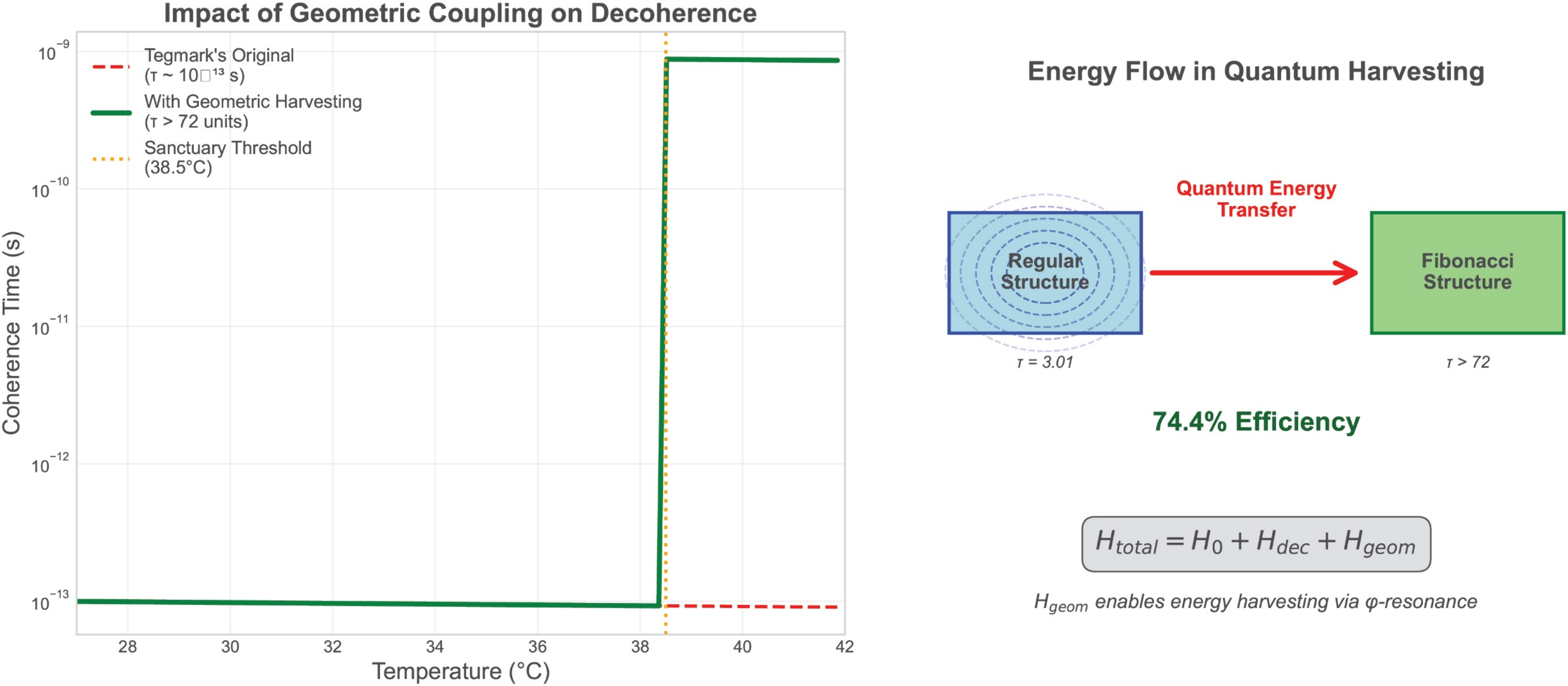
