## Supplementary material for "Quantum Coherence Preservation in Fibonacci-Structured Microtubules During HIV-Induced Neuroinflammation": References

@article{muema2020,
  author = {Muema, D. M. and Akilimali, N. A. and Ndumnego, O. C. and Rasehlo, S. S. and Durgiah, R. and Ojwach, D. B. A. and others},
  title = {Association between the cytokine storm, immune cell dynamics, and viral replicative capacity in hyperacute {HIV} infection},
  journal = {BMC Med},
  year = {2020},
  volume = {18},
  number = {1},
  pages = {81}
}

@article{forthechartergroup2011,
  author = {{for the CHARTER Group} and Moore, D. J. and Letendre, S. L. and Morris, S. and Umlauf, A. and Deutsch, R. and others},
  title = {Neurocognitive functioning in acute or early {HIV} infection},
  journal = {J Neurovirol},
  year = {2011},
  volume = {17},
  number = {1},
  pages = {50--57}
}

@article{hellmuth2019,
  author = {Hellmuth, J. and Slike, B. M. and Sacdalan, C. and Best, J. and Kroon, E. and Phanuphak, N. and others},
  title = {Very Early Initiation of Antiretroviral Therapy During Acute {HIV} Infection Is Associated With Normalized Levels of Immune Activation Markers in Cerebrospinal Fluid but Not in Plasma},
  journal = {J Infect Dis},
  year = {2019},
  volume = {220},
  number = {12},
  pages = {1885--1891}
}

@article{tegmark2000,
  author = {Tegmark, M.},
  title = {Importance of quantum decoherence in brain processes},
  journal = {Phys Rev E},
  year = {2000},
  volume = {61},
  number = {4},
  pages = {4194--4206}
}

@article{engel2007,
  author = {Engel, G. S. and Calhoun, T. R. and Read, E. L. and Ahn, T. K. and Man{\v{c}}al, T. and Cheng, Y. C. and others},
  title = {Evidence for wavelike energy transfer through quantum coherence in photosynthetic systems},
  journal = {Nature},
  year = {2007},
  volume = {446},
  number = {7137},
  pages = {782--786}
}

@article{ritz2002,
  author = {Ritz, T. and Damjanovi{\'c}, A. and Schulten, K.},
  title = {The Quantum Physics of Photosynthesis},
  journal = {ChemPhysChem},
  year = {2002},
  volume = {3},
  number = {3},
  pages = {243}
}

@article{yang2025,
  author = {Yang, I. and Agrenius, T. and Usova, V. and Romero-Isart, O. and Kirchmair, G.},
  title = {Hot {S}chr{\"o}dinger cat states},
  journal = {Sci Adv},
  year = {2025},
  volume = {11},
  number = {14},
  pages = {eadr4492}
}

@article{perez2010,
  author = {P{\'e}rez, J. C.},
  title = {Codon populations in single-stranded whole human genome {DNA} are fractal and fine-tuned by the golden ratio 1.618},
  journal = {Interdiscip Sci},
  year = {2010},
  volume = {2},
  number = {3},
  pages = {228--240}
}

@article{jaeger2021,
  author = {Jaeger, L.},
  title = {The golden ratio in nature: A tour across length scales},
  journal = {Symmetry},
  year = {2021},
  volume = {13},
  number = {11},
  pages = {2019}
}

@article{blokhuis2019,
  author = {Blokhuis, C. and Peeters, C. F. W. and Cohen, S. and Scherpbier, H. J. and Kuijpers, T. W. and Reiss, P. and others},
  title = {Systemic and intrathecal immune activation in association with cerebral and cognitive outcomes in paediatric {HIV}},
  journal = {Sci Rep},
  year = {2019},
  volume = {9},
  number = {1},
  pages = {8004}
}

@article{eckard2017,
  author = {Eckard, A. R. and Rosebush, J. C. and O'Riordan, M. A. and Graves, C. C. and Alexander, A. and Grover, A. K. and others},
  title = {Neurocognitive Dysfunction in {HIV}-Infected Youth: Investigating the Relationship with Immune Activation},
  journal = {Antivir Ther},
  year = {2017},
  volume = {22},
  number = {8},
  pages = {669--680}
}

@article{oh2018,
  author = {Oh, S. W. and Shin, N. Y. and Choi, J. Y. and Lee, S. K. and Bang, M. R.},
  title = {Altered White Matter Integrity in Human Immunodeficiency Virus-Associated Neurocognitive Disorder: A Tract-Based Spatial Statistics Study},
  journal = {Korean J Radiol},
  year = {2018},
  volume = {19},
  number = {3},
  pages = {431}
}

@article{qi2023,
  author = {Qi, Y. and Wang, W. and Rao, B. and Yang, X. and Yu, W. and Li, J. and others},
  title = {Value of Radiomic Analysis Combined With Diffusion Tensor Imaging in Early Diagnosis of {HIV}-Associated Neurocognitive Disorders},
  journal = {J Magn Reson Imaging},
  year = {2023},
  volume = {58},
  number = {6},
  pages = {1882--1891}
}

@article{han2022,
  author = {Han, S. and Aili, X. and Ma, J. and Liu, J. and Wang, W. and Yang, X. and others},
  title = {Altered regional homogeneity and functional connectivity of brain activity in young {HIV}-infected patients with asymptomatic neurocognitive impairment},
  journal = {Front Neurol},
  year = {2022},
  volume = {13},
  pages = {982520}
}

@article{xu2024,
  author = {Xu, F. and Ma, J. and Wang, W. and Li, H.},
  title = {A longitudinal study of the brain structure network changes in {HIV} patients with {ANI}: combined {VBM} with {SCN}},
  journal = {Front Neurol},
  year = {2024},
  volume = {15},
  pages = {1388616}
}

@article{zhou2024,
  author = {Zhou, Z. and Gong, W. and Hu, H. and Wang, F. and Li, H. and Xu, F. and others},
  title = {Large-scale brain network analysis reveals functional-structural dissynchrony in {HIV}-associated asymptomatic neurocognitive disorders: Functional disturbances precede structural changes},
  journal = {Radiology and Imaging},
  year = {2024},
  note = {Available from: \url{http://medrxiv.org/lookup/doi/10.1101/2024.11.17.24317453}}
}
